# Syn-tasiR-VIGS: Virus-Based Targeted RNAi in Plants By Synthetic Trans Acting Small Interfering RNAs Derived from Minimal Precursors

**DOI:** 10.1101/2024.12.18.629176

**Authors:** Adriana E. Cisneros, Ana Alarcia, Juan José Llorens-Gámez, Ana Puertes, María Juárez-Molina, Anamarija Primc, Alberto Carbonell

**Affiliations:** Instituto de Biología Molecular y Celular de Plantas (Consejo Superior de Investigaciones Científicas–Universitat Politècnica de València), 46022 Valencia, Spain

**Keywords:** RNAi, small RNA, syn-tasiRNA, VIGS, antiviral resistance

## Abstract

Synthetic trans-acting small interfering RNAs (syn-tasiRNAs) are 21-nucleotide (nt) small RNAs designed to silence plant transcripts with high specificity. Their use as biotechnological tools for functional genomics and crop improvement is limited by the need to transgenically express long *TAS* precursors to produce syn-tasiRNAs *in vivo*. Here, we show that authentic and highly effective syn-tasiRNAs can be produced from minimal, non-*TAS* precursors consisting of a 22-nt endogenous microRNA target site, an 11-nt spacer and the 21 nt syn-tasiRNA sequence(s). These minimal precursors, when transgenically expressed in *Arabidopsis thaliana* and *Nicotiana benthamiana*, generated highly phased syn-tasiRNAs that silenced one or multiple plant genes with high efficacy. Remarkably, minimal but not full-length *TAS* precursors produced authentic syn-tasiRNAs and induced widespread gene silencing in *N. benthamiana* when expressed from an RNA virus, which can be applied by spraying infectious crude extracts onto leaves in a GMO-free manner. This strategy, named syn-tasiRNA-based virus-induced gene silencing (syn-tasiR-VIGS), was further used to vaccinate plants against a pathogenic virus, resulting in complete plant immunization. Our results reveal that syn-tasiRNA precursors can be significantly shortened without compromising silencing efficacy, and that syn-tasiR-VIGS represents a versatile, scalable and non-transgenic platform for precision RNAi and antiviral vaccination in plants.

**Graphical abstract:** 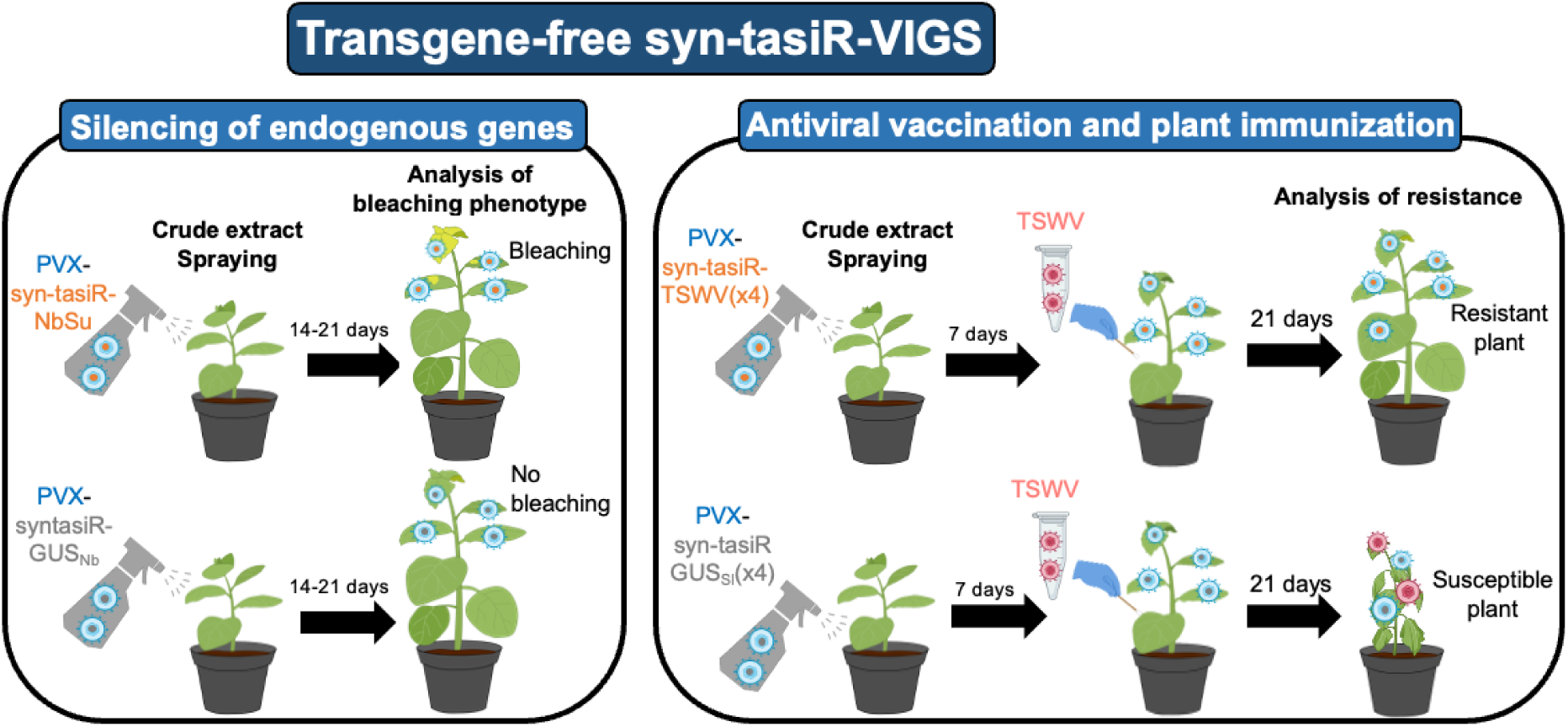

## Introduction

Classic RNA interference (RNAi) strategies for gene silencing rely on the processing of double-stranded RNA (dsRNA) by Dicer ribonucleases into small interfering RNAs (siRNAs) (1, 2). One strand of the duplex, the guide strand, is loaded into an ARGONAUTE (AGO) protein, forming an RNA-induced silencing complex (RISC) that binds and silences target RNAs with high sequence complementarity to the guide strand (2, 3) (Hammond et al., 2000, 2001). Despite their widespread use, classic RNAi strategies present a major limitation, which is the lack of high specificity due to the generation of an uncontrolled array of siRNAs of unpredicted sequence and size that may accidentally bind to cellular transcripts and cause toxic off-target effects (4).

In plants, second generation RNAi tools are based on 21-nucleotide (nt) single-stranded RNA molecules, named artificial sRNAs (art-sRNAs), computationally designed to bind and cleave target RNAs with high specificity and no off-target effects (5). There are two main classes of art-sRNAs: artificial miRNAs (amiRNAs) and synthetic trans-acting siRNAs (syn-tasiRNAs), which are functionally similar but differ in their biogenesis pathway. AmiRNAs are typically produced from *MIR* transgenes, where endogenous miRNA and miRNA* sequences are replaced with amiRNA and amiRNA* sequences. While amiRNAs are often designed to target a single transcript, a unique feature of syn-tasiRNAs is their multiplexing capability, allowing the production of multiple syn-tasiRNAs from a single precursor and, therefore, the multitargeting of different sites within one or multiple target RNAs (6). Syn-tasiRNAs are typically produced from *TAS* transgenes in which the endogenous tasiRNA sequences are replaced by one or more 21-nt syn-tasiRNA sequences in tandem, designed to target the gene(s) of interest (6, 7). These transgenes are transcribed in the nucleus by DNA-DEPENDENT RNA POLYMERASE II into a primary *TAS* transcript (or *TAS* precursor) that includes a 5’ cap structure, a poly-A tail and a target site (TS) specific for a miRNA, usually of 22 nt (8, 9). Once the *TAS* precursor is exported to the cytoplasm, a miRNA/AGO complex mediates the endonucleolytic cleavage of the *TAS* precursor. Next, SUPRESSOR OF GENE SILENCING 3 (SGS3) stabilizes one of the cleaved *TAS* fragments, which is then used by RNA-DEPENDENT RNA POLYMERASE 6 (RDR6) to synthesize a dsRNA molecule. This dsRNA is processed into phased, 21-nt syn-tasiRNA duplexes by DICER LIKE 4 (DCL4), as observed for endogenous tasiRNAs (10, 11). Finally, HUA ENHANCER1 (HEN1) methylates the 3’ end of both strands of the duplex (12), and the guide strand is loaded into AGO1 to cleave or translationally repress complementary RNAs.

*Arabidopsis thaliana* (Arabidopsis) has eight *TAS* loci, belonging to four gene families (*AtTAS1a-c*, *AtTAS2*, *AtTAS3a-c* and *AtTAS4*). Syn-tasiRNAs have been produced in plants from transgenes including modified *AtTAS1a*, *AtTAS1c* and *AtTAS3a* precursors (13–15). *AtTAS1c* is the preferred precursor, as it is accurately processed and yields high levels of authentic syn-tasiRNAs (16, 17) that efficiently silence endogenous genes (18) and confer resistance to plant viruses and viroids (19, 20). Importantly, *AtTAS1c*-derived syn-tasiRNA biogenesis is triggered by the Arabidopsis 22-nt miRNA miR173a, which is only present in this species and its close relatives. Therefore, in non-Arabidopsis species, the *AtMIR173a* gene must be co-expressed with the *AtTAS1c* precursor to produce syn-tasiRNAs.

Syn-tasiRNAs possess unique features that have boosted their application for functional genomics and crop improvement (7, 21): (i) their multiplexing capability enables the simultaneous production of multiple syn-tasiRNAs from a single precursor for multitargeting, (ii) their expression at different precursor positions allows for fine-tuning targeted RNAi efficacy (18), and (iii) the development of fast-forward methodologies for their design and cloning accelerates the generation of syn-tasiRNA constructs (16, 22–24). Still, a major limitation of syn-tasiRNA technology is the need for integrating *TAS*-based transgenes into plant genomes, a process that is time-consuming and raises regulatory concerns about the generation of genetically-modified plants (25). Also, another limitation is the use of relatively long *TAS*-based precursors, which i) may increase the cost of the *in vitro* or bacterial synthesis of RNA precursors for topical delivery, and ii) may not be stable when incorporated into viral vectors, as reported recently for full-length amiRNA precursors (26). Therefore, further optimization of the syn-tasiRNA technology is required to expand its applicability while avoiding the generation of transgenic plants.

Here, we show that authentic and highly effective syn-tasiRNAs can be produced from minimal, non-*TAS* precursors consisting of a 22-nt endogenous microRNA target site, an 11-nt spacer and the 21 nt syn-tasiRNA sequence(s). These minimal precursors were functional when expressed in transgenic plants or from an RNA virus, allowing for multi-gene silencing and for plant antiviral vaccination in a DNA-free manner.

## Materials and methods

### Plant species and growth conditions

*N. benthamiana* plants were grown in a growth chamber at 25°C with a 12 h-light/12 h dark photoperiod. Arabidopsis Col-0 plants were grown in a growth chamber at 22°C with a 16 h-light/8 h-dark photoperiod. Genetic transformation of Arabidopsis was done following the floral dip method (27) using the *Agrobacterium tumefaciens* GV3101 strain. T1 transgenic Arabidopsis were done as described (18). A Nikon D3000 digital camera with AF-S DX NIKKOR 18-55 mm f/3.5–5.6G VR lens was used for photographing plants.

### Arabidopsis phenotyping

Arabidopsis phenotyping analyses were performed in blind as described (18). Briefly, the flowering time of each independent line results from the number of days elapsed since seed plating to first bud opening (or ‘days to flowering’). The ‘Ft’ phenotype was defined as a higher ‘days to flowering’ value when compared to the average ‘days to flowering’ value of the *35S:AtTAS1c-GUS_At_* control set. The ‘CH42’ phenotype was scored in 10 day-old seedlings and was considered ‘weak’, ‘intermediate’ or ‘severe’ if seedlings had more than two leaves, exactly two leaves or no leaves at all (only two cotyledons), respectively. A line was considered to have a ‘Trich’ phenotype when presenting a visually obvious higher number of trichomes in rosette leaves of 14-day-old seedlings when compared to transformants of the *35S:AtTAS1c-GUS_At_*control set.

### Artificial small RNA design

Syn-tasiR-GUS_At_, syn-tasiR-AtFT, syn-tasiR-AtCH42, syn-tasiR-AtTRY, syn-tasiR-GUS_Nb_, syn-tasiR-NbSu, syn-tasiR-GUS_Nb_-1/amiR-GUS_Nb_, syn-tasiR-GUS_Nb_-2, syn-tasiR-TSWV-1/amiR-TSWV, syn-tasiR-TSWV-2, syn-tasiR-TSWV-3 and syn-tasiR-TSWV-4 guide sequences were described before (17–20, 28).

### DNA constructs

Oligonucleotides AC-674 and AC-675 were annealed and ligated into *pENTR-D-TOPO* (Invitrogen) following manufacturer’s instructions to generate *pENTR-BB* including two inverted *Bsa*I restriction sites. The *B/c* cassette was excised from *Bsa*I-digested *pENTR-AtMIR390a-B/c* (Addgene plasmid #51778) (16) and ligated into *Bsa*I-digested *pENTR-BB* to generate *pENTR-B/c*. The *BB* cassette from *pENTR-BB* was transferred by LR recombination into *pMDC32B* (16) to generate *pMDC32B-BB*. Finally, the *B/c* cassette was excised from *Bsa*I-digested *pENTR-B/c* and ligated into *Bsa*I-digested *pMDC32B-BB* to generate *pMDC32B-B/c*. Oligonucleotides AC-1093/AC-1094 and AC-900/AC-901 including minimal *AtmiR173aTS* and *NbmiR482aTS*-based precursors, respectively, were annealed and ligated into *Bsa*I-digested *pENTR-B/c* and *pMDC32B-B/c* to generate *pENTR-AtmiR173aTS-BB pENTR-NbmiR482aTS-BB*, *pMDC32B-AtmiR173aTS-BB* and *pMDC32B-NbmiR482aTS-BB*, respectively. The B/c cassette was excised from *Bsa*I-digested *pENTR-AtTAS1c-D2-B/c* (Addgene plasmid #137883) (18) and ligated into *Bsa*I-digested *pENTR-AtmiR173aTS-BB, pENTR-NbmiR482aTS-BB*, *pMDC32B-AtmiR173aTS-BB* and *pMDC32B-NbmiR482aTS-BB* to generate *pENTR-AtmiR173aTS-B/c pENTR-NbmiR482aTS-B/c*, *pMDC32B-AtmiR173aTS-B/c* and *pMDC32B-NbmiR482aTS-B/c*. New B/c vectors are available from Addgene: *pENTR-B/c* (Addgene plasmid #227962), *pMDC32B-B/c* (Addgene plasmid #227963), *pENTR-AtmiR173aTS-B/c* (Addgene plasmid #227964), *pMDC32B-AtmiR173aTS-B/c* (Addgene plasmid #227965), *pENTR-NbmiR482aTS-B/c* (Addgene plasmid #227966), *pMDC32B-NbmiR482aTS-B/c* (Addgene plasmid #227967).

Oligonucleotide pairs AC-676/AC-677, AC-678/AC-679, AC-680/AC-681, AC-682/AC-683, AC-684/AC-685, AC-639/AC-640, AC-643/AC-644, AC-641/AC-642 and AC-645/AC-646 were annealed and ligated into *pENTR-D-TOPO* (Invitrogen) following manufacturer’s instructions to generate *pENTR-AtmiR173aTS-GUS_At_*, *pENTR-AtmiR173aTS-AtCH42*, *pENTR-AtmiR173aTS-AtFT*, *pENTR-AtmiR173aTS-AtFT-AtTrich*, *pENTR-AtmiR173aTS-AtTrich-AtFT*, *pENTR-NbmiR482aTS-GUS_Nb_*, *pENTR-NbmiR6019a/bTS-GUS_Nb_*, *pENTR-NbmiR482aTS-NbSu* and *pENTR-NbmiR6019a/bTS-NbSu*, respectively. Minimal precursor cassettes were transferred by LR recombination into *pMDC32* (29) to generate *35S:AtmiR173aTS-GUS_At_*, *35S:AtmiR173aTS-AtCH42*, *35S:AtmiR173aTS-AtFT*, *35S:AtmiR173aTS-AtFT-AtTrich*, *35S:AtmiR173aTS-AtTrich-AtFT*, *35S:NbmiR482aTS-GUS_Nb_*, *35S:NbmiR6019a/bTS-GUS_Nb_*, *35S:NbmiR482aTS-NbSu* and *35S:NbmiR6019a/bTS-NbSu*.

Target site swaps were done by mutagenic PCR using *pENTR:AtTAS1c-GUS_Nb_*(17) as template and oligo pairs AC-631/AC-505 and AC-632/AC-507 to generate *pENTR-AtTAS1c(NbmiR482aTS)-NbSu* and *pENTR-AtTAS1c(NbmiR6019a/bTS)-NbSu*. *AtTAS1c*-based cassettes were transferred by LR recombination into *pMDC32* to generate *35S:AtTAS1c(NbmiR482aTS)-NbSu* and *35S:AtTAS1c(NbmiR6019a/bTS)-NbSu*.

Constructs *35S:AtmiR156aTS-NbSu*/*35S:AtmiR173aTS-NbSu* and *35S:NbmiR482aTS-TSWV(x4* were obtained by ligating annealed oligonucleotide pairs AC-719/AC-720–AC-721/AC-722 and AC-935/AC-936 into *pMDC32B-B/c* and *pMDC32B-NbmiR482aTS-B/c*, respectively, as described (16) (Figure S1 and S2). A detailed protocol for cloning syn-tasiRNA minimal precursors in B/c vectors is described in Text S1.

For PVX-based amiRNA constructs, amiRNA cassettes *amiR-GUS_Nb_*and *amiR-TSWV* were amplified from *35S:AtMIR390a-GUS_Nb_* and *35S:AtMIR390a-TSWV-L-5* with oligonucleotide pair AC-650/AC-663 (20) and gel purified. For PVX-based syn-tasiRNA constructs, syn-tasiRNA cassettes *AtTAS1c(NbmiR482aTS)-NbSu* and *NbmiR482aTS-TSWV(x4)* were amplified from *35S:AtTAS1c(NbmiR482aTS)-NbSu* and *35S:NbmiR482aTS-TSWV(x4)* with oligonucleotide pairs AC-712/AC-713 and AC-988/AC-989 and gel purified. Syn-tasiRNA cassettes *NbmiR482aTS-NbSu* and *NbmiR482aTS-GUSNb(x4)* were ordered as dsDNA oligonucleotides AC-666 and AC-987. All amiRNA and syn-tasiRNA cassettes were assembled into *Mlu*I-digested and gel-purified *pLBPVXBa-M* (Addgene plasmid #229079) (30) in the presence of GeneArt Gibson Assembly HiFi Master Mix (Invitrogen) to generate *35S:PVX-amiR-GUS_Nb_*, *35S:PVX-amiR-TSWV*, *35S:PVX-NbmiR482aTS-NbSu*, *35S:PVX-NbmiR482aTS-GUS_Nb_(x4)* and *35S:PVX-NbmiR482aTS-TSWV(x4)*.A detailed protocol for cloning syn-tasiRNA minimal precursors into *pLBPVXBa-M* is described in Text S2.

Syn-tasiRNA constructs *35S:AtTAS1c-GUS_At_*, 35S*:AtTAS1c-AtFT, 35S:AtTAS1c-AtCH42, 35S:AtTAS1c-AtFT-AtTrich, 35S:AtTAS1c-AtTrich-AtFT, 35S:AtTAS1c-GUS_Nb_/AtMIR173*, *35S:AtTAS1c-NbSu/AtMIR173* and *35S:PVX* constructs were described before (16, 18, 26). All DNA oligonucleotides used in this study are listed in Table S1. The sequences of all syn-tasiRNA precursors are listed in Text S1. The sequences of newly developed B/c vectors are listed in Text S2.

### Transient expression of constructs and spray-based inoculation of viruses

*Agrobacterium*-mediated infiltration of DNA constructs in *N. benthamiana* leaves was done as previously (8, 31). Preparation and spraying of crude extracts obtained from virus infected *N. benthamiana* plants was done as previously (26) including 5% silicon carbide (carborundum) in the inoculation buffer.

### Chlorophyll extraction and analysis

Chlorophyll and other pigments from *N. bentamiana* leaves were extracted and analyzed as described (18, 32).

### RNA preparation

Total RNA form *N. benthamiana* leaves or from Arabidopsis seedlings or inflorescences was isolated as before (26). Triplicate samples from pools of two *N. benthamiana* leaves or 9-12 Arabidopsis seedlings or inflorescences were analyzed.

### Real-time RT-qPCR

cDNA was obtained from 500 ng of DNAseI-treated total RNA from 10 day-old Arabidopsis seedlings, 60 day-old Arabidopsis plants, or from *N. benthamiana* leaves at 2 days post-agroinfiltration (dpa) using the PrimeScript RT Reagent Kit (Perfect Real Time, Takara) according to manufacturer’s instructions. Real time RT-qPCR was done using the same RNA samples that were used for sRNA-blot analysis. RT-qPCR was done on optical 96-well plates in a QuantStudio 3 Real-Time PCR system (Thermo Fisher Scientific, Waltham, MA, USA) using the following program: 20 s at 95°C, followed by 40 cycles of 95°C for 3 s and 60°C for 30 s, with an additional melt curve stage consisting of 15 s at 95°C, 1 min at 60°C and 15 s at 95°C. The 20-ml reaction mixture contained 10 ml of 2× TB Green Premix Ex Taq (Takara), 2 ml of diluted complementary DNA (1:5), 0.4 ml of ROX II Reference Dye (50X) and 300 nM of each gene-specific primer. Oligonucleotides used for RT-qPCR are listed in Table S1. Target mRNA expression levels were calculated relative to reference genes *AtACT2* and *NbPP2A* in Arabidopsis and *N. benthamiana*, respectively, using the delta delta cycle threshold comparative method of QuantStudio Design and Analysis software, version 1.5.1 (Thermo Fisher Scientific). Three independent biological replicates, and two technical replicates for each biological replicate were analyzed.

### Stability and sequence analyses of syn-tasiRNA precursors during viral infections

Total RNA from apical leaves of each of the three biological replicates was pooled before cDNA synthesis. PCR to detect syn-tasiRNA precursors, PVX and *NPP2A* was performed using oligonucleotide pairs AC-654/AC-655, AC-657/AC-658 and AC365/AC-366 (Table S1), respectively, and Phusion DNA polymerase (ThermoFisher Scientific). PCR products were analyzed by agarose gel electrophoresis, and products of the expected sized were excised from the gel and sequenced when necessary.

### Small RNA blot assays

Small RNA blot assays and band quantification from radioactive membranes were done as described (17). Oligonucleotides used as probes for sRNA blots are listed in Table S1.

### Small RNA sequencing and data analysis

The quantity, purity and integrity of total RNA was analyzed with a 2100 Bioanalyzer (RNA 6000 Nano kit, Agilent) and submitted to BGI (Hong Kong, China) for sRNA library construction and SE50 high-throughput sequencing in a DNBSEQ-G-400 sequencer. Quality-trimmed, adaptor removed clean reads received from BGI were used with the fastx_collapser toolkit (http:// hannonlab.cshl.edu/fastx toolkit) (33) to collapse identical reads into a single sequence, while maintaining read counts. Mapping of each clean, unique read against the forward strand of the syn-tasiRNA precursor expressed in each sample (Data S1) was done with a custom Python script not allowing mismatches or gaps, and also to calculate the counts and RPMs (reads per million mapped reads) for each mapping position. Processing accuracy of syn-tasiRNA precursors was assessed by quantifying the proportion of 19–24 nt sRNA (+) reads that mapped within ± 4 nt of the 5′ end of the syn-tasiRNA guide as reported before (8, 32). Phasing register tables were built by calculating the proportion of 21-nucleotide sRNA (+) reads in each register relative to the corresponding amiRNA cleavage site for all 21-nucleotide positions downstream of the cleavage site, as described previously (16).

### Protein blot analysis

Proteins were separated in NuPAGE Novex 4-12% Bis-Tris gels (Invitrogen), transferred to Protran nitrocellulose membranes (Amersham) and detected by chemiluminescence using specific antibodies and SuperSignal West Pico PLUS chemiluminescent substrate (Thermo Fisher Scientific). For detection of TSWV, anti-TSWV nucleocapsid (N) (Bioreba) was used at 1:10000 dilution and conjugated with 1:20000 of goat anti-rabbit IgG horseradish peroxidase secondary antibody (Thermo Fisher Scientific). Images were acquired with an ImageQuant 800 CCD imager (Cytiva) and analyzed with ImageQuantTL v10.2 (Cytiva). Ponceau red S solution (Thermo Fisher Scientific) staining of membranes was used to verify the global protein content of the samples.

### Statistical analysis

Statistical tests are described in the figure legends. Significant differences were determined with two-tailed Student’s *t-*test.

### Gene and virus identifiers

Arabidopsis and *N. benthamiana* gene identifiers are *AtACT2* (AT3G18780), *AtCH42* (AT4G18480), *AtFT* (AT1G65480), *AtTRY* (AT5G53200), *NbSu* (Nbv5.1tr6204879) and *NbPP2A* (Nbv5.1tr6224808), TSWV LL-N.05 segment L, M and S genome identifiers are KP008128, FM163373 and KP008129, respectively. PVX-based constructs include PVX sequence variant MT799816.1. *Escherichia coli* b-glucuronidase gene sequence corresponds to GenBank accession number S69414.1.

## Results

### Effective gene silencing in *Arabidopsis thaliana* by syn-tasiRNAs derived from minimal precursors

Previous work has shown that secondary siRNAs can be generated from non-*TAS* constructs including gene fragments fused to a 22-nt miRNA target site (TS) (14, 34). Thus, we hypothesized that syn-tasiRNAs could similarly be produced from minimal precursors consisting exclusively of an endogenous 22-nt miRNA target site, followed by a 11-nt spacer from *AtTAS1c* 3’D1[+] fused to the syn-tasiRNA sequence. To test this, we used the syn-tasiR-AtFT/*AtFT* and syn-tasiR-AtCH42/*AtCH42* silencing sensor systems in Arabidopsis where transgenic expression of syn-tasiR-AtFT or syn-tasiR-AtCH42 from full-length *AtTAS1c* precursors at DCL4-processing position 3’D2[+] induces a significant delay in flowering time or an intense bleaching due to the targeting of *FLOWERING LOCUS T* (*AtFT*) or *CHLORINA 42* (*AtCH42*), respectively (18). Here, we generated the *35S:AtmiR173aTS-AtFT* and *35S:AtmiR173aTS-AtCH42* constructs for expressing syn-tasiR-AtFT and syn-tasiR-AtCH42, respectively, from minimal precursors including exclusively the 22-nt (TS) sequence (GTGATTTTTCTCTACAAGCGAA) of Arabidopsis miR173a (AtmiR173a) followed by the 11-nt spacer sequence (TAGACCATTTA) from *AtTAS1c* 3’D1[+] (Figure 1A). In addition, the *35S:syn-tasiR-GUS_At_* construct was also generated for expressing a syn-tasiRNA against *Escherichia coli uidA* b-glucuronidase gene (or *GUS*) with no predicted off-targets in Arabidopsis (Figure 1A). These constructs were independently transformed into Arabidopsis Col-0 plants together with control constructs *35S:AtTAS1c-GUS_At_*, *35S:AtTAS1c-AtFT* and *35S:AtTAS1c-AtCH42* expressing a syn-tasiRNA against *GUS*, *AtFT* and *AtCH42*, respectively, from position 3’D2[+] in *AtTAS1c* (18). To compare the activity of syn-tasiR-AtFT and syn-tasiR-AtCH42 produced from minimal and *AtTAS1c* precursors, plant phenotypes, syn-tasiRNA and target mRNA accumulation as well as precursor processing efficiency were measured in Arabidopsis T1 transgenic lines.

**Figure 1.**
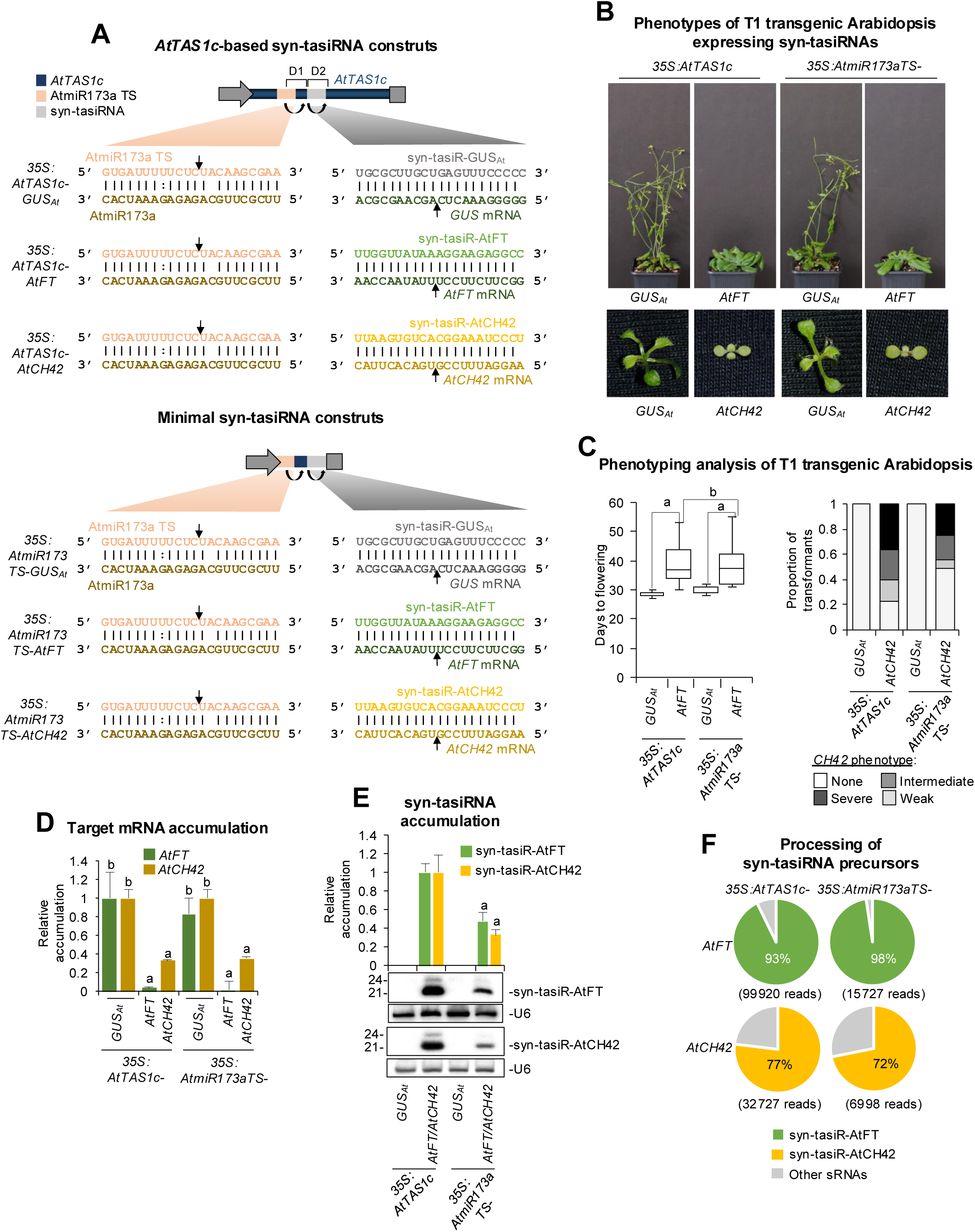
Functional analysis of *AtTAS1c* and minimal precursors expressing a single syn-tasiRNA against Arabidopsis *FT* (*AtFT*) or *CH42* (*AtCH42*) in T1 transgenic plants. (**A**) Organization of syn-tasiRNA constructs. Diagram of *AtTAS1c* (up) and minimal (down) precursors including the AtmiR173a/AtmiR173a target site (TS) and syn-tasiRNA/target mRNA base-paring interactions. Nucleotides corresponding to AtmiR173a and its target site (AtmiR173aTS) are in dark and light orange, respectively. Nucleotides corresponding to syn-tasiR-GUS_At_, syn-tasiR-AtFT and syn-tasiR-AtCH42 are shown in light grey, green and yellow, respectively, while nucleotides of their target mRNAs are shown in dark grey, green or yellow, respectively. Arrows indicate the predicted cleavage sites for AtmiR173a and syn-tasiRNAs. (**B**) Representative photographs of Arabidopsis plants expressing syn-tasiRNAs from *AtTAS1c* or minimal precursors. Upper panel: 45-day-old plants expressing syn-tasiR-GUS_At_ or syn-tasiR-AtFT. Lower pannel: 10-day-old seedlings expressing syn-tasiR-GUS_At_ or syn-tasiR-AtCH42. (**C**) Phenotypic analysis of plants expressing syn-tasiRNAs from *AtTAS1c* or minimal precursors. Left: mean flowering time of plants expressing syn-tasiR-GUS_At_ or syn-tasiR-AtFT. Pairwise Student’s t-test comparisons are represented with black lines including the letter ‘a’ if significantly different (*P* < 0.05) and the letter ‘b’ if not (*P* > 0.05). Right: bar graphs representing, for each line, the proportion of seedlings displaying a severe (black areas), intermediate (dark grey areas), or weak (light grey areas) bleaching phenotype, or with wild-type appearance (white areas). (**D**) Target *AtFT* and *AtCH42* mRNA accumulation in RNA preparations from Arabidopsis plants [mean relative level (n = 3) + standard error] after normalization to *ACTIN* 2 (*AtACT2*), as determined by quantitative RT-qPCR (*35S:AtTAS1c-GUS_At_* = 1). Bars with the letter ‘a’ or ‘b’ are significantly different (*P* < 0.05) or not (*P* > 0.05) from the corresponding *35S:AtTAS1c-GUS_At_* control samples, respectively, based on pairwise Student’s t-test comparisons. (**E**) Northern blot detection of syn-tasiR-AtFT and syn-tasiR-AtCH42 in RNA preparations from Arabidopsis plants. The graph at the top shows the mean + standard deviation (n = 3) syn-tasiRNA relative accumulation (*35S:AtTAS1c-AtFT* = 1.0 and *35S:AtTAS1c-AtCH42* = 1.0). Bars with the letter ‘a’ are significantly different from that of *35S:AtTAS1c-AtFT* or *35S:AtTAS1c-AtCH42* control samples. One blot from three biological replicates is shown. Each biological replicate is a pool of at least nine independent lines selected randomly. U6 RNA blots are shown as loading controls. (**F**) Syn-tasiRNA processing from *AtTAS1c* or minimal precursors. Pie charts show the percentages of reads corresponding to the expected, accurately processed 21-nt mature syn-tasiR-AtFT and syn-tasiR-AtCH42 (green and yellow sections, respectively) or to other 19–24-nt sRNAs (grey sectors).

All *35S:AtTAS1c-AtFT* (n = 65) and *35S:AtmiR173aTS-AtFT* (n = 53) transformants flowered later than the average flowering time of the *35S:AtTAS1c-GUS_At_*and *35S:AtmiR173aTS-GUS_At_* control lines (n = 58 and n = 64, respectively) (Figure 1B and 1C, Table S2). The average flowering time (39.3 ± 6.6 and 38.9 ± 6.6, respectively) was not significantly different between the two (Figure 1C). RT-qPCR assays revealed that *AtFT* mRNA accumulation was similar in lines expressing syn-tasiR-AtFT from each of the two precursors (Figure 1D), while RNA-blots showed that syn-tasiR-AtFT accumulated to significantly higher levels when expressed from full-length *AtTAS1c* precursors (Figure 1E). Finally, high-throughput sequencing of sRNAs showed that *AtTAS1c* and minimal precursors were efficiently processed, with 93% and 98% of the reads corresponded to authentic syn-tasiR-AtFT, respectively (Figure 1F). Similarly, all *35S:AtTAS1c-AtCH42* (n = 402) and *35S:AtmiR173aTS-AtCH42* (n = 389) transformants displayed bleaching to comparable degrees (Figure 1B and C; Table S3), accumulated similar levels of *AtCH42* mRNA (Figure 1D) and displayed effective precursor processing (Figure 1E and 1F), though reduced amounts of syn-tasiR-AtCH42 were observed in *35S:AtmiR173TS-AtCH42* transformants (Figure 1E).

Next, we analyzed whether minimal precursors could be used to express multiple, accurately processed and phased syn-tasiRNAs for the simultaneous silencing of different endogenous genes. For this purpose, we used the syn-tasiR-AtFT/syn-tasiR-AtTRY silencing sensor system in Arabidopsis, in which plants co-expressing both syn-tasiRNAs exhibit non-overlapping silencing phenotypes of delay in flowering time and an increase in trichome number, due to the silencing of *AtFT* and of three *MYB* transcripts [*TRIPTYCHON (AtTRY)*, *CAPRICE* (*AtCPC*) and *ENHANCER OF TRIPTYCHON AND CAPRICE2 (AtETC2)*] (16) (Figure 2A). Here, we compared the silencing efficacy of syn-tasiR-AtFT and syn-tasiR-AtTRY when co-expressed in unique dual configurations from positions 3’D2[+] and 3’D3[+] in full-length *AtTAS1c* (constructs *35S:AtTAS1c-AtFT-AtTRY* and *35S:AtTAS1c-AtTRY-AtFT*) or minimal precursors (constructs *35S:AtmiR173aTS-AtFT-AtTRY* and *35S:AtmiR173aTS-AtTRY-AtF*T) (Figure 2B). We also included in the analysis control constructs *35S:AtTAS1c-GUS_At_*and *35S:AtmiR173aTS-GUS_At_* for expressing syn-tasiR-GUS_At_ in single configuration from position 3’D2[+] in full-length *AtTAS1c* or minimal precursors, respectively. All these constructs were transformed into Arabidopsis Col-0 plants, and T1 transformants were analyzed phenotypically by scoring the day of flowering and the trichome number in rosette leaves for each transformant, and molecularly by quantifying target mRNA and syn-tasiRNA accumulation in the different lines by RT-qPCR and RNA blot assays, respectively.

**Figure 2.**
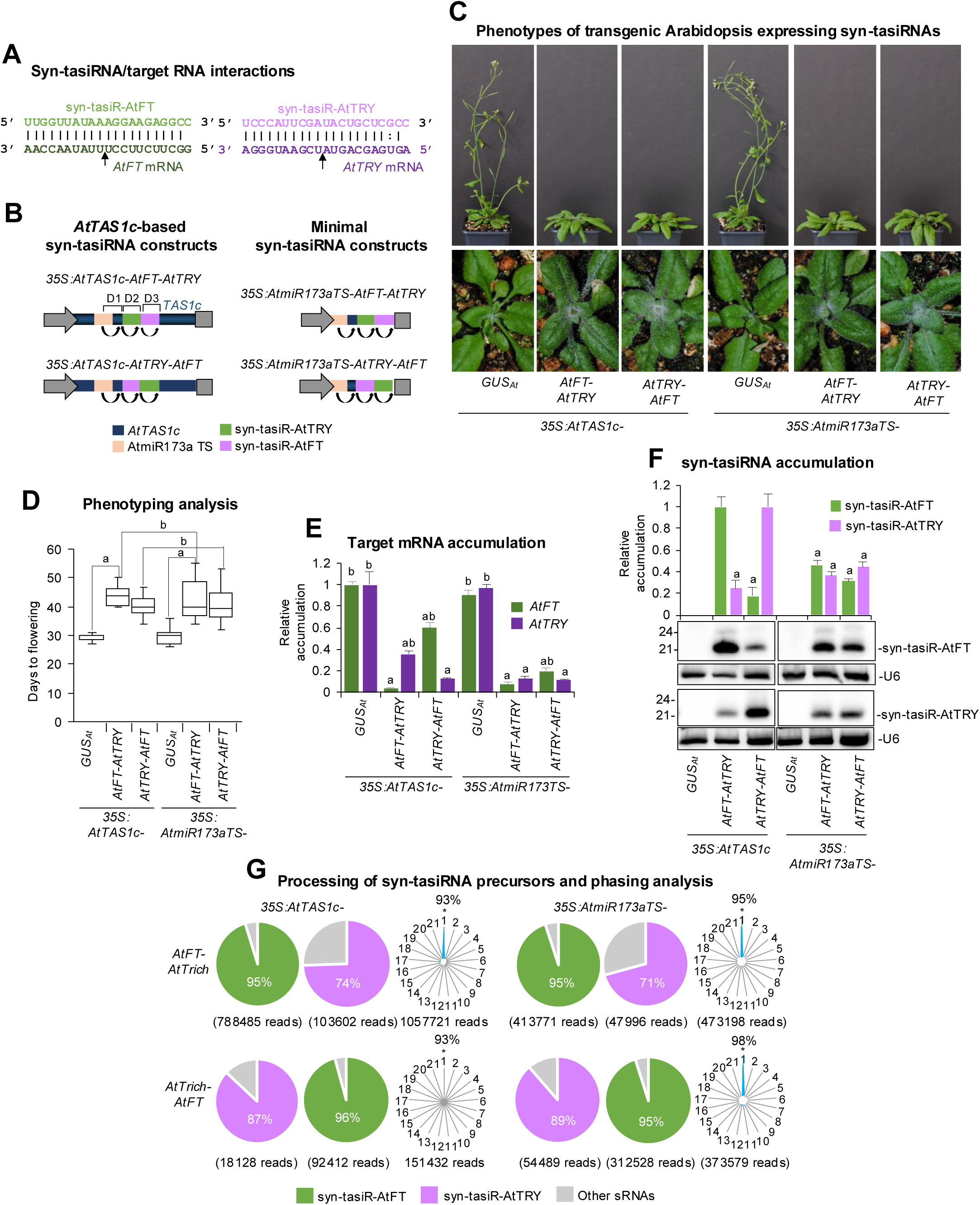
Functional analysis of *AtTAS1c* and minimal precursors expressing syn-tasiRNAs from distinct dual configurations in Arabidopsis T1 transgenic plants. (**A**) Base-pairing of syn-tasiRNAs and target mRNAs. Nucleotides corresponding to syn-tasiR-AtFT and *AtFT* mRNA are shown in light and dark green, respectively. Nucleotides corresponding to syn-tasiR-AtTRY and *AtTRY* mRNA are shown in light and dark purple, respectively. Arrows indicate the syn-tasiRNA cleavage sites. (**B**) Diagram of *AtTAS1c* (left) and minimal (right) precursor constructs. The AtmiR173a target site (TS), syn-tasiR-AtFT and syn-tasiR-AtTRY are represented in orange, green and purple, respectively. Other details as in Figure 1A. (**C**) Representative photographs of Arabidopsis plants expressing two syn-tasiRNAs in tandem from *AtTAS1c* or minimal precursors. The same plant was photographed at 20 days post-plating (dpp) to visualize the increased number of trichomes (bottom) and at 45 dpp to confirm the delay in flowering (top). (**D**) Phenotypic analysis of plants expressing syn-tasiRNAs from *AtTAS1c* or minimal precursors. Mean flowering time of plants expressing syn-tasiR-AtFT and syn-tasiR-AtTRY is represented. Other details as in Figure 1C. (**E**) Target *AtFT* and *AtTRY* mRNA accumulation [mean relative level (n = 3) + standard error] after normalization to *ACTIN* 2 (*AtACT2*), as determined by quantitative RT-qPCR, (*35S:AtTAS1c-GUS_At_* = 1). Other details as in Figure 1D. (**F**) Northern blot detection of syn-tasiR-AtFT and syn-tasiR-AtTrich in RNA preparations from Arabidopsis plants. The graph at the top shows the mean + standard deviation (n = 3) syn-tasiRNA relative accumulation (*35S:AtTAS1c-AtFT-AtTRY* and *35S:AtTAS1c-AtTRY-AtFT* = 1.0). Other details as in Figure 1E. (**G**) Syn-tasiRNA processing and phasing analysis from *AtTAS1c*- and minimal precursors. Pie charts show the percentage of reads corresponding to expected, accurately processed 21-nt mature syn-tasiR-AtFT or syn-tasiR-AtTrich (green or purple sections, respectively) or to other 19-24 nt sRNAs (grey sectors). Radar plots show the proportion of 21-nt reads corresponding to each of the 21 registers from *AtTAS1c* transcripts, with position 1 designated as immediately after the AtmiR173a guided cleavage site. The percentage of 21-nt reads corresponding to phasing register 1 is indicated.

Regarding flowering time analysis, all transformants expressing the dual-configuration syn-tasiRNA constructs showed a delay in flowering compared to *35S:AtTAS1c-GUS_At_* or *35S:AtmiR173aTS-GUS_At_*control transformants (Figure 2C, Table S4). In particular, transformants expressing syn-tasiR-AtFT from minimal precursors at positions 3’D2[+] or 3’D3[+] had a mean flowering time of 43 ± 7.3 or 40.5 ± 5.4 days, respectively, which was similar to that of transformants expressing the same syn-tasiRNA from the same positions (43 ± 5.4 and 40.2 ± 3.9 days, respectively) (Figure 2D). Regarding the trichome number analysis, 80% and 72% of transformants expressing syn-tasiR-AtTRY from minimal precursors at positions 3’D2[+] or 3’D3[+], respectively, had higher number of trichomes compared to the *35S:AtmiR173aTS-GUS_At_* control group, similarly to the transformants expressing the same syn-tasiRNA from the same positions (88% and 82%, respectively) in full-length precursors (Table S4). Interestingly, target mRNA accumulation analysis by RT-qPCR showed a similar and drastic decrease of *AtFT* and *AtTRY* mRNA levels in dual-configuration transformants compared to *35S:AtTAS1c-GUS_At_* and *35S:AtmiR173aTS-GUS_At_* controls, regardless of the size of precursor used (Figure 2E). RNA blot assays confirmed that both minimal and full-length precursors produced detectable levels of syn-tasiRNAs that accumulated as single 21-nt bands (Figure 2F), suggesting an accurate processing from both types of precursors. To further confirm the accuracy of precursor processing, sRNAs from the four dual-configuration transformants were sequenced and analyzed. Both classes of precursors were efficiently processed, with 95-96% and 71-89% of the reads corresponding to authentic syn-tasiR-AtFT or syn-tasiR-AtTRY, respectively (Figure 2G). Moreover, the tasiRNA pools triggered by AtmiR173a were highly phased, with 93-98% of 21-nt reads corresponding to the first register (Figure 2G).

Overall, these results indicate that minimal syn-tasiRNA precursors, including the 22-nt AtmiR173a TS and the 11-nt *AtTAS1c* spacer, produce authentic 21-nt phased syn-tasiRNA species when stably expressed in Arabidopsis. These syn-tasiRNAs induce highly effective silencing of endogenous genes, similarly to those expressed from full-length *AtTAS1c* precursors.

### Effective gene silencing in *Nicotiana benthamiana* by syn-tasiRNAs derived from minimal precursors

Next, we aimed to confirm in another plant species such as *Nicotiana benthamiana* that accurately processed and phased syn-tasiRNAs could be produced from minimal precursors. Since AtmiR173a is exclusively present in Arabidopsis and its close relatives, we hypothesized that syn-tasiRNAs could be generated in *N. benthamiana* from minimal precursors including a TS from an endogenous 22-nt miRNA such as NbmiR482a or NbmiR6019a/b, instead of the original AtmiR173a TS.

First, we examined whether syn-tasiRNA biogenesis could be triggered from the *35S:AtTAS1c-(NbmiR482aTS)-NbSu* or *35S:AtTAS1c-(NbmiR6019a/bTS)-NbSu* constructs engineered to produce syn-tasiR-NbSu, a syn-tasiRNA silencing the *N. benthamiana* magnesium chelatase subunit CHLI-encoding *SULPHUR* (*NbSu*) (17), from modified *AtTAS1c* precursors including NbmiR482a or NbmiR6019a/b TSs, respectively (Figure 3A). Both constructs were independently agroinfiltrated in two areas of two leaves from three different plants, together with control constructs *35S:AtTAS1c-NbSu/AtMIR173a* and *35S:AtTAS1c-GUS_Nb_/AtMIR173a* (18) for producing syn-tasiR-NbSu and syn-tasiR-GUS_Nb_, respectively, due to the co-expression of AtmiR173a (Figure 3A). At 7 dpa, all areas agroinfiltrated with anti-NbSu syn-tasiRNA constructs displayed a strong bleaching, as expected from *NbSu* knockdown, while areas expressing the control syn-tasiRNA did not (Figure 3A). These bleached areas accumulated significantly reduced amounts of chlorophyll *a* compared to areas expressing the control syn-tasiRNA (Figure 3B). Next, two leaves of three different plants were independently agroinfiltrated over the entire leaf surface with each of the syn-tasiRNA constructs described above. RT-qPCR and RNA blot assays of RNA preparations obtained at 2 dpa from agroinfiltrated leaves showed that, in all samples expressing canonical or modified *AtTAS1c* precursors, *NbSu* mRNA accumulation was drastically reduced and syn-tasiR-NbSu accumulated as a single 21-nt band, although samples expressing *AtTAS1c(NbmiR6019a/bTS)* accumulated lower levels of syn-tasiR-NbSu than the others (Figure 3C and 3D). Importantly, high-throughput sequencing analysis of sRNAs showed that *AtTAS1c(NbmiR482aTS)* and canonical *AtTAS1c* precursors were processed with similar accuracy, with 59% and 58% of reads within ± 4 nt of 3’D2[+] corresponded to authentic syn-tasiR-NbSu, respectively, while this percentage was lower (37%) in samples expressing *AtTAS1c(NbmiR6019a/bTS)* precursors (Figure 3E). Moreover, highly phased siRNAs were generated from canonical *AtTAS1c* or modified *AtTAS1c(NbmiR482aTS)* precursors, with 63% and 54% of 21-nt [+] reads, respectively, corresponding to the first register (Figure 3E). In the case of *AtTAS1c(NbmiR6019aTS)* precursors, only 39% of 21-nt [+] reads corresponded to the first register (Figure 3E).

**Figure 3.**
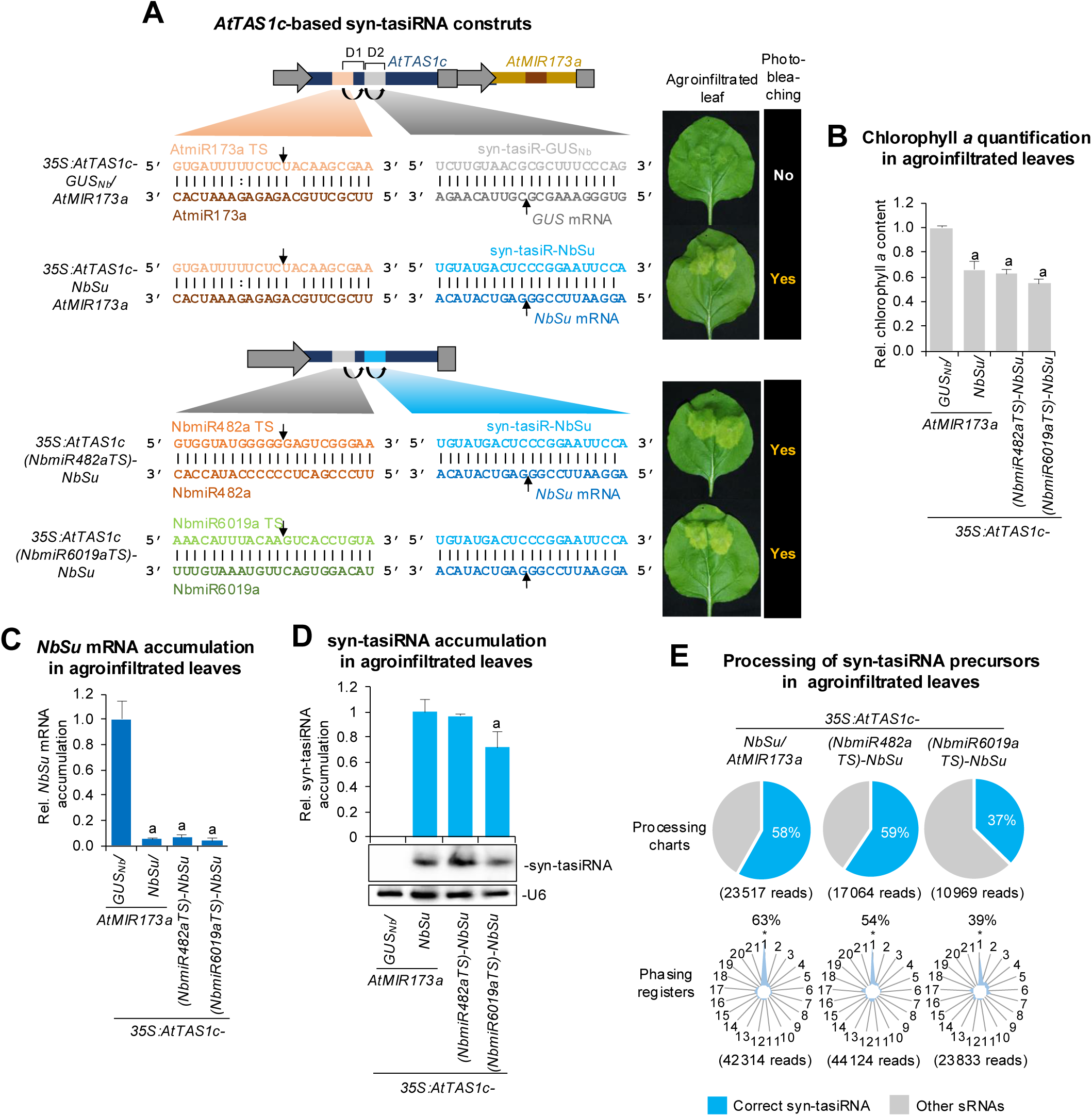
Functional analysis of syn-tasiRNAs against *N. benthamiana SULPHUR* (*NbSu*) expressed from modified *AtTAS1c* precursors including endogenous 22-nt miRNA target sites (TS). (**A**) Organization of *AtTAS1c*-based syn-tasiRNA constructs. Left: upper section shows constructs with canonical *AtTAS1c* precursors for expressing syn-tasiR-GUSNb or syn-tasiR-NbSu together with AtmiR173a; lower section shows constructs with modified *AtTAS1c* precursors including NbmiR482a or NbmiR6019 TSs. Nucleotides corresponding to AtmiR173a and AtmiR173aTS are shown in dark and light orange, respectively. Nucleotides corresponding to syn-tasiR-NbSu and target *NbSu* mRNA are shown in light and dark blue, respectively. Nucleotides corresponding to NbmiR482a and NbmiR482a TS are shown in light and dark brown, respectively. Nucleotides corresponding to NbmiR6019a/b and NbmiR6019a/b TS are shown in light and dark green, respectively. Other details as in Figure 1A. Right: photographs at 7 days post agroinfiltration (dpa) of leaves agroinfiltrated with each of construct. The presence or absence of bleaching on the agroinfiltrated patches is labelled as “Yes” or “No”, respectively. (**B**) Relative content of chlorophyll *a* in agroinfiltrated patches (*35S:AtTAS1c-AtMIR173-GUS_Nb_* = 1.0). Bars with letter “a” are significantly different from the control sample (P < 0.05 in pairwise Student’s t -test comparisons). (**C**) Target *NbSu* mRNA accumulation in agroinfiltrated leaves at 2 dpa [mean relative level (n = 3) + standard error] after normalization to *PROTEIN PHOSPHATASE 2A* (*NbPP2A*), as determined by quantitative RT-qPCR (*35S:AtTAS1c-AtMIR173-GUS_Nb_*= 1). Other details are as in B. (**D**) Northern blot detection of syn-tasiR-NbSu in RNA preparations from agroinfiltrated leaves at 2 dpa. The graph at top shows the mean + standard deviation (n = 3) syn-tasiRNA relative accumulation (*35S:AtTAS1c-AtMIR173-NbSu* = 1.0). Other details are as in Figure 1E. (**E**) Syn-tasiRNA processing and phasing analysis from *AtTAS1c*-derived precursors. Pie charts show the percentages of reads corresponding to expected, accurately processed 21-nt mature syn-tasiR-NbSu (in blue) or to other 19-24 nt sRNAs (in grey). Radar plots show the proportion of 21-nt reads corresponding to each of the 21 registers from *AtTAS1c* transcripts, with position 1 designated as immediately after AtmiR173a, NbmiR482a or NbmiR6019a/b guided cleavage site.

Next, we decided to test whether syn-tasiRNA biogenesis could be triggered in *N. benthamiana* from the *35S:NbmiR482aTS-NbSu* and *35S:NbmiR6019a/bTS-NbSu* constructs engineered for syn-tasiR-NbSu expression from minimal precursors including NbmiR482a or NbmiR6019a/b TS, respectively (Figure 4A). Both constructs were agroinfiltrated in *N. benthamiana* and analyzed as explained before together with *35S:AtTAS1-GUS_Nb_/AtMIR173a*, *35S:AtTAS1c(NbmiR482aTS)-NbSu* and *35S:AtTAS1c(NbmiR6019aTS)-NbSu* Figure 4A), analyzed in parallel for comparative purposes (Figure 4A). All constructs engineered for expressing syn-tasiR-NbSu induced bleaching in the agroinfiltrated areas (Figure 4A), which correlated with drastic *NbSu* mRNA downregulation compared to the negative control samples (Figure 4B). RNA blot assays confirmed the accumulation of syn-tasiR-NbSu as a single 21-nt sRNA species in all samples displaying bleaching, with samples expressing *NbmiR482aTS* precursors accumulating syn-tasiRNAs to levels similar to those of samples expressing *AtTAS1c(NbmiR482aTS)* precursors (Figure 4C). Moreover, the minimal *NbmiR482aTS* precursor was efficiently processed, as 63% of ±4 nt of 3’D2[+] reads corresponded to authentic syn-tasiR-NbSu. This percentage was similar to that of full-length *AtTAS1c* precursor (58%, Figure 3E) and slightly higher than that of the minimal *NbmiR6019a/b* precursor (Figure 4D).

**Figure 4.**
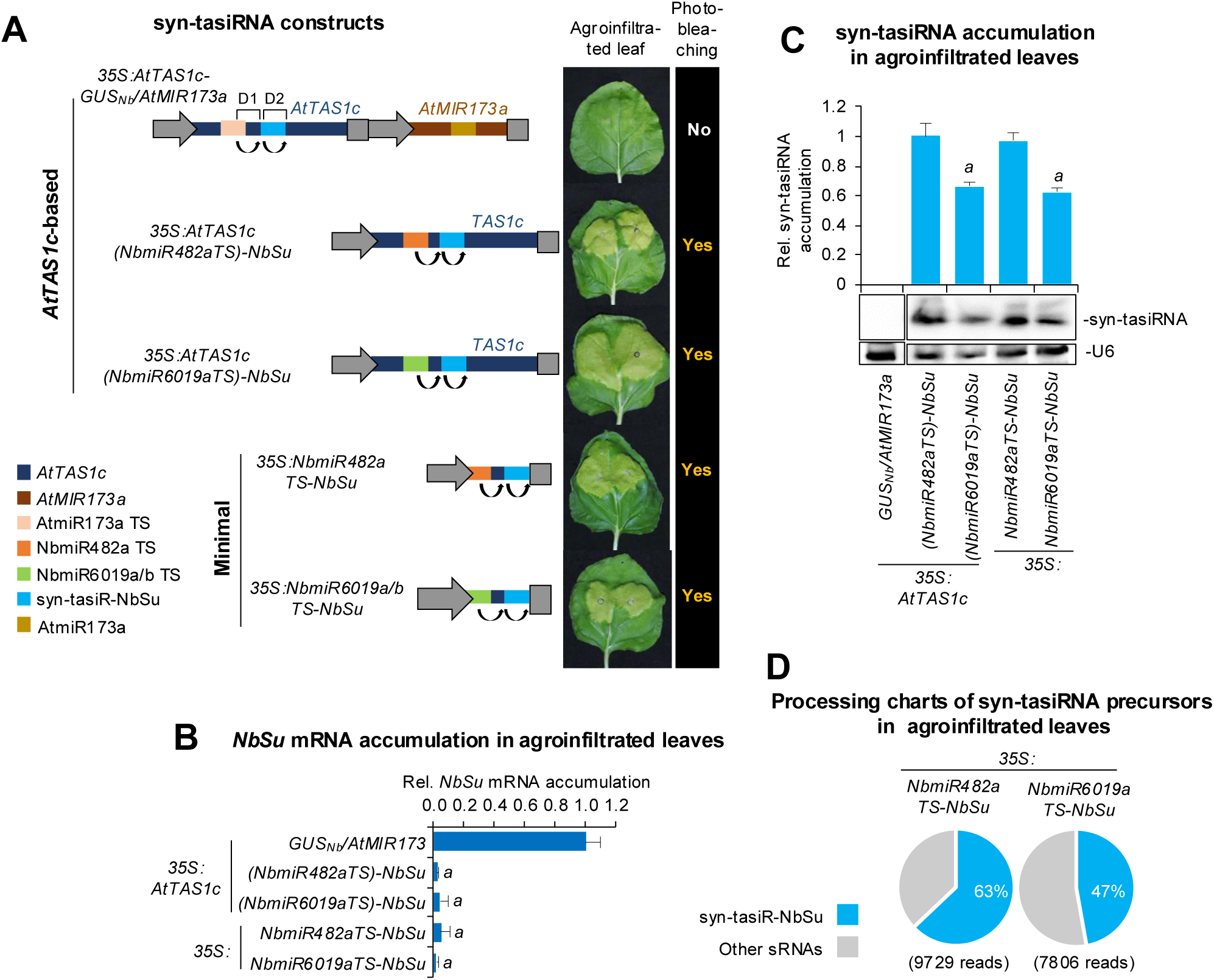
Functional analysis in *N. benthamiana* leaves of syn-tasiR-NbSu expressed from minimal precursors including NbmiR482a or NbmiR6019a/b target site (TS). (**A**) Diagram of *AtTAS1c*-based or minimal precursor constructs. NbmiR482a TS and NbmiR6019a/b TS are shown in brown and green boxes, respectively. Other details as in Figure 3A. (**B**) Target *NbSu* mRNA accumulation in agroinfiltrated leaves at 2 days post-agroinfiltration (dpa) [mean relative level (n = 3) + standard error] after normalization to *PROTEIN PHOSPHATASE 2A* (*NbPP2A*), as determined by RT-qPCR. Other details are as in Figure 3B. (**C**) Northern blot detection of syn-tasiR-NbSu in RNA preparations from agroinfiltrated leaves at 2 dpa. The graph at top shows the mean + standard deviation (n = 3) syn-tasiRNA relative accumulation [*35S:AtTAS1c(NbmiR482aTS)-NbSu* = 1.0]. Other details are as in Figure 1E. (**D**) Syn-tasiRNA processing from minimal precursors. Pie charts show the percentages of reads corresponding to expected, accurately processed 21-nt mature syn-tasiR-NbSu (in blue) or to other 19-24 nt sRNAs (in grey).

Finally, to confirm that syn-tasiRNA biogenesis from minimal precursors in *N. benthamiana* requires an endogenous 22-nt miRNA, leaves agroinfiltrated with the *35S:NbmiR156aTS-NbSu* and *35S:AtmiR173aTS-NbSu* constructs were analyzed. These constructs were engineered for expressing syn-tasiR-NbSu from minimal precursors including either a *N. benthamiana* 21-nt miRNA (NbmiR156a) TS or a heterologous 22-nt miRNA (AtmiR173a) TS, respectively (Figure S3A). Results show that leaves agroinfiltrated with either of these two constructs displayed neither bleaching (Figure S3A) nor reduced chlorophyll *a* content (Figure S3B), nor accumulation of syn-tasiR-NbSu (Figure S3C). These results also suggest that bleaching observed in leaves agroinfiltrated with *35S:NbmiR482aTS-NbSu* is due to the activity of syn-tasiR-NbSu rather than of potential siRNAs generated from transgene silencing. Taken together, all these findings demonstrate that highly phased, accurately processed syn-tasiRNAs can be produced in *N. benthamiana* from minimal precursors including a heterologous TS from an endogenous 22-nt miRNA.

### Widespread gene silencing in *N. benthamiana* triggered by syn-tasiRNAs derived from minimal precursors and expressed from a viral vector

The use of viral vectors to express syn-tasiRNAs for silencing plant genes has not been reported. Here, we tested this possibility and hypothesized that the 54-nt long *NbmiR482aTS*-based minimal precursor (when including a single syn-tasiRNA) could be more stable when inserted in a viral vector than the classic, 1011-nt long *AtTAS1c(NbmiR482aTS)*-based precursor. Minimal precursors including NbmiR482a TS were more accurately processed and produced increased syn-tasiRNA accumulation compared to NbmiR6019a/bTS (Figure 4C and 4D) most likely due to higher NbmiR482a abundance compared to precursors including NbmiR6019a TS (Figure S4). Thus, *NbmiR482aTS*-based precursors were preferred and analyzed in further experiments.

To analyze syn-tasiRNA biogenesis from a viral vector, *AtTAS1c(NbmiR482aTS)-NbSu* and *NbmiR482aTS-NbSu* sequences were inserted into a potato virus X (PVX) infectious clone to generate the *35S:PVX-AtTAS1c(NbmiR482aTS)-NbSu* and *35S:PVX-NbmiR482aTS-NbSu* constructs, respectively (Figure 5A). Both constructs were expected to express syn-tasiR-NbSu from full-length *AtTAS1c(NbmiR482aTS)* and minimal *NbmiR482aTS* precursors, respectively, during PVX infection and induce bleaching of PVX-infected tissues. These two constructs, along with the insert-free *35S:PVX* construct (Figure 5A), were independently agroinoculated into one leaf of three *N. benthamiana* plants. A negative control set of plants (“mock”) was infiltrated with agroinfiltration solution. The appearance of mild PVX-induced symptoms and bleaching silencing phenotypes were monitored during 28 dpa.

**Figure 5.**
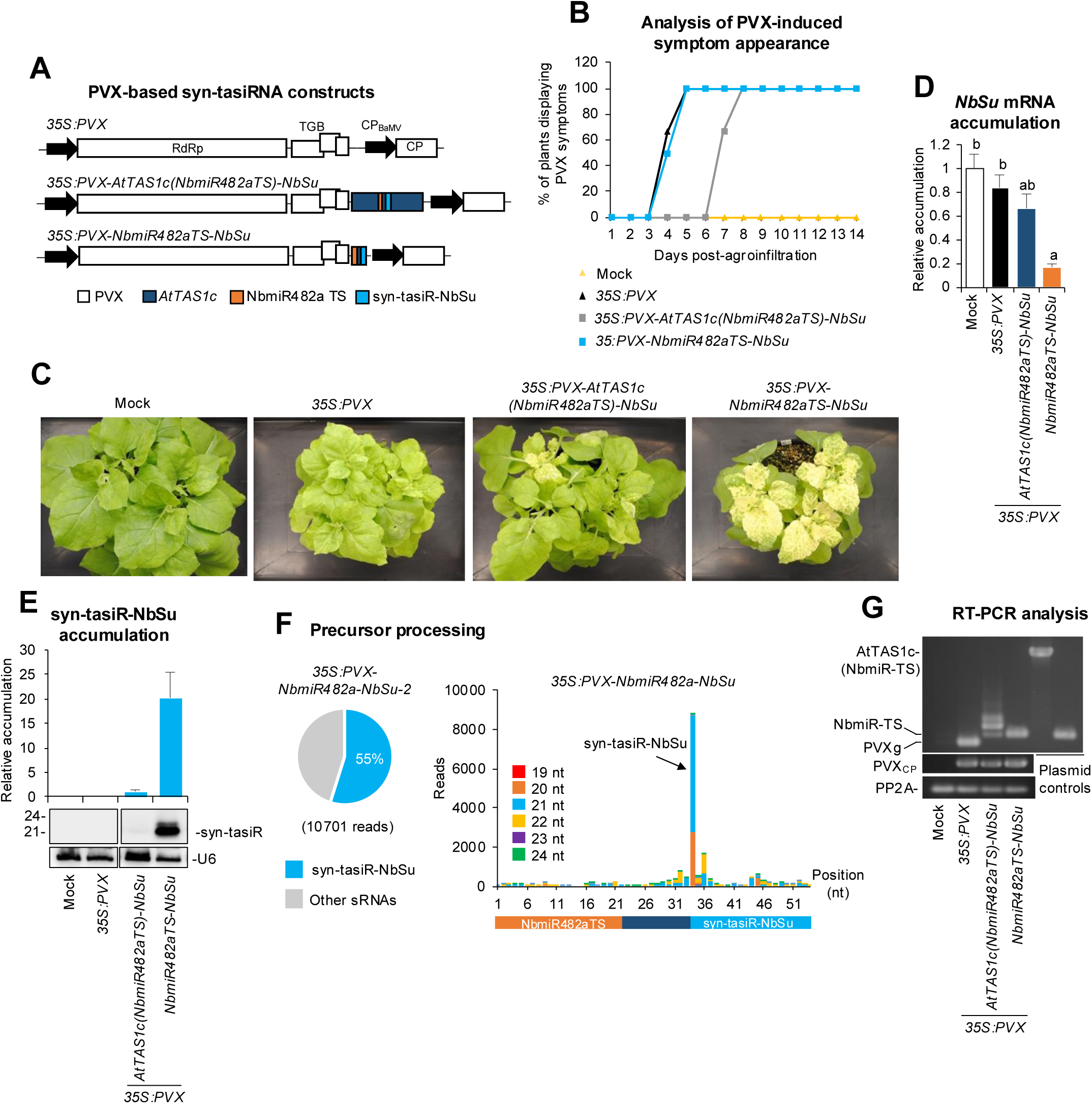
Functional analysis of potato virus X (PVX) constructs expressing syn-tasiR-NbSu from *AtTAS1c* or minimal syn-tasiRNA precursors in *N. benthamiana*. (**A**) Diagram of PVX-based constructs. *AtTAS1c*, NbmiR482aTS and syn-tasiR-NbSu sequences are represented by dark blue, orange and light blue boxes, respectively. PVX ORFs and promoters are represented as white boxes and black arrows, respectively. RdRP, RNA-dependent RNA-polymerase; TGB, triple gene block; CP, coat protein; CP_BaMV_, Bamboo mosaic virus CP promoter. (**B**) Two-dimensional line graph showing, for each of the three-plant sets listed, the percentage of symptomatic plants per day during 14 days. (**C**) Photos at 14 days post-agroinfiltration (dpa) of sets of three plants agroinoculated with the different constructs. (**D**) Target *NbSu* mRNA accumulation in RNA preparations from apical leaves collected at 14 dpa and analysed individually (mock = 1.0 in all comparisons). Bars with the letter ‘a’ or ‘b’ indicate whether the mean values are significantly different from mock control or *35S:PVX-NbmiR482aTS-NbSu* samples, respectively (*P* < 0.05 in pairwise Student’s *t*-test comparison). (**E**) Northern blot detection of syn-tasiR-NbSu in RNA preparations from apical leaves collected at 14 dpa and pooled from three independent plants. The graph at the top shows the mean + standard deviation (n = 3) syn-tasiRNA relative accumulation [*35S:AtTAS1c(NbmiR482aTS)-NbSu* = 1.0]. (**F**) syn-tasiRNA processing from *PVX-NbmiR482a-NbSu*. Left: pie chart shows the percentage of reads corresponding to expected, accurately processed 21-nt mature syn-tasiR-NbSu (in blue) or to other 19-24 nt sRNAs (in grey). Right: sRNA profile of 19-24 nt [+] reads mapping to each of the 54 nucleotide positions in the *NbmiR482aTS-NbSu* precursor from samples expressing *35S:PVX-NbmiR482aTS-NbSu*. Orange, dark blue and light blue boxes represent the nucleotides from NbmiR482aTS, *AtTAS1c*-derived spacer and syn-tasiR-NbSu, respectively. (**G**) RT-PCR detection of PVX, *AtTAS1c* and minimal precursors in apical leaves at 7 dpa. RT-PCR products corresponding to the *NbPP2A* and PVX vector controls are also shown (bottom), as well as control amplifications of *AtTAS1c* and minimal precursor fragments from plasmids (right). PVXg, band amplified from *35S:PVX* samples corresponding to the genomic region lacking a syn-tasiRNA precursor.

All plants agroinoculated with *35S:PVX* or *35S:PVX-NbmiR482aTS-NbSu* showed mild leaf curling typical of PVX-induced symptoms between 4 and 5 dpa (Figure 5B). In contrast, plants agroinoculated with *35S:PVX-AtTAS1c(NbmiR482aTS)-NbSu* displayed the curling phenotype 2-3 days later, between 7 and 8 dpa (Figure 5B). Interestingly, bleaching was observed in areas of certain apical leaves as soon as 7 dpa in plants agroinoculated with *35S:PVX-NbmiR482aTS-NbSu*, and by 14-21 dpa it extended to most of the apical tissues (Figure 5C). At these same timepoints, limited and mild bleaching was observed in plants agroinoculated with *35S:PVX-AtTAS1c(NbmiR482aTS)-NbSu* while no bleaching was observed in plants expressing *35S:PVX* (Figure 5C). RT-qPCR and RNA blot analyses showed that only tissues expressing *35S:PVX-NbmiR482aTS-NbSu* accumulated low levels of *NbSu* mRNA (Figure 5D) and high levels of syn-tasiR-NbSu, respectively (Figure 5E). In contrast, plants agroinoculated with *35S:PVX-AtTAS1c(NbmiR482aTS)-NbSu* displayed a slight decrease in *NbSu* mRNA levels and low levels of syn-tasiR-NbSu barely detectable by RNA blot assay (Figure 5E). Importantly, sRNA sequencing of RNA preparations from apical leaves of plants agroinoculated with *35S:PVX-NbmiR482aTS-NbSu* revealed that the *NbmiR482aTS-NbSu* precursor was accurately processed, with 55% of the ± 4 nt of 3’D2[+] reads corresponding to authentic syn-tasiR-NbSu (Figure 5F). Plotting all 19-24-nt sRNA [+] reads that map to the whole *NbmiR482aTS-NbSu* precursor revealed a relatively low number of sRNAs overlapping with the 3’D2[+] position or, more generally, produced from the rest of the precursor (Figure 5F). Indeed, the sRNA profile of 19-24-nt [+] reads mapping to the *NbmiR482aTS-NbSu* precursor was similar in *35S:PVX-NbmiR482aTS-NbSu* and *35S:NbmiR482aTS-NbSu* samples thus indicating that processing of the minimal precursor is not particularly altered during PVX replication (Figure S5). Finally, the presence of *AtTAS1c(NbmiR482aTS)-NbSu* and *NbmiR482aTS-NbSu* precursors was analyzed by RT-PCR at 7 dpa with oligonucleotides flanking the precursor insertion site. The 278-bp fragment corresponding to the *NbmiR482aTS-NbSu* precursors was clearly amplified while the 1235-bp fragment of the *AtTAS1c(NbmiR482aTS)-NbSu* could not be detected. PVX CP was detected in all samples expressing PVX-based constructs and *NbPP2A* in all samples. Sanger sequencing of RT-PCR fragments amplified from the three *PVX-NbmiR482aTS-NbSu*-infected samples revealed no mutations in the whole insert.

Importantly, to further confirm that *NbSu* silencing was due to syn-tasiR-NbSu activity and not to potential siRNAs generated from the *NbmiR482aTS-NbSu* precursor during PVX replication, a set of three plants were agroinoculated with the *35S:PVX-AtmiR173aTS-NbSu* construct (Figure S6A). As controls, sets of three plants were also mock-inoculated, or agroinoculated with the *35S:PVX* or *35S:PVX-NbmiR482aTS-NbSu* constructs. At 14 dpa, plants agroinoculated with the *35S:PVX-AtmiR173aTS-NbSu* construct did not display any bleaching (Figure S6B) or reduced *NbSu* mRNA levels (Figure S6C). These plants did not accumulate syn-tasiR-NbSu (Figure S6D), although they did accumulate PVX variants containing the *AtmiR173aTS-NbSu* precursors (Figure S6D). Overall, these results demonstrate that accurately processed syn-tasiRNAs can be produced from minimal precursors inserted into potato virus X, enabling widespread silencing of endogenous plant genes such as *NbSu*.

### Plant immunization against a pathogenic virus with antiviral syn-tasiRNAs produced from the potato virus X viral vector

Previous work showed that transgenically expressed syn-tasiRNAs induce enhanced antiviral resistance compared to amiRNAs due to the combined silencing effect of each individual syn-tasiRNA (35). Here, we explored the possibility of using syn-tasiR-VIGS to induce antiviral resistance against tomato spotted wilt virus (TSWV), an economically important plant pathogen affecting different crops worldwide (36), in *N. benthamiana*. To this end, we generated the *35S:PVX-NbmiR482aTS-TSWV(x4)* construct, which includes the 22-nt miR482aTS followed by the 11-nt *AtTAS1c* spacer and four different 21-nt syn-tasiRNA sequences with known anti-TSWV activity (20) (Figure 6A). A similar construct named *35S:PVX-NbmiR482aTS-GUS_Nb_(x4)*, expected to produce four innocuous anti-GUS syn-tasiRNAs of different sequences, was also generated (Figure 6A). Additionally, the *35S:PVX-amiR-GUS_Nb_*and *35S:PVX-amiR-TSWV* constructs, which are expected to produce amiR-GUS_Nb_ and amiR-TSWV amiRNAs (20), were generated for comparative purposes (Figure 6A). These constructs were agroinoculated into one leaf of nine *N. benthamiana* plants, six of which were mechanically inoculated seven days later with a TSWV infectious extract. An additional set of mock-inoculated plants was also included. To determine the antiviral activity of art-sRNAs, the appearance of typical TSWV-induced symptoms (leaf epinasty and chlorosis) in apical non-inoculated tissues was monitored over 21 days post-TSWV inoculation (dpi), and the presence of PVX and TSWV by RT-PCR or western blot.

**Figure 6.**
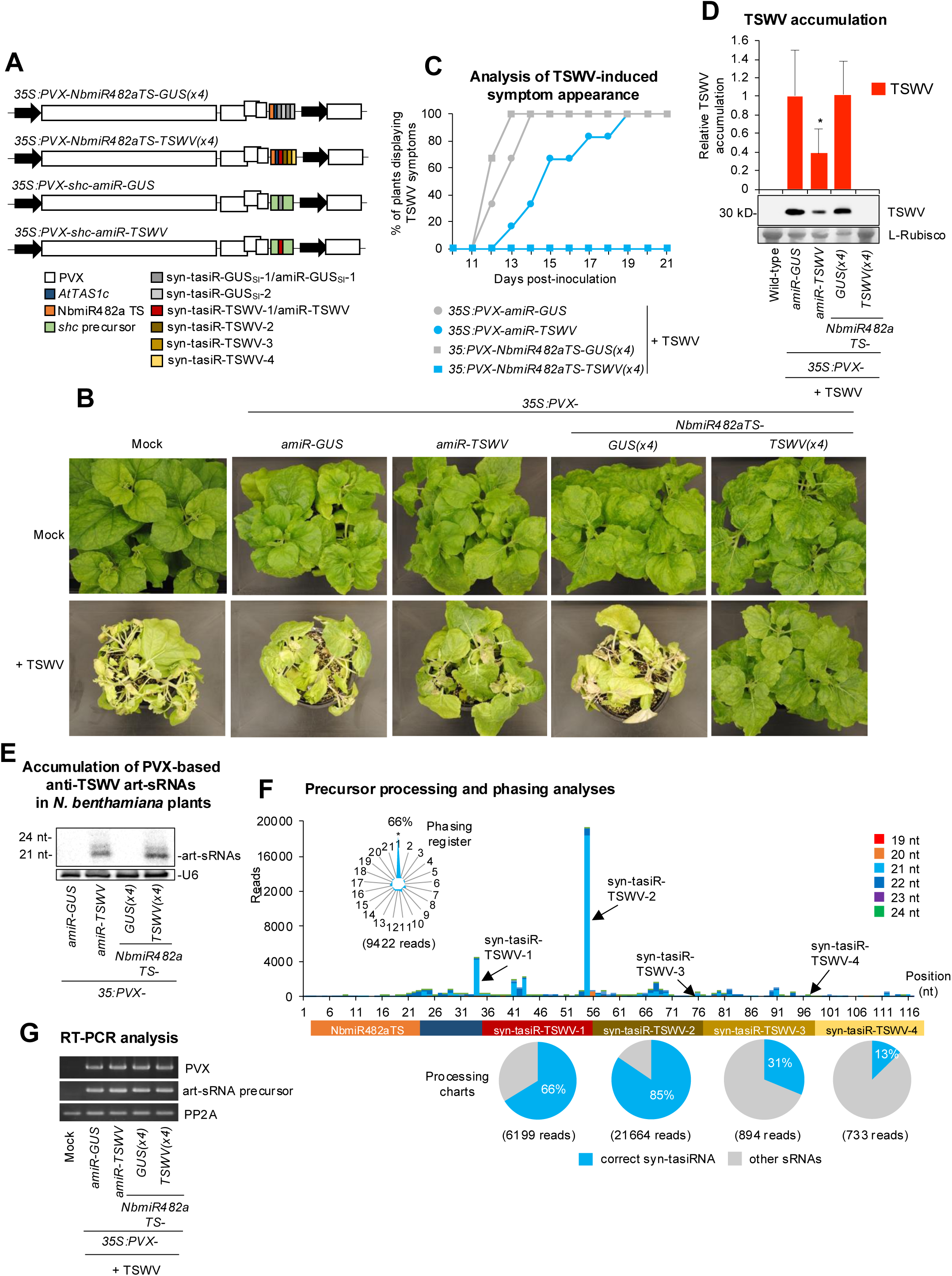
Functional analysis of potato virus X (PVX) constructs expressing syn-tasiRNAs against tomato spotted wilt virus in *N. benthamiana*. (**A**) Diagram of PVX-based constructs. Anti-TSWV art-sRNA sequences 1 (syn-tasiR-TSWV-1/amiR-TSWV), 2 (syn-tasiR-TSWV-2), 3 (syn-tasiR-TSWV-3) and 4 (syn-tasiR-TSWV-4) are represented by red, dark brown, light brown and yellow boxes, respectively. Anti-GUS art-sRNA sequences 1 (syn-tasiR-GUS_Nb_-1/amiR-GUS_Nb_-1) and 2 (syn-tasiR-GUS_Nb_-2) are represented by dark and light boxes, respectively. Other details are as in Figure 5A. (**B**) Photos at 21 days post-inoculation (dpi) of sets of three plants agroinoculated with the different constructs and inoculated or not (mock) with TSWV. (**C**) Two-dimensional line graph showing, for each of the six-plant sets listed, the percentage of symptomatic plants per day during 21 days. (**D**) Western blot detection of TSWV in protein preparations from apical leaves collected at 14 dpi and pooled from six independent plants. The graph at top shows the mean + standard deviation (n = 6) TSWV relative accumulation [*35S:PVX-amiR-GUS_Nb_* = 1.0]. Bars with the letter ‘a’ are significantly different from that of *35S:PVX-amiR-GUS_Nb_*control sample (P < 0.05 in pairwise Student’s t-test comparison). The membrane stained with Ponceau red showing the large subunit of Rubisco (ribulose1, 5-biphosphate carboxylase/oxygenase) is included as loading control. (**E**) Northern blot detection of anti-TSWV art-sRNAs in RNA preparations from apical leaves collected at 7 dpa and pooled from three independent mock-inoculated plants. (**F)** syn-tasiRNA processing from *PVX-NbmiR482a-TSWV(x4)*. Top: sRNA profile of 19-24 nt [+] reads mapping to each of the 117 nucleotide positions in the *NbmiR482aTS-TSWV(x4)* precursor from samples expressing *35S:PVX-NbmiR482aTS-TSWV(x4)*. Other details are as in A. Bottom: pie charts showing the percentages of reads corresponding to expected, accurately processed 21-nt mature forms of each of the four anti-TSWV syn-tasiRNAs (in blue) or to other 19-24 nt sRNAs (in grey). (**G**) RT-PCR detection of PVX and minimal precursors in apical leaves at 7 dpa. RT-PCR products corresponding to the *NbPP2A* and PVX vector controls are also shown (bottom).

None of the plants agroinoculated with the *35S:PVX-NbmiR482aTS-TSWV(x4)* construct displayed typical TSWV-induced symptoms throughout the experiment, while plants expressing *35S:PVX-amiR-TSWV* became symptomatic, although with a significant one to six-day delay compared to plants agroinoculated with control constructs (Figure 6B and 6C). At 14 dpi, TSWV was undetectable in protein extracts from plants agroinoculated with *35S:PVX-NbmiR482aTS-TSWV(x4)*, while high or low levels of TSWV were detected in control and amiR-TSWV-expressing plants, respectively (Figure 6D). Importantly, PVX variants including the corresponding art-sRNA precursor were detected by RT-PCR, indicating that the lack of protection was not due to the lack of PVX infection or the loss of the art-sRNA precursor (Figure 6E). Finally, northern blot analysis confirmed art-sRNA accumulation in plants agroinoculated with PVX-based constructs, with both amiR-TSWV and anti-TSWV syn-tasiRNAs accumulating predominantly as 21-nt sRNA species (Figure 6F). The accuracy of *NbmiR482aTS-TSWV(x4)* precursor processing and the production of authentic anti-TSWV syn-tasiRNAs were analyzed by high-throughput sequencing of sRNA libraries from plants agroinoculated with *35S:PVX-NbmiR482aTS-TSWV(x4)* (Figure 6G). All four syn-tasiRNA sequences were detected as predominant when plotting all 19-24-nt sRNA [+] reads mapping to the precursor, with x%, x%, x% and x% of reads within ± 4 nt of 3’D2 [+], 3’D3 [+], 3’D4 [+] and 3’D5 [+], respectively, corresponded to authentic syn-tasiRNAs (Figure 6G). Additionally, highly phased syn-tasiRNAs were generated, with 63% of 21-nt [+] reads corresponding to the first register (Figure 6G). Taken together, these findings indicate that PVX-based syn-tasiR-VIGS can be used to vaccinate plants against TSWV for complete immunization. They also highlight that multiple syn-tasiRNAs can be produced simultaneously *in planta* from an RNA viral vector such as PVX.

### Transgene-free, PVX-based syn-tasiR-VIGS

We finally explored the possibility of applying syn-tasiR-VIGS to plants in a DNA-free, non-transgenic manner for both silencing endogenous genes or for plant vaccination (Figure 7A). To do so, six *N. benthamiana* plants were independently agroinoculated with either *35S:PVX*, *35S:PVX-NbmiR482a-NbSu*, *35S:PVX-NbmiR482aTS-GUS_Nb_(x4)* or *35S:PVX-NbmiR482aTS-TSWV(x4)*. At 6 dpa, apical leaves displaying mild PVX-induced symptoms were collected together and a crude extract was prepared for each case (Figure 7A).

**Figure 7.**
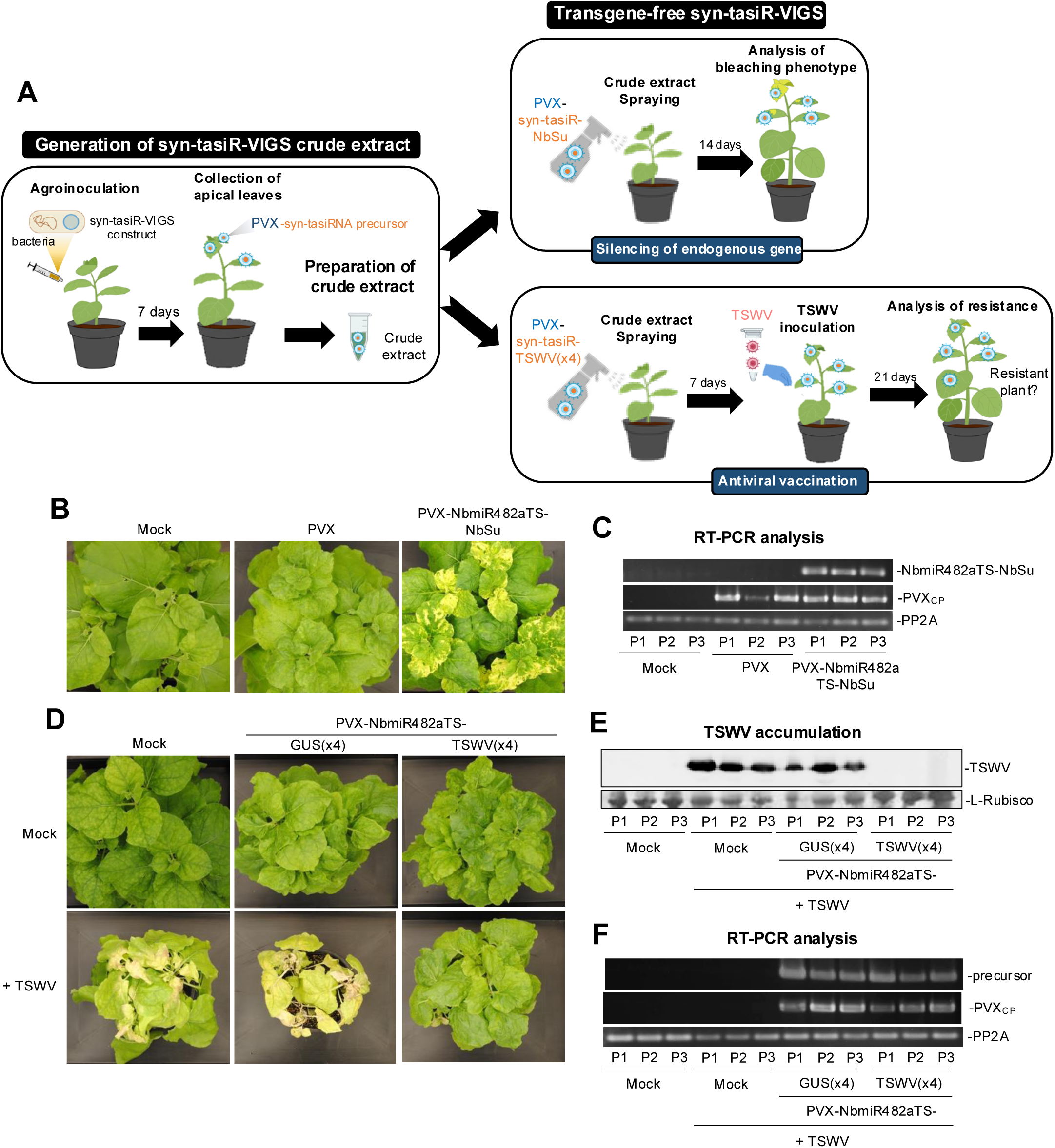
Transgene-free widespread gene silencing and plant antiviral vaccination through PVX-based syn-tasiR-VIGS. (**A**) Experimental procedure for transgene-free syn-tasiR-VIGS in *N. benthamiana* plants. Left: crude extracts are prepared from plants previously agroinfiltrated with the corresponding syn-tasiR-VIGS construct. Right: young plants are spray-inoculated with syn-tasiR-VIGS extracts to induce bleaching derived from *NbSu* silencing (top) or antiviral resistance against TSWV (bottom). (**B**) Widespread silencing of *NbSu* induced by sprayed crude extracts. Photographs at 14 days post-spray (dps) of sets of three plants sprayed with different crude extracts obtained from agroinoculated plants. (**C**) RT-PCR detection at 14 dps of *NbmiR482aTS-NbSu* precursors and PVX coat protein fragment (PVX_CP_) in apical leaves of each of the three sprayed plants (P1-P3). RT-PCR products corresponding to the control *NbPP2A* are also shown. (**D**) Plant vaccination with syn-tasiR-VIGS extracts for immunization against TSWV. Photographs at 21 days post-inoculation (dpi) of sets of three plants sprayed with different crude extracts obtained from agroinoculated plants. (**E**) Western blot detection of TSWV in protein preparations from apical leaves collected at 14 dpi of each of the three sprayed plants (P1-P3). Other details are as in Figure 6D. (**F**) RT-PCR detection at 14 dpi of *NbmiR482aTS-TSWV(x4)* precursors and PVX coat protein fragment (PVX_CP_) in apical leaves of each of the three plants sprayed (P1-P3). Other details are as in Figure 5G.

First, crude extracts including PVX-NbmiR482aTS-NbSu were sprayed onto three *N. benthamiana* plants and the appearance of bleaching was monitored over 14 days (Figure 7A). Control crude extracts from mock-and empty PVX-agroinoculated plants were sprayed in parallel. Remarkably, all three *N. benthamiana* plants sprayed with crude extracts derived from *35S:PVX-NbmiR482a-NbSu* displayed bleaching indicative of *NbSu* silencing (Figure 7B) and accumulated minimal syn-tasiRNA precursors (Figure 7C), with no sequence alterations as confirmed by Sanger sequencing of RT-PCR products. Control plants sprayed with empty extracts displayed mild PVX symptoms but no bleaching and accumulated PVX RNAs, while plants sprayed with mock extracts were symptomless as well as bleaching-and virus-free (Figure 7B and C).

Next, crude extracts containing PVX-NbmiR482aTS-TSWV(x4) were sprayed onto three *N. benthamiana* plants, and seven days later the plants were mechanically inoculated with a TSWV infectious extract. TSWV-induce symptoms were monitored over 21 dpi. Control crude extracts from mock- and *35S:PVX-NbmiR482aTS-GUSNb(x4)*-agroinoculated plants were sprayed in parallel, and plants were inoculated with TSWV as described. Plants vaccinated with PVX-NbmiR482aTS-TSWV(x4) extracts did not show any TSWV symptom and did not accumulate TSWV (Figure 7D and E), while plants treated with mock or PVX-NbmiR482aTS-GUSNb(x4) extracts displayed severe plant chlorosis and leaf epinasty and accumulated high levels of TSWV (Figure 7D and E). As before, the presence of PVX and *NbmiR482aTS*-based precursors was analyzed by RT-PCR (Figure 7F), confirming that the lack of protection in plants treated with PVX-NbmiR482aTS-GUSNb(x4) extracts was not due to the absence of PVX or the loss of the syn-tasiRNA precursor. Collectively, these findings demonstrate that crude extracts can be directly sprayed onto leaves to trigger syn-tasiR-VIGS in *N. benthamiana* in a DNA-free, non-transgenic manner, enabling both silencing of endogenous genes and plant vaccination against pathogenic viruses.

## Discussion

Here, we show that minimal RNA precursors consisting exclusively of a 22-nt miRNA TS and an 11-nt *AtTAS1c*-derived spacer produce high levels of accurately processed syn-tasiRNAs in different plant species for efficient gene silencing. Remarkably, minimal precursors have the unique ability to express authentic syn-tasiRNAs from an RNA viral vector such as PVX, enabling transgene-free widespread gene silencing in plants.

### Effect of minimizing the precursor length on syn-tasiRNA biogenesis and function in Arabidopsis

The effect on secondary siRNA biogenesis of deleting sequences from *AtTAS* precursors was examined before when looking for regulatory functions of putative *cis* elements included in *AtTAS* sequences. For instance, the 930-nt full-length *AtTAS1a* was shortened to 252-nt by removing the 5’ region upstream of the AtmiR173a TS and/or the region downstream of the tasiRNA 3’D9[+] position (37). None of the deleted sequences were determinant for syn-tasiRNA biogenesis, although syn-tasiRNA accumulation was negatively affected. In a different study, the 5’ region upstream of AtmiR173a TS was also deleted from *AtTAS1c* without a clear effect on syn-tasiRNA accumulation (14). However, it was proposed that other precursor elements such as the poly(A) tail or the total transcript length may be crucial for proper syn-tasiRNA accumulation. In any case, both studies suggest that the presence of AtmiR173a TS is sufficient for triggering tasiRNA biogenesis. Indeed, phased tasiRNAs were generated from gene fragments placed downstream a AtmiR173a TS without the need for any regulatory sequences (14, 37). Here, syn-tasiRNAs accumulated to lower levels in Arabidopsis plants expressing the minimal precursor compared to those expressing the full-length *AtTAS1c*, suggesting the existence of *cis* elements in *AtTAS1c* that positively regulate syn-tasiRNA accumulation. Remarkably, some *TAS* genes have short open reading frames (ORFs) located immediately upstream of the tasiRNA-producing region, which are translated and trigger the synthesis of small peptides (38, 39). Indeed, mutations affecting either the stop codon or the overall ORF length lower tasiRNA accumulation, most likely due to a decreased stability of the *TAS* precursor caused by lower protection from ribosomes (40, 41). More recently, a model was proposed in which ribosomes start translating the ORFs and stall near the miRNA TS through the interaction with the SGS3-AGO1-miRNA complex (42). This interaction is thought to stabilize the complex, thus increasing the amount of tasiRNA produced. In the case of the *AtTAS1c* precursor, two ORFs are located in the 5’ region upstream of the AtmiR173a TS (ORF1 and ORF2), and a third ORF overlaps with ORF2 and includes the miRNA TS (39). These findings may explain the higher accumulation of syn-tasiR-AtFT and syn-tasiR-AtCH42 in Arabidopsis plants expressing full-length *AtTAS1c* precursors compared to those expressing ORF-deficient minimal precursors. Still, these differences in syn-tasiRNA accumulation seem to be less drastic than those reported before (14, 37), maybe because here syn-tasiRNAs are expressed from the 3’D2[+] position, which maximizes both syn-tasiRNA accumulation and gene silencing efficiency (18). Intriguingly, syn-tasiRNAs expressed from minimal precursors accumulated to similar levels than those expressed from *AtTAS1c* in *N. benthamiana*. It is possible that *AtTAS1c* ORFs are not recognized in *N. benthamiana* (where the *TAS1c*/miR173a pathway is missing) and, consequently, full-length *AtTAS1c* precursors are not protected by ribosomes. Finally, highly phased syn-tasiRNAs were produced from minimal precursors, indicating an accurate processing of the minimal precursors that prevents off-target effects caused by misprocessed sRNA species. Indeed, phasing maintenance during the processing of the minimal precursors is consistent with previous studies indicating that phasing depends on miRNA-AGO cleavage of the *TAS* transcript rather than on any *cis* regulatory element (43).

### Highly phased, accurately processed syn-tasiRNAs are produced in *Nicotiana benthamiana* with precursors including TSs from 22-nt endogenous miRNAs

The co-expression of *AtMIR173a* together with *AtTAS1c*-based syn-tasiRNA constructs is required to produce syn-tasiRNAs in non-Arabidopsis species such as *N. benthamiana* and *Solanum lycopersicum* (18, 23, 35). Here, highly abundant and phased syn-tasiRNAs were produced in *N. benthamiana* from full-length *AtTAS1c*-based precursors including a TS from *N. benthamiana* endogenous 22-nt miRNAs. This result indicates that the AtmiR173a TS is not essential for *AtTAS1c*-dependent syn-tasiRNA biogenesis, and that alternative TSs from other 22-nt miRNAs can be used. *N. benthamiana* has seven miRNA families with 22-nt miRNA members. Among them, NbmiR482a and NbmiR6019a/b are present in members of the *Solanaceae* family, play roles in immune responses to pathogens, and trigger phased siRNA formation (44–47). NbmiR482a is highly expressed in *N. benthamiana* seedlings, leaves and stems, while NbmiR6019a/b expression in these tissues is more modest (46) (Figure S4). These differences in miRNA expression may explain the higher accumulation of syn-tasiRNAs produced from precursors with NbmiR482a TS. Interestingly, syn-tasiRNA accumulation from *AtTAS1c* full-length precursors and gene silencing efficiency were similar in plants overexpressing AtmiR173a and in those using NbmiR482a as the endogenous trigger, indicating that NbmiR482a levels are not limiting. It is worth noting that miRNA expression profiles are dynamic and vary between tissues, developmental stages and during infections (48). Therefore, selecting the appropriate miRNA as endogenous trigger may require understanding its expression pattern across tissues or under different plant growth conditions. On the other hand, the specific expression profiles of certain miRNAs may allow syn-tasiRNA biogenesis to be tissue- or condition-specific, which could be an attractive strategy to pursue in specific cases.

### Transgene-free syn-tasiR-VIGS for widespread gene silencing in plants

Syn-tasiR-VIGS was developed by incorporating a minimal precursor with an endogenous 22-nt miRNA TS in the genome of a viral vector such as PVX. As previously observed with PVX-based amiR-VIGS (26), long precursors such as *AtTAS1c* (1011 nt) are not stably maintained in the viral genome for extended periods, highlighting the limited cargo capacity of viral vectors. In this context, minimal precursors offer a unique advantage due to their small size, allowing stable maitenance in the viral genome while reducing the accumulation of mutations during viral replication. Here, we used a PVX cDNA sequence with a deletion of the amino-terminal end of the coat protein (CP) and a heterologous promoter derived from bamboo mosaic virus (BaMV) (49), enhancing insert stability in PVX-based constructs. Remarkably, PVX-based syn-tasiRNAs were accurately processed from minimal precursors and accumulated to high levels, as shown by high-throughput sRNA sequencing and northern blot. Since PVX replicates in the cytoplasm, it is plausible that DCL4, a cytoplasmic DCL, is the main DCL processing syn-tasiRNA duplexes from dsRNAs generated after NbmiR482a cleavage, as occurs during the synthesis of endogenous tasiRNAs (50, 51). On the other hand, DCL4 is considered the primary antiviral DCL (52, 53), particularly during PVX infections in *N. benthamiana* (54), and together with DCL2 and DCL3 processes viral dsRNA replicative intermediates into virus-derived siRNAs as part of the natural plant’s antiviral defense. Here, sRNA sequencing confirmed the *in vivo* production of high levels of authentic 21-nt syn-tasiRNAs phased with the NbmiR482a cleavage site, with a relatively low proportion of sRNAs derived from the rest of the precursor. These results suggest that syn-tasiRNAs are produced from RDR6/SGS3-dependent dsRNAs post-NbmiR482a cleavage rather than from RDR-derived viral replicative intermediates. They also support that target silencing is a direct result of syn-tasiRNA function, not of potential siRNAs generated from the minimal precursor that might share sequence complementarity with target mRNAs. This is further supported by the lack of *NbSu* silencing observed when expressing PVX-based constructs that include AtmiR173a TS upstream the syn-tasiR-NbSu sequence. In any case, NbmiR482a targeting and subsequent processing of minimal precursors from PVX are efficient enough to produce high levels of syn-tasiRNAs while allowing continued PVX replication to sustain syn-tasiRNA synthesis over time and across new tissues.

A key feature of syn-tasiR-VIGS is its scalability and non-transgenic application via the spraying of plant leaves with infectious crude extracts from plants accumulating the viral vector with the minimal precursor, as shown recently for PVX-based amiR-VIGS (26). Previously, syn-tasiRNAs were expressed from transgenic tissues, which limited their practical applications due to commercial and/or regulatory constraints. Another common limitation of the syn-tasiRNA technology has been the need of co-expressing AtmiR173a together with *AtTAS1c*-based constructs in non-Arabidopsis species. The finding that authentic syn-tasiRNAs can be efficiently produced from minimal precursors consisting exclusively of an endogenous 22-nt miRNA TS and an 11-nt spacer significantly broadens the biotechnological applications of syn-tasiRNAs. Moreover, the new set of “B/c” plasmids listed in Figure S7 simplifies in a time-and cost-effective manner the generation of constructs containing minimal syn-tasiRNA precursors, especially when combined with high-throughput cloning methodologies that eliminate the need for gel purification and amplification steps, employing zero-background and Golden Gate strategies (16).

### New generation of antiviral vaccines based on syn-tasiR-VIGS

Traditional antiviral vaccines for plants rely on cross-protection, where a mild or attenuated virus strain protects the plant against more severe strains of the same or a closely related virus (55). However, identifying mild strains for a particular virus may can be challenging, and their protective effect is not always guaranteed. Moreover, cross-protection is limited to closed related viruses and does not defend against distantly related or unrelated viruses, limiting its effectiveness in fields with diverse viral populations. Here, we developed of a new generation of antiviral vaccines for plants based on syn-tasiR-VIGS, which can be applied in a non-transgenic and single-dose manner. Using PVX as viral vector, we vaccinated *N. benthamiana* plants for complete immunization against an entirely unrelated virus, TSWV, one of the top 10 plant viruses based on scientific/economic importance (36). Vaccination consisted in inoculating plants with PVX crude extracts, allowing the virus to spread and produce anti-TSWV syn-tasiRNAs throughout the plant. When a few days later TSWV is inoculated, vaccinated plants were loaded with antiviral syn-tasiRNAs that targeted TSWV and ultimately blocked its infection. Syn-tasiR-VIGS vaccines offer several advantages over traditional vaccines: i) syn-tasiRNAs can be designed *a la carte* against any virus of interest, ii) they may be applicable to any host-virus combination, as long as the viral vector can infect the host, and iii) they could protect against multiple unrelated viruses by multiplexing different antiviral syn-tasiRNAs in the same minimal precursor. Further research in these areas should expand the applications of syn-tasiR-VIGS vaccines and position them as a next-generation antiviral strategy for protecting plants against pathogenic viruses.

## Data availability

All original data will be made available upon request. High-throughput sequencing data can be found in the Sequence Read Archive (SRA) database under accession number PRJNA1172601. New B/c and PVX vectors are available from Addgene: *pENTR-B/c* (Addgene plasmid #227962, https://www.addgene.org/227962), *pMDC32B-B/c* (Addgene plasmid #227963, https://www.addgene.org/227963), *pENTR-AtmiR173aTS-B/c* (Addgene plasmid #227964, https://www.addgene.org/227964), *pMDC32B-AtmiR173aTS-B/c* (Addgene plasmid #227965, https://www.addgene.org/227965), *pENTR-NbmiR482aTS-B/c* (Addgene plasmid #227966, https://www.addgene.org/227966), *pMDC32B-NbmiR482aTS-B/c* (Addgene plasmid #227967, https://www.addgene.org/227967).

## Supplementary data

Supplementary Data are available at NAR Online.

## Acknowledgements

We thank José-Antonio Daròs (IBMCP) for sharing the *pLBPVXBa-M* plasmid, Javier Forment (IBMCP) for his assistance with the sRNA sequencing data analysis, and the greenhouse staff at IBMCP for their help in maintaining the plants.

## Author contributions

A.E.C. and A.A.G. did most of the experimental work with the help of J.L.G. M.J.M., A.P.E. and A.P. A.E.C., A.A.G. and A.C. analyzed the data. A.C. conceived the research, supervised the project and wrote the manuscript with input from the rest of authors.

## Funding

This work was supported by grants or fellowships from MCIN/AEI/10.13039/501100011033 and/or by the “European Union NextGenerationEU/PRTR” [PID2021-122186OB-100, CNS2022-135107 and RYC-2017-21648 to A.C.; PRE2019-088439, PRE2022-102565 and PRE2022-103177 to A.E.C., J.L.G. and M.J.M, respectively], from Consejo Superior de Investigaciones Científicas (CSIC, Spain) [JAEINT_20_01312 to A.P.E.] and from European Commission [Erasmus+ Grant Agreement 2020-1-DE01-KA103-005653 to A.P.].

## Conflict of interest

None declared.

## Notes

### Competing Interest Statement

The authors have declared no competing interest.

