## Supplemental information for "Syn-tasiR-VIGS: Virus-Based Targeted RNAi in Plants By Synthetic Trans Acting Small Interfering RNAs Derived from Minimal Precursors"

### SUPPLEMENTARY DATA

Supplementary Data are available at NAR Online.

**Data S1**. sRNA reads from art-sRNA-expressing tissues.

**Figure S1.** Direct syn-tasiRNA cloning in *B/c* (*Bsa*I/*ccd*B)-based vectors including a *ccd*B cassette flanked by two *Bsa*I sites.

**Figure S2.** Direct syn-tasiRNA cloning downstream the AtmiR173a or NbmiR482a target sites (TSs) in *B/c* (*Bsa*I/*ccd*B)-based vectors including a *ccd*B cassette flanked by two *Bsa*I sites.

**Figure S3.** Functional analysis of syn-tasiRNAs against *N. benthamiana SULPHUR* (*NbSu*) expressed from modified minimal precursors including different miRNA target sites (TS).

**Figure S4.** Analysis of NbmiR482a and NbmiR6019a presence in *Nicotiana benthamiana* tissues.

**Figure S5.** Mapping of 19-24-nucleotide small RNA reads to *NbmiR482aTS-NbSu* precursors.

**Figure S6.** Functional analysis of potato virus X (PVX) constructs expressing syn-tasiR-NbSu from minimal precursors including endogenous or heterologous 22-nt miRNA target sites (TSs).

**Figure S7.** *B/c*-based vectors for direct cloning of syn-tasiRNAs downstream the AtmiR173a and NbmiR482a target sites (TSs), or the TS of choice.

**Table S1.** Name, sequence and use of DNA oligonucleotides used in this study.

**Table S2:** Phenotypic penetrance of syn-tasiRNAs expressed in *A. thaliana* Col-0 T1 transgenic plants for silencing *FT*.

**Table S3:** Phenotypic penetrance of syn-tasiRNAs expressed in *A. thaliana* Col-0 T1 transgenic plants for silencing *CH42*.

**Table S4:** Phenotypic penetrance of syn-tasiRNAs expressed in Arabidopsis Col-0 T1 transgenic plants for silencing *FT* and *TRY*.

**Text S1.** Protocol to design and clone syn-tasiRNAs downstream the 3'D1[+] position in *Bsa*I/*ccd*B-based ('B/c') vectors *pENTR-B/c*, *pMDC32-B/c*, *pENTR-AtmiR173aTS-B/c*, *pMDC32B-AtmiR173aTS-B/c*, *pENTR-NbmiR482aTS-B/c* and *pMDC32B-NbmiR482aTS-B/c*.

**Text S2.** Protocol to generate PVX-based syn-tasiRNA constructs.

**Text S3**. DNA sequence in FASTA format of all precursors used to express art-sRNAs in plants.

**Text S4.** DNA sequence of *Bsa*I-*ccd*B-based (B/c) vectors used for direct cloning of syn-tasiRNAs.

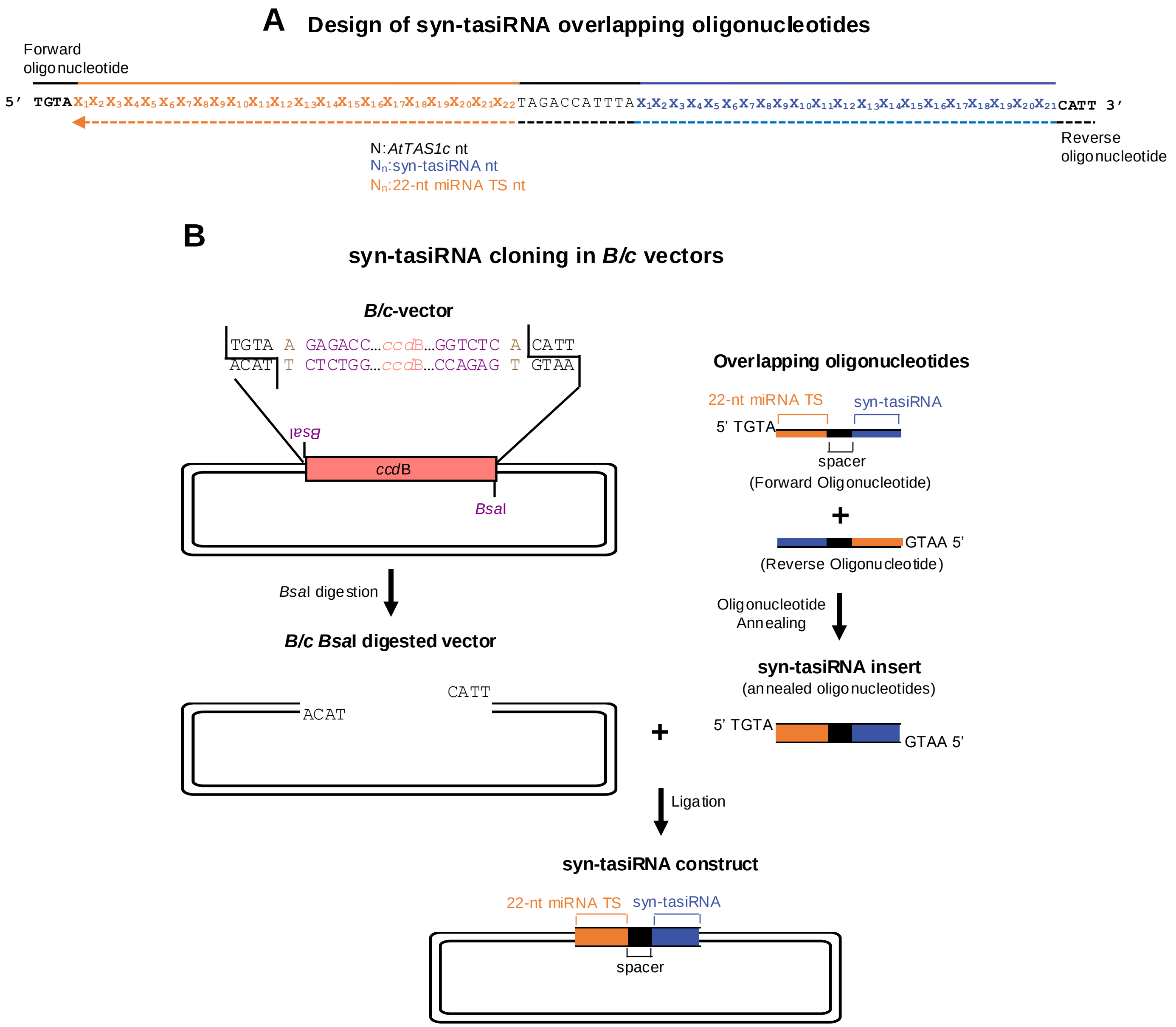


**Figure S1**. Direct syn-tasiRNA cloning in *B/c* (BsaI/ccdB)-based vectors including a *ccd*B cassette flanked by two *Bsa*I sites. (**A**) Design of two overlapping oligonucleotides for syn-tasiRNA cloning. Sequence covered by the forward and reverse oligonucleotides are represented with continuous or dotted lines, respectively. Nucleotides of the 22-nt miRNA target site, the *AtTAS1c*-derived spacer and the syn-tasiRNA sequences are in orange, black and blue, respectively. Oligonucleotide 5' overhangs are in black and bold. (**B**) Diagram of the steps for syn-tasiRNA cloning in *B/c* vectors. The syn-tasiRNA insert, including the 22-nt miRNA target site sequence of interest followed by a 11-nt *AtTAS1c*-derived spacer, obtained after annealing the two overlapping oligonucleotides has 5' TGTA and 5'-AATG overhangs and is directly inserted into the *Bsa*I-linearized *B/c* vector. Nucleotides of the *Bsa*I sites and arbitrary nucleotides used as spacers between the *Bsa*I recognition site and the *AtTAS1c* sequence are in purple and light brown, respectively. Other details are as in A.

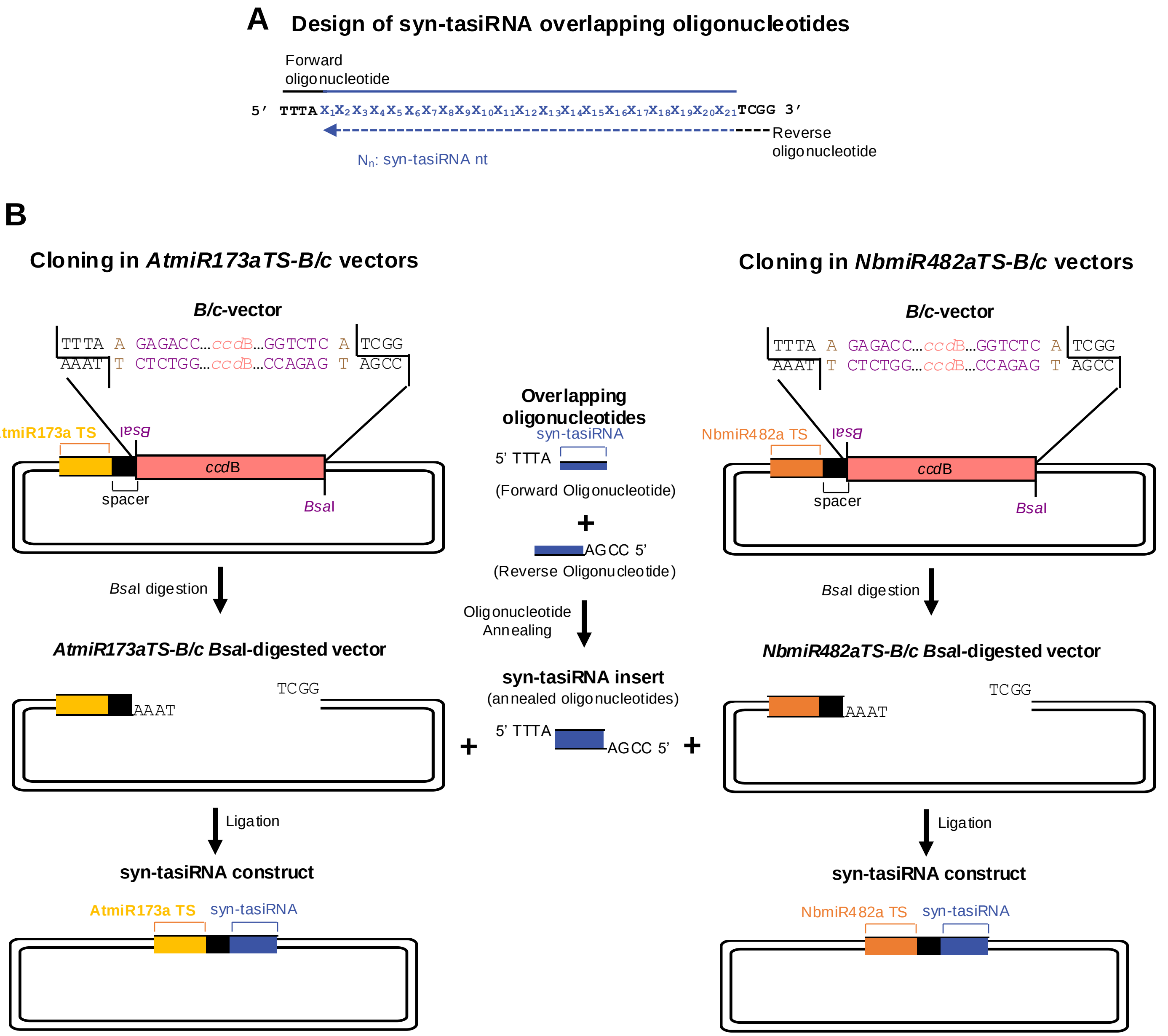


**Figure S2**. Direct syn-tasiRNA cloning downstream the AtmiR173a or NbmiR482a target sites (TSs) in *B/c* (*Bsa*I/*ccd*B)-based vectors including a *ccd*B cassette flanked by two *Bsa*I sites. (**A**) Design of two overlapping oligonucleotides for syn-tasiRNA cloning. Sequence covered by the forward and reverse oligonucleotides are represented with continuous or dotted lines, respectively. Nucleotides of the syn-tasiRNA sequence are in blue, and oligonucleotide 5' overhangs are in black and bold. (**B**) Diagram of the steps for syn-tasiRNA cloning in *AtmiR173aTS-B/c* or *NbmiR482aTS-B/c* vectors. The syn-tasiRNA insert obtained after annealing the two overlapping oligonucleotides has 5' TTTA and 5'-CCGA overhangs and is directly inserted into the *Bsa*I-linearized *AtmiR173aTS-B/c*- or *NbmiR482aTS-B/c*-based vectors. Nucleotides of the *Bsa*I sites and arbitrary nucleotides used as spacers between the *Bsa*I recognition site and the *AtTAS1c* sequence are in purple and light brown, respectively. Nucleotides of the AtmiR173a and NbmiR482a target sites are in yellow and orange, respectively. The *AtTAS1c*-derived spacer and the syn-tasiRNA sequences are in black and blue, respectively.

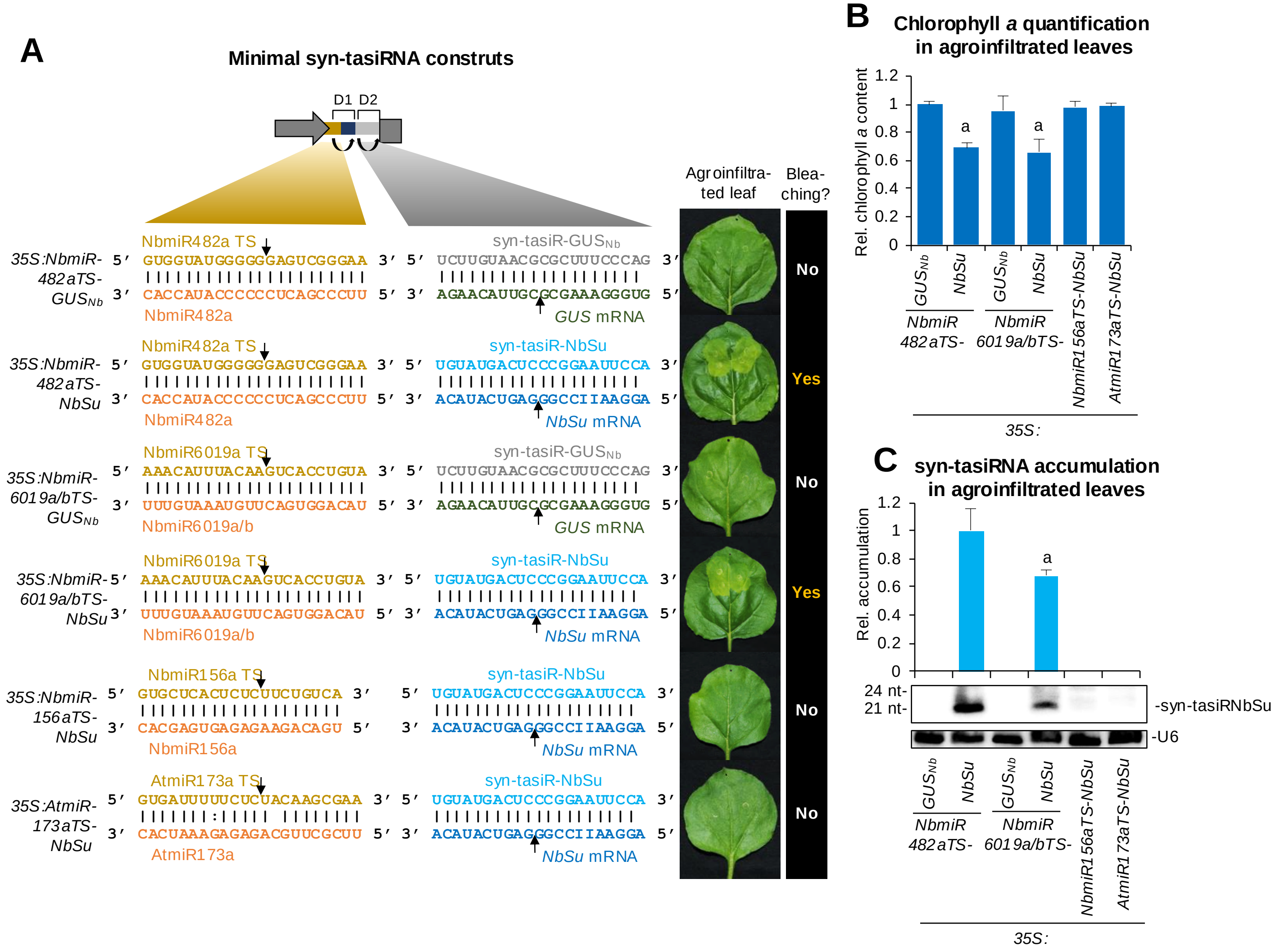


**Figure S3. Functional analysis of syn-tasiRNAs against *N. benthamiana SULPHUR* (*NbSu*) expressed from modified minimal precursors including different miRNA target sites (TS).** (**A**) Organization of minimal syn-tasiRNA constructs. Left, Minimal syn-tasiRNA constructs including 22-nt miRNA target sites from *N. benthamiana* (NbmiR482aTS and NbmiR6019TS) or *A. thaliana* (AtmiR173a), or the 21-nt TS from *N. benthamiana* miR156a (NbmiR156aTS). Nucleotides corresponding to miRNA TSs and miRNAs are shown in dark yellow and orange, respectively. Nucleotides corresponding to syn-tasiR-NbSu and target *NbSu* mRNA are shown in light and dark blue, respectively. Nucleotides corresponding to syn-tasiR-GUSNb and *GUS* mRNA are shown in grey and dark green, respectively. Arrows indicate the predicted cleavage sites for miRNAs and syn-tasiRNAs. Right: photographs at 7 days post agroinfiltration (dpa) of leaves agroinfiltrated with each of construct. The presence or absence of bleaching on the agroinfiltrated patches is labelled as "Yes" or "No", respectively. (**B**) Relative content of chlorophyll *a* in agroinfiltrated patches (*35S:NbmiR482aTS-GUS*$_{Nb}$ = 1.0). Bars with letter "a" are significantly different from the control sample (P < 0.05 in pairwise Student's t -test comparisons). (**C**) Target *NbSu* mRNA accumulation in agroinfiltrated leaves at 2 dpa [mean relative level (n = 3) + standard error] after normalization to *PROTEIN PHOSPHATASE 2A* (*NbPP2A*), as determined by quantitative RT-qPCR (*35S:NbmiR482aTS-GUS*$_{Nb}$ = 1). Other details are as in B.

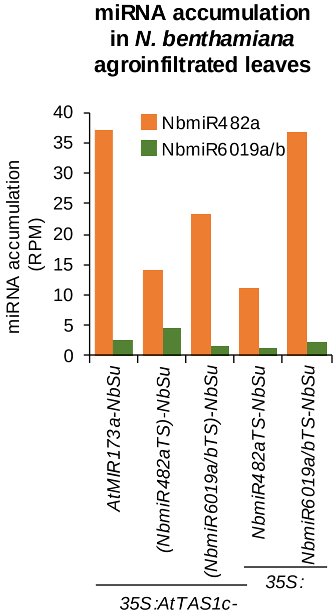


**Figure S4**. Analysis of NbmiR482a and NbmiR6019a/b presence in *Nicotiana benthamiana* tissues. Left, bar graph showing the accumulation (reads per million, RPM) of NmiR482a and NbmiR6019a/b revealed by high-throughput sequencing of small RNA libraries prepared from leaves agroinfiltrated with different constructs.

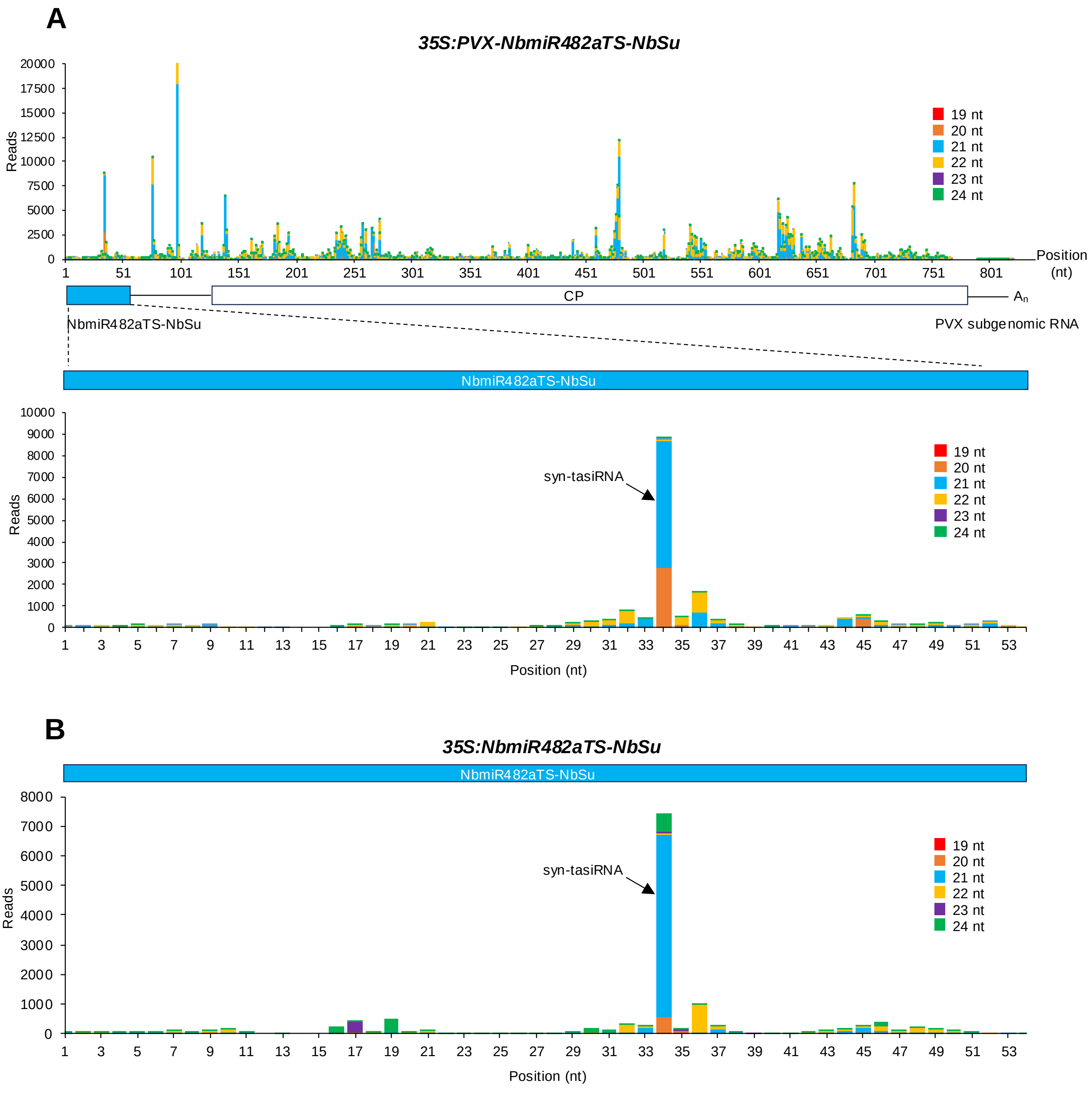


**Figure S5. Mapping of 19-24-nucleotide small RNA reads to *NbmiR482aTS-NbSu* precursors. A)** Top, mapping of reads to the whole subgenomic RNA sequence including PVX coat protein (CP). Bottom, mapping of reads exclusively to the *NbmiR482aTS-NbSu* precursor included in PVX. The *x*-axis indicates the position on the corresponding RNA sequence (subgenomic RNA or *NbmiR482aTS-NbSu* precursor in top and bottom graphs, respectively) in nucleotides of the 5' end of the sequence plotted. The *y*-axis is the small RNA coverage in total number of reads for each nucleotide position. B) Mapping of reads to the *NbmiR482TS-NbSu* precursor expressed from the *35S:NbmiR482aTS-NbSu* construct. Other details are as in A.

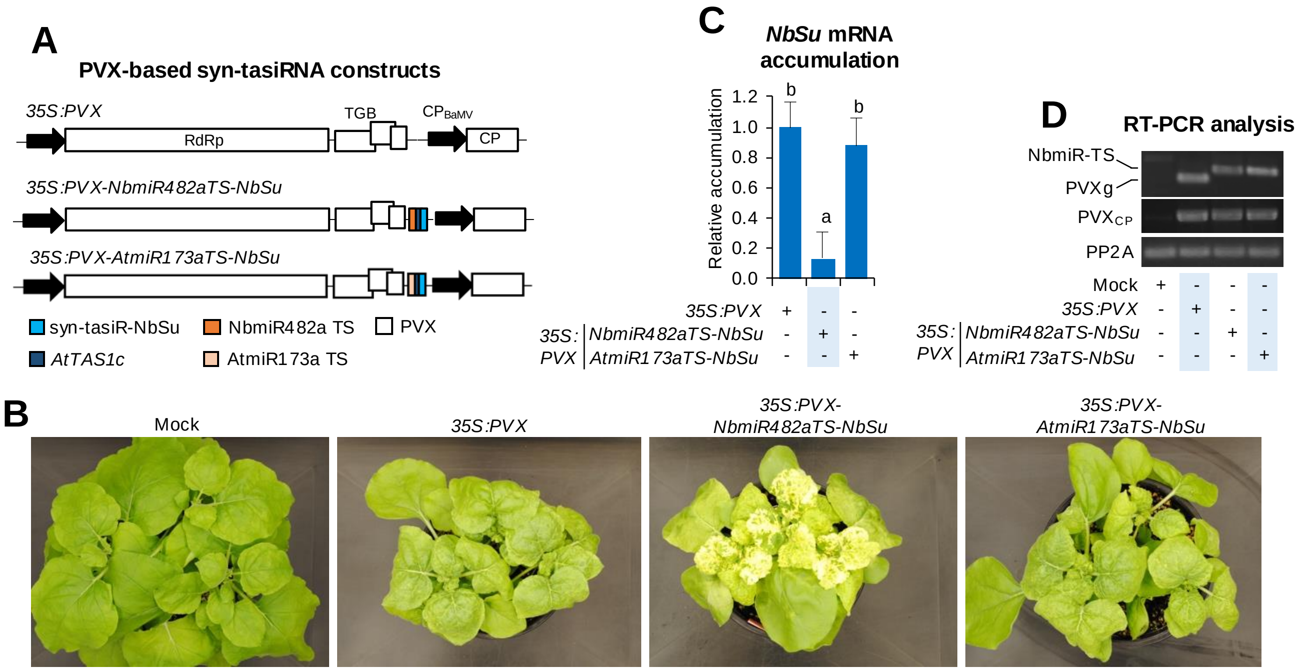


**Figure S6.** Functional analysis of potato virus X (PVX) constructs expressing syn-tasiR-NbSu from minimal precursors including endogenous or heterologous 22-nt miRNA target sites (TSs). (**A**) Diagram of PVX-based constructs. *AtTAS1c*, NbmiR482aTS, AtmiR173aTS and syn-tasiR-NbSu sequences are represented by dark blue, orange, light orange and light blue boxes, respectively. PVX ORFs and promoters are represented as white boxes and black arrows, respectively. RdRP, RNA-dependent RNA-polymerase; TGB, triple gene block; CP, coat protein; CPBaMV, Bamboo mosaic virus CP promoter. (**B**) Photos at 14 days post-agroinfiltration (dpa) of sets of three plants agroinoculated with the different constructs. (**C**) Target *NbSu* mRNA accumulation in RNA preparations from apical leaves collected at 14 days post-agroinfiltration (dpa) and analyzed individually (mock = 1.0 in all comparisons). Bars with the letter 'a' or 'b' indicate whether the mean values are significantly different from mock control or *35S:PVX-NbmiR482aTS-NbSu* samples, respectively ($P < 0.05$ in pairwise Student's *t*-test comparison). (**D**) RT-PCR detection of PVX and minimal precursors in apical leaves at 7 dpa. RT-PCR products corresponding to the *NbPP2A* and PVX vector controls are also shown (bottom). PVXg, band amplified from *35S:PVX* samples corresponding to the genomic region lacking a syn-tasiRNA precursor.

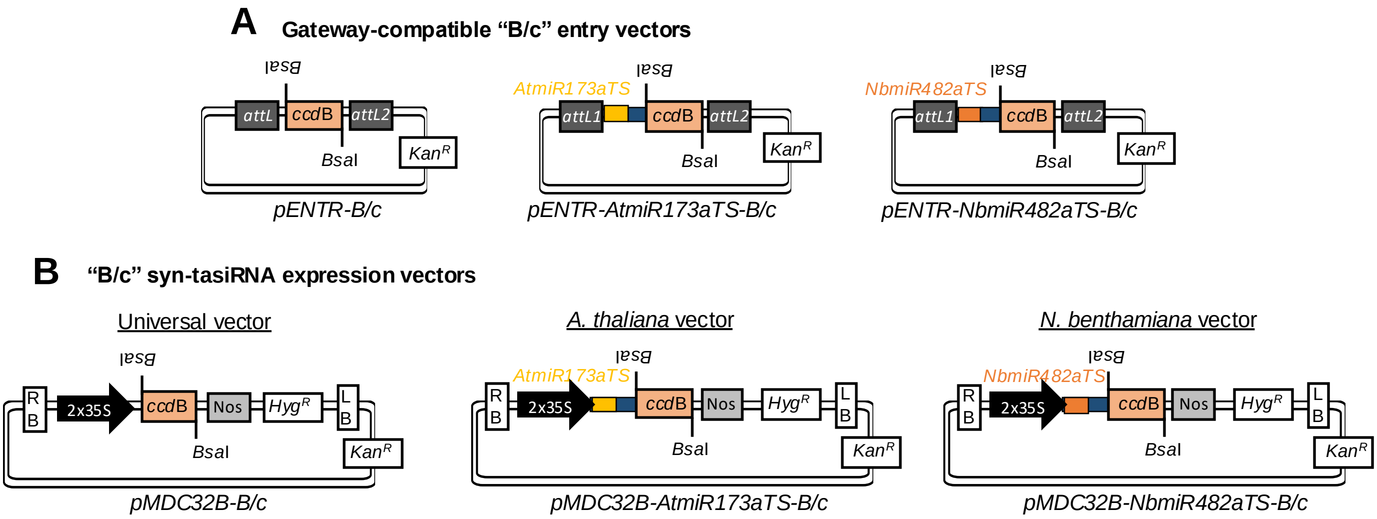


**Figure S7**. *B/c*-based vectors for direct cloning of syn-tasiRNAs downstream the AtmiR173a and NbmiR482a target sites (TSs), or the TS of choice. Sequences of AtmiR173a and NbmiR482a TSs are in yellow and orange, respectively. The spacer sequence derived from *AtTAS1c* is in blue. Top, diagram of the Gateway-compatible entry vectors. Bottom, diagram of binary vectors for in plant expression of amiRNAs. RB: right border; 35S: Cauliflower mosaic virus promoter; *Bsa*I: *Bsa*I recognition site, *ccd*B: gene encoding the gyrase toxin; LB: left border; attL1 and attL2: GATEWAY recombination sites. $Kan^R$: kanamycin resistance gene; $Hyg^R$: hygromycin resistance gene.

**Table S1.** Name, sequence and use of DNA oligonucleotides used in this study.

| Name | Sequence | Type* | Construct/Aim |
|---|---|---|---|
| AC-55 | AGGGGCCATGCTAATCTTCTC | ssDNA | Probe for U6 detection |
| AC-157 | GGCCTCTTCCTTTATAACCAA | ssDNA | Probe for syn-tasiR-AtFT detection |
| AC-158 | AGGGATTTCCGTGACACTTAA | ssDNA | Probe for amiR-AtCH42 detection |
| AC-159 | AAAAATGGCTGAGGCTGATGA | ssDNA | qPCR amplification of *AtACT2* mRNA |
| AC-160 | GAAAAACAGCCCTGGGAGC | ssDNA | |
| AC-163 | CATGCACAAGTAGGGACGGTT | ssDNA | qPCR amplification of *AtCH42* mRNA |
| AC-164 | GTCACGGAAATCCTTTGGGTT | ssDNA | |
| AC-169 | TGGAACAACCTTTGGCAATG | ssDNA | qPCR amplification of *AtFT* mRNA |
| AC-170 | CGACACGATGAATTCCTGCA | ssDNA | |
| AC-355 | GACCCTGATGTTGATGTTCGCT | ssDNA | qPCR amplification of *NbSu* mRNA |
| AC-356 | GAGGGATTTGAAGAGAGATTTC | ssDNA | |
| AC-365 | GACCCTGATGTTGATGTTCGCT | ssDNA | PCR&qPCR amplification of *NbPP2A* mRNA |
| AC-366 | GAGGGATTTGAAGAGAGATTTC | ssDNA | |
| AC-416 | A+GGA+CAC+AAT+CAC+GTC+TTA+CA | ssLNA | Probe for amiR-TSWV/syn-tasiR-TSWV-1 detection |
| AC-417 | G+CGG+GAA+GTC+CAC+CAC+GGT+TA | ssLNA | Probe for syn-tasiR-NbSu detection |
| AC-505 | CCCCATACCACAACAAAACAATGCTGTTTC ATCTCG | ssDNA | *35S:AtTAS1c(NbmiR482aTS)-NbSu* |
| AC-507 | TGTAAATGTTTAACAAAACAATGCTGTTTC ATCTCG | ssDNA | *35S:AtTAS1c(NbmiR6019a/bTS)-NbSu* |
| AC-631 | GGAGTCGGGAATAGACCATTTATGTATGAC TCCCGG | ssDNA | *35S:AtTAS1c(NbmiR482aTS)-NbSu* |
| AC-632 | AGTCACCTGTATAGACCATTTATGTATGACT CCCGG | ssDNA | *35S:AtTAS1c(NbmiR6019a/bTS)-NbSu* |
| AC-639 | CACCGTGGTATGGGGGGAGTCGGGAATAG ACCATTTATCTTGTAACGCGCTTTCCCAG | ssDNA | *35S:NbmiR482aTS-GUS*$_{Nb}$ |
| AC-640 | CTGGGAAAGCGCGTTACAAGATAAATGGTC TATTCCCGACTCCCCCCATACCACGGTG | ssDNA | |
| AC-641 | CACCGTGGTATGGGGGGAGTCGGGAATAG ACCATTTATGTATGACTCCCGGAATTCCA | ssDNA | *35S:NbmiR482aTS-NbSu* |
| AC-642 | TGGAATTCCGGGAGTCATACATAAATGGTC TATTCCCGACTCCCCCCATACCACGGTG | ssDNA | |
| AC-643 | CACCAAACATTTACAAGTCACCTGTATAGA CCATTTATCTTGTAACGCGCTTTCCCAG | ssDNA | *35S:NbmiR6019a/bTS-GUS*$_{Nb}$ |
| AC-644 | CTGGGAAAGCGCGTTACAAGATAAATGGTC TATACAGGTGACTTGTAAATGTTTGGTG | ssDNA | |
| AC-645 | CACCAAACATTTACAAGTCACCTGTATAGA CCATTTATGTATGACTCCCGGAATTCCA | ssDNA | *35S:NbmiR6019a/bTS-NbSu* |
| AC-646 | TGGAATTCCGGGAGTCATACATAAATGGTC TATACAGGTGACTTGTAAATGTTTGGTG | ssDNA | |
| AC-650 | GAGGTCAGCACCAGCTAGCAGTAGAGAAG AATCTGTA | ssDNA | *35S:PVX-amiR-GUS*$_{Nb}$, *35S:PVX-amiR-TSWV* |
| AC-654 | GGGAATCAATCACAGTGTTGGC | ssDNA | syn-tasiRNA precursors detection |
| AC-655 | GCTACTATGGCACGGGCTGTAC | ssDNA | |
| AC-657 | ATGTCAGGCCTGTTCACTATCC | ssDNA | PVX diagnostic |
| AC-658 | TGGTGGTGGTAGAGTGACAAC | ssDNA | |
| AC-663 | GGGAAACTTAACAAACCCTAGTAAGAAGA GCCAA | ssDNA | *35S:PVX-amiR-GUS*$_{Nb}$, *35S:PVX-amiR-TSWV* |
| AC-666 | AGAGGTCAGCACCAGCTAGCGTGGTATGGG GGGAGTCGGGAATAGACCATTTATGTATGA CTCCCGGAATTCCAAGGGTTTGTTAAGTTTC CCT | dsDNA | *35S:PVX-NbmiR482aTS-GUS*$_{Nb}$*(x4)* |
| AC-674 | CACCTGTAAGAGACCGGTCTCACATT | ssDNA | *pENTR-BB, pENTR-B/c pMDC32B-BB, pMDC32B-B/c* |
| AC-675 | AATGTGAGACCGGTCTCTTACAGGTG | ssDNA | |
| AC-676 | CACCGTGATTTTTCTCTACAAGCGAATAGA CCATTTATGCGCTTGCTGAGTTTCCCCC | ssDNA | *35S:AtmiR173aTS-GUS*$_{At}$ |
| AC-677 | GGGGGAAACTCAGCAAGCGCATAAATGGT CTATTCGCTTGTAGAGAAAAATCACGGTG | ssDNA | |

| | | | |
|---|---|---|---|
| AC-678 | CACCGTGATTTTTCTCTACAAGCGAATAGACCATTTATTAAGTGTCACGGAAATCCCT | ssDNA | *35S:AtmiR173aTS-AtCH42* |
| AC-679 | AGGGATTTCCGTGACACTTAATAAATGGTCTATTCGCTTGTAGAGAAAAATCACGGTG | ssDNA | |
| AC-680 | CACCGTGATTTTTCTCTACAAGCGAATAGACCATTTATTGGTTATAAAGGAAGAGGCC | ssDNA | *35S:AtmiR173aTS-AtFT* |
| AC-681 | GGCCTCTTCCTTTATAACCAATAAATGGTCTATTCGCTTGTAGAGAAAAATCACGGTG | ssDNA | |
| AC-682 | CACCGTGATTTTTCTCTACAAGCGAATAGACCATTTATTGGTTATAAAGGAAGAGGCCTCCCATTCGATACTGCTCGCC | ssDNA | *35S:AtmiR173aTS-AtFT-AtTrich* |
| AC-683 | GGCGAGCAGTATCGAATGGGAGGCCTCTTCCTTTATAACCAATAAATGGTCTATTCGCTTGTAGAGAAAAATCACGGTG | ssDNA | |
| AC-684 | CACCGTGATTTTTCTCTACAAGCGAATAGACCATTTATCCCATTCGATACTGCTCGCCTTGGTTATAAAGGAAGAGGCC | ssDNA | *35S:AtmiR173aTS-AtTrich-AtFT* |
| AC-685 | GGCCTCTTCCTTTATAACCAAGGCGAGCAGTATCGAATGGGATAAATGGTCTATTCGCTTGTAGAGAAAAATCACGGTG | ssDNA | |
| AC-712 | AGAGGTCAGCACCAGCTAGCAAACCTAAACCTAAACGG | ssDNA | *35S:PVX-AtTAS1c(NbmiR482aTS)-NbSu* |
| AC-713 | AGGGAAACTTAACAAACCCTATTTCACTTTACGATGTGG | ssDNA | |
| AC-716 | TGTAGTGGTATGGGGGGAGTCGGGAATAGACCATTTATGTATGACTCCCGGAATTCCA | ssDNA | *35S:NbmiR482aTS-NbSu* |
| AC-717 | AATGTGGAATTCCGGGAGTCATACATAAATGGTCTATTCCCGACTCCCCCCATACCAC | ssDNA | |
| AC-719 | TGTAGTGCTCACTCTCTTCTGTCATAGACCATTTATGTATGACTCCCGGAATTCCA | ssDNA | *35S:NbmiR156aTS-NbSu* |
| AC-720 | AATGTGGAATTCCGGGAGTCATACATAAATGGTCTATGACAGAAGAGAGTGAGCAC | ssDNA | |
| AC-721 | TGTAGTGATTTTTCTCTACAAGCGAATAGACCATTTATGTATGACTCCCGGAATTCCA | ssDNA | *35S:AtmiR173aTS-NbSu* |
| AC-722 | AATGTGGAATTCCGGGAGTCATACATAAATGGTCTATTCGCTTGTAGAGAAAAATCAC | ssDNA | |
| AC-900 | TGTAGTGGTATGGGGGGAGTCGGGAATAGACCATTTAAGAGACCGGTCTCATCGG | ssDNA | *pENTR-NbmiR482aTS-B/c*<br>*pMDC32B-NbmiR482aTS-B/c* |
| AC-901 | AATGCCGATGAGACCGGTCTCTTAAATGGTCTATTCCCGACTCCCCCCATACCAC | ssDNA | |
| AC-902 | TGTAGGCATGGGCGGTGTAGGCAAGATAGACCATTTAAGAGACCGGTCTCATCGG | ssDNA | *pENTR-NbmiR6019a/bTS-B/c*<br>*pMDC32B-NbmiR6019a/bTS-B/c* |
| AC-903 | AATGCCGATGAGACCGGTCTCTTAAATGGTCTATCTTGCCTACACCGCCCATGCC | ssDNA | |
| AC-935 | TTTATGTAAGACGTGATTGTGTCCTTATCAGCTCTGGGTGAATCGGTTGGTATAGTGGGGCATACCGTCAGAGTGCACAATCCATCTT | ssDNA | *35S:NbmiR482aTS-TSWV(x4)* |
| AC-936 | CCGAAAGATGGATTGTGCACTCTGACGGTATGCCCCACTATACCAACCGATTCACCCAGAGCTGATAAGGACACAATCACGTCTTACA | ssDNA | |
| AC-987 | AGAGGTCAGCACCAGCTAGCGTGGTATGGGGGGAGTCGGGAATAGACCATTTATATTGACCCACACTTTGCCGATAACCTTCACCCGGTTGCCACTATTGACCCACACTTTGCCGATAACCTTCACCCGGTTGCCACAGGGTTTGTTAAGTTTCCCT | dsDNA | *35S:PVX-NbmiR482aTS-GUS$_{Nb}$(x4)* |
| AC-988 | AGAGGTCAGCACCAGCTAGCGTGGTATGGGGGGAGTCGGGAA | ssDNA | *35S:PVX-NbmiR482aTS-TSWV(x4)* |
| AC-989 | AGGGAAACTTAACAAACCCTAAGATGGATTGTGCACTCTGA | ssDNA | |
| AC-1093 | TGTAGTGATTTTTCTCTACAAGCGAATAGACCATTTAAGAGACCGGTCTCATCGG | ssDNA | *pENTR-AtmiR173aTS-B/c*<br>*pMDC32B-AtmiR173aTS-B/c* |
| AC-1094 | AATGCCGATGAGACCGGTCTCTTAAATGGTCTATTCGCTTGTAGAGAAAAATCAC | ssDNA | |

*ssDNA: single-stranded DNA; dsDNA: double-stranded DNA; LNA: locked nucleic acid.

**Table S2:** Phenotypic penetrance of syn-tasiRNAs expressed in *A. thaliana* Col-0 T1 transgenic plants for silencing *FT*.

| Construct | T1 analyzed | Phenotypic penetrance[a] |
|---|---|---|
| *35S:AtTAS1c-GUS*$_{At}$ | 58 | 0% |
| *35S:AtTAS1c-AtFT* | 65 | 100% |
| *35S:AtmiR173aTS-AtFT* | 53 | 0% |
| *35S:AtmiR173aTS- GUS*$_{At}$ | 64 | 100% |

[a] The FT phenotype was defined as a higher 'days to flowering' value when compared to the average 'days to flowering' value of the *35S:AtTAS1c-GUS*$_{At}$ and *35S:AtmiR173aTS-GUS*$_{At}$ control sets.

**Table S3:** Phenotypic penetrance of syn-tasiRNAs expressed in *A. thaliana* Col-0 T1 transgenic plants for silencing *CH42*.

| **Construct** | **T1 analyzed** | **Phenotypic penetrance**[a] |
|---|---|---|
| *35S:AtTAS1c-GUS*$_{At}$ | 70 | 0% |
| *35S:AtTAS1c-AtCH42* | 402 | 78%<br>17.9% weak<br>23.6% intermediate<br>36.1 % severe |
| *35S:AtmiR173aTS-GUS*$_{At}$ | 134 | 0% |
| *35S:AtmiR173aTS-AtCH42* | 389 | 51%<br>7.2% weak<br>19.3% intermediate<br>24.7 % severe |

[a] Ch42 phenotype is scored in 10 days-old seedling and is considered 'weak', 'intermediate' or 'severe' if seedlings have >2 leaves, exactly 2 leaves or no leaves (only 2 cotyledons), respectively.

**Table S4:** Phenotypic penetrance of syn-tasiRNAs expressed in Arabidopsis Col-0 T1 transgenic plants for silencing *FT* and *TRY*.

| **Construct** | **T1 analyzed** | **Phenotypic penetrance**[a] |
|---|---|---|
| *35S:AtTAS1c-GUS*$_{At}$ | 46 | 0% FT<br>0% TRY |
| *35S:AtTAS1c-AtFT-AtTRY* | 16 | 100% FT<br>88% TRY |
| *35S:AtTAS1c-AtTRY-AtFT* | 16 | 100% FT<br>82% TRY |
| *35S:AtmiR173TS-GUS*$_{At}$ | 27 | 0% FT<br>0% TRY |
| *35S:AtmiR173TS-AtFT-AtTRY* | 34 | 100% FT<br>80% TRY |
| *35S:AtmiR173TS-AtTRY-AtFT* | 20 | 100% FT<br>72% TRY |

[a] The FT phenotype was defined as a higher ‘days to flowering’ value when compared to the average ‘days to flowering’ value of the *35S:AtTAS1c-GUS*$_{At}$ control set.

The TRY phenotype was defined as a higher number of trichomes when compared to transformants of the *35S:AtTAS1c-GUS*$_{At}$ control set.

**Text S1.** Protocol to design and clone syn-tasiRNAs downstream the 3'D1[+] position in *Bsa*I/*ccd*B-based ('B/c') vectors *pENTR-B/c*, *pMDC32-B/c*, *pENTR-AtmiR173aTS-B/c*, *pMDC32B-AtmiR173aTS-B/c*, *pENTR-NbmiR482aTS-B/c* and *pMDC32B-NbmiR482aTS-B/c*.

### 1. Selection of the syn-tasiRNA sequence(s)

Use the Syn-tasiRNA Designer app from the P-SAMS webtool at http://p-sams.carringtonlab.org/syntasi/designer.

### 2. Design of syn-tasiRNA oligonucleotides for cloning

Next are described some designs for cloning two syn-tasiRNAs in tandem downstream the 3'D1[+] position.

2.1. Using your 22-nt miRNA target site of choice:

Use vectors *pENTR-B/c* or *pMDC32-B/c* and order the following oligos

-Forward oligonucleotide (79 b):

**TGTA**$X_1X_2X_3X_4X_5X_6X_7X_8X_9X_{10}X_{11}X_{12}X_{13}X_{14}X_{15}X_{16}X_{17}X_{18}X_{19}X_{20}X_{21}X_{22}$TAGACCATTTA$X_1X_2X_3X_4X_5X_6X_7X_8X_9X_{10}X_{11}X_{12}X_{13}X_{14}X_{15}X_{16}X_{17}X_{18}X_{19}X_{20}X_{21}X_1X_2X_3X_4X_5X_6X_7X_8X_9X_{10}X_{11}X_{12}X_{13}X_{14}X_{15}X_{16}X_{17}X_{18}X_{19}X_{20}X_{21}$

-Reverse oligonucleotide (79 b):

**AATG**$Y_{21}Y_{20}Y_{19}Y_{18}Y_{17}Y_{16}Y_{15}Y_{14}Y_{13}Y_{12}Y_{11}Y_{10}Y_9Y_8Y_7Y_6Y_5Y_4Y_3Y_2Y_1Y_{21}Y_{20}Y_{19}Y_{18}Y_{17}Y_{16}Y_{15}Y_{14}Y_{13}Y_{12}Y_{11}Y_{10}Y_9Y_8Y_7Y_6Y_5Y_4Y_3Y_2Y_1$TAAATGGTCTA$Y_{22}Y_{21}Y_{20}Y_{19}Y_{18}Y_{17}Y_{16}Y_{15}Y_{14}Y_{13}Y_{12}Y_{11}Y_{10}Y_9Y_8Y_7Y_6Y_5Y_4Y_3Y_2Y_1$

Where:

TAGACCATTTA=*AtTAS1c*-derived spacer sequence

$X_1X_2X_3X_4X_5X_6X_7X_8X_9X_{10}X_{11}X_{12}X_{13}X_{14}X_{15}X_{16}X_{17}X_{18}X_{19}X_{20}X_{21}X_{22}$=22-nt miRNA target site sequence

$X_1X_2X_3X_4X_5X_6X_7X_8X_9X_{10}X_{11}X_{12}X_{13}X_{14}X_{15}X_{16}X_{17}X_{18}X_{19}X_{20}X_{21}$=syn-tasiRNA-1 sequence

$X_1X_2X_3X_4X_5X_6X_7X_8X_9X_{10}X_{11}X_{12}X_{13}X_{14}X_{15}X_{16}X_{17}X_{18}X_{19}X_{20}X_{21}$=syn-tasiRNA-2 sequence

$Y_{21}Y_{20}Y_{19}Y_{18}Y_{17}Y_{16}Y_{15}Y_{14}Y_{13}Y_{12}Y_{11}Y_{10}Y_9Y_8Y_7Y_6Y_5Y_4Y_3Y_2Y_1$=syn-tasiRNA-1 reverse-complement sequence

$Y_{21}Y_{20}Y_{19}Y_{18}Y_{17}Y_{16}Y_{15}Y_{14}Y_{13}Y_{12}Y_{11}Y_{10}Y_9Y_8Y_7Y_6Y_5Y_4Y_3Y_2Y_1$=syn-tasiRNA-2 reverse-complement sequence

TAAATGGTCTA=*AtTAS1c*-derived reverse-complement spacer sequence

$Y_{22}Y_{21}Y_{20}Y_{19}Y_{18}Y_{17}Y_{16}Y_{15}Y_{14}Y_{13}Y_{12}Y_{11}Y_{10}Y_9Y_8Y_7Y_6Y_5Y_4Y_3Y_2Y_1$=22-nt miRNA target site reverse-complement sequence

**Example**

The sequences of the two oligonucleotides to clone syn-tasiRNAs ‘syn-tasiR-TRY’ (**TCCCATTCGATACTGCTCGCC**) and ‘syn-tasiR-Ft’ (**TTGGTTATAAAGGAAGAGGCC**) in positions 3’D2[+] and 3’D3[+], respectively, of a minimal precursors including the *AtmiR173aTS* (**GTGATTTTTCTCTACAAGCGAA**) are:

-<u>Forward oligonucleotide</u> (79b):

**TGTAGTGATTTTTCTCTACAAGCGAA**TAGACCATTTA**TCCCATTCGATACTGCTCGCCTTGGTTATAA AGGAAGAGGCC**

-<u>Reverse oligonucleotide</u> (79 b):

**AATGGGCCTCTTCCTTTATAACCAAGGCGAGCAGTATCGAATGGGA**TAAATGGTCTA**TTCGCTTGTAG AGAAAAATCAC**

<u>2.2. Using AtmiR173a or NbmiR482a miRNAs target site:</u>

Use vectors *pENTR-AtmiR173aTS-B/c*, *pMDC32-AtmiR173aTS-B/c*, *pENTR-NbmiR482aTS-B/c* or *pMDC32-AtmiR482aTS-B/c* and order the following oligos:

-<u>Forward oligonucleotide</u> (46 b):

**TTTA**$X_1X_2X_3X_4X_5X_6X_7X_8X_9X_{10}X_{11}X_{12}X_{13}X_{14}X_{15}X_{16}X_{17}X_{18}X_{19}X_{20}X_{21}X_1X_2X_3X_4X_5X_6X_7X_8X_9X_{10}X_{11}X_{12}$ $X_{13}X_{14}X_{15}X_{16}X_{17}X_{18}X_{19}X_{20}X_{21}$

-<u>Reverse oligonucleotide</u> (46 b):

**CCGA**$Y_{21}Y_{20}Y_{19}Y_{18}Y_{17}Y_{16}Y_{15}Y_{14}Y_{13}Y_{12}Y_{11}Y_{10}Y_9Y_8Y_7Y_6Y_5Y_4Y_3Y_2Y_1Y_{21}Y_{20}Y_{19}Y_{18}Y_{17}$ $Y_{16}Y_{15}Y_{14}Y_{13}Y_{12}Y_{11}Y_{10}Y_9Y_8Y_7Y_6Y_5Y_4Y_3Y_2Y_1$

Where:

$X_1X_2X_3X_4X_5X_6X_7X_8X_9X_{10}X_{11}X_{12}X_{13}X_{14}X_{15}X_{16}X_{17}X_{18}X_{19}X_{20}X_{21}$=syn-tasiRNA-1 sequence

$X_1X_2X_3X_4X_5X_6X_7X_8X_9X_{10}X_{11}X_{12}X_{13}X_{14}X_{15}X_{16}X_{17}X_{18}X_{19}X_{20}X_{21}$=syn-tasiRNA-2 sequence

$Y_{21}Y_{20}Y_{19}Y_{18}Y_{17}Y_{16}Y_{15}Y_{14}Y_{13}Y_{12}Y_{11}Y_{10}Y_9Y_8Y_7Y_6Y_5Y_4Y_3Y_2Y_1$=syn-tasiRNA-1 reverse-complement sequence

$Y_{21}Y_{20}Y_{19}Y_{18}Y_{17}Y_{16}Y_{15}Y_{14}Y_{13}Y_{12}Y_{11}Y_{10}Y_9Y_8Y_7Y_6Y_5Y_4Y_3Y_2Y_1$=syn-tasiRNA-2 reverse-complement sequence

**Example**

The sequences of the two oligonucleotides to clone syn-tasiRNAs ‘syn-tasiR-TRY’ (**TCCCATTCGATACTGCTCGCC**) and ‘syn-tasiR-Ft’ (**TTGGTTATAAAGGAAGAGGCC**) in positions 3’D2[+] and 3’D3[+], respectively, of minimal precursors included in AtmiR173aTS- or NbmiR482aTS-based B/c vectors are:

-<u>Forward oligonucleotide</u> (46 b):

**TTTATCCCATTCGATACTGCTCGCCTTGGTTATAAAGGAAGAGGCC**

-<u>Reverse oligonucleotide</u> (46 b):

**CCGAGGCCTCTTCCTTTATAACCAAGGCGAGCAGTATCGAATGGGA**

#### 3. Cloning of the syn-tasiRNA sequence(s) in B/c-based vectors

*Notes:*

*-New available -B/c vectors are listed in Table I at the end of the section.*

-B/c-*based vectors must be propagated in a* ccd*B resistant* E. coli *strain such as DB3.1.*

-Alternatively, Bsa*I digestion of the B/c vector and subsequent ligation of the amiRNA oligonucleotide insert can be done in separate reactions*

<u>3.1. Oligonucleotide annealing</u>

-Dilute sense oligonucleotide and antisense oligonucleotide in sterile $H_2O$ to a final concentration of 100 µM.

-Prepare Oligo Annealing Buffer:

60 mM Tris-HCl (pH 7.5)

500 mM NaCl

60 mM $MgCl_2$

10 mM DTT

***Note:*** *Prepare 1 ml aliquots of Oligo Annealing Buffer and store at -20$^0$C.*

-Assemble the annealing reaction in a PCR tube as described below:

| | |
|---|---|
| Forward oligonucleotide (100 µM) | 2 µL |
| Reverse oligonucleotide (100 µM) | 2 µL |
| Oligo Annealing Buffer | 46 µL |
| Total volume | 50 µL |

The final concentration of each oligonucleotide is 4 µM.

-Use a thermocycler to heat the annealing reaction 5 min at 94°C and then cool down (0.05ºC/sec) to 20°C.

-Dilute the annealed oligonucleotides just prior to assembling the digestion-ligation reaction as described below:

| | |
|---|---|
| Annealed oligonucleotides | 3 µL |
| $dH_2O$ | 37 µL |
| Total volume | 40 µL |

The final concentration of each oligonucleotide is 0.15 µM.

***Note**: Do not store the diluted oligonucleotides.*

3.2. Digestion-ligation reaction

- Assemble the digestion-ligation reaction as described below:

| | |
|---|---|
| B/c vector (x ug/uL) | Y µL (50 ng) |
| Diluted annealed oligonucleotides | 1 µL |
| 10x T4 DNA ligase buffer | 1 µL |
| T4 DNA ligase (400 U/µL) | 1 µL |
| *Bsa*I (10U/ µL, NEB) | 1 µL |
| $dH_2O$ | to 10 µL |
| Total volume | 10 µL |

Prepare a negative control reaction lacking *Bsa*I.

-Mix the reactions by pipetting. Incubate the reactions at room temperature for 5 minutes at 37°C.

3.3. *E.coli* transformation and analysis of transformants

-Transform 1-5 ul of the digestion-ligation reaction into an *E. coli* strain that doesn't have *ccd*B resistance (e.g. DH10B, TOP10, …) to do counter-selection.

-Pick two colonies/construct, grow LB-Kan (100 mg/ml) cultures and purify plasmids.

-Sequence with appropriate primers: M13-F (CCCAGTCACGACGTTGTAAAACGACGG) and

M13-R (CAGAGCTGCCAGGAAACAGCTATGACC) for *pENTR*-based vectors; attB1 (ACAAGTTTGTACAAAAAAGCAGGCT) and attB2 (ACCACTTTGTACAAGAAAGCTGGGT) primers for *pMDC32B*-based vectors).

**Table I**: *Bsa*I/*ccd*B-based ('B/c') vectors for direct cloning of syn-tasiRNAs downstream position 3'D1[+] in minimal precursor.

| Vector | Small RNA expressed | Bacterial antibiotic resistance | Plant antibiotic resistance | GATEWAY use | Backbone | Promoter of syn-tasiRNA cassette | Terminator of syn-tasiRNA cassette | Plant species tested |
|---|---|---|---|---|---|---|---|---|
| *pENTR-B/c* | – | Kanamycin | – | Donor | *pENTR* | – | – | – |
| *pMDC32B-B/c* | | Kanamycin<br>Hygromycin | Hygromycin | – | *pMDC32* | *CaMV* 2x35S | *Nos* | *A. thaliana*<br>*N. benthamiana* |
| *pENTR-AtmiR173aTS-B/c* | syn-tasiRNAs | Kanamycin | – | Donor | *pENTR* | – | – | – |
| *pMDC32B-AtmiR173aTS-B/c* | syn-tasiRNAs | Kanamycin<br>Hygromycin | Hygromycin | – | *pMDC32* | *CaMV* 2x35S | *Nos* | *A. thaliana* |
| *pENTR-NbmiR482aTS-B/c* | syn-tasiRNAs | Kanamycin | – | Donor | *pENTR* | – | – | – |
| *pMDC32B-NbmiR482aTS-B/c* | syn-tasiRNAs | Kanamycin<br>Hygromycin | Hygromycin | – | *pMDC32* | *CaMV* 2x35S | *Nos* | *N. benthamiana* |

**Text S2.** Protocol to generate PVX-based syn-tasiRNA constructs.

### 1. Preparation of the dsDNA syn-tasiRNA insert

Design and order a dsDNA (eg. ultramer duplex in IDT) including the sequences of your syn-tasiRNA(s) (2 in the following example) following the 22-nt miRNA target site of interest, as follows:

agaggtcagcaccagctagc$X_1X_2X_3X_4X_5X_6X_7X_8X_9X_{10}X_{11}X_{12}X_{13}X_{14}X_{15}X_{16}X_{17}X_{18}X_{19}X_{20}X_{21}X_{22}$TAGACCATTTA$X_1X_2X_3X_4X_5X_6X_7X_8X_9X_{10}X_{11}X_{12}X_{13}X_{14}X_{15}X_{16}X_{17}X_{18}X_{19}X_{20}X_{21}X_1X_2X_3X_4X_5X_6X_7X_8X_9X_{10}X_{11}X_{12}X_{13}X_{14}X_{15}X_{16}X_{17}X_{18}X_{19}X_{20}X_{21}$agggtttgttaagtttccct

Where:

- -X is a DNA base of the 22-nt miRNA target site sequence, and the subscript number is the base position
- -X is a DNA base of the syn-tasiRNA-1 sequence, and the subscript number is the base position in the syn-tasiRNA* 21-mer
- -X is a DNA base of the syn-tasiRNA-2 sequence, and the subscript number is the base position in the syn-tasiRNA 21-mer
- -x is a DNA base of the PVX sequence, required for Gibson-based assembly
- -X is a DNA base of the *AtTAS1c* sequence

Note that:

- -In general, **$X_1$=T** and **$X_1$=T** for amiRNA association with AGO1.

Fragment #1 (syn-tasiRNA precursor) is ready.

### 2. Preparation of the vector

-Digest *pLB-PVX* with *Mlu*I.

-Gel purify the 9921 bp band corresonding to linearized plasmid.

-Quantify 1 ul in Nanodrop.

Fragment #2 (backbone vector) is ready.

### 3. Assembly

-Assemble the Gibbson reaction as described below:

| | |
|---|---|
| Fragment 1 (dsDNA insert)[a] | |
| Fragment 2 (vector)[b,c,d] | |
| GeneArt Gibson Assembly HiFI Master Mix | 5 µL |
| $dH_2O$ | to 10 µL |

Total volume 10 µL

[a]The optimal amount of vector is between 50-100 ng

[b]Insert/vector molar excess is between 2-3.

[c]Total DNA amount is between 0.02-0.5 pmol

[d]Mass to moles conversions can be calculated here: http://nebiocalculator.neb.com/#!/ssdnaamt

-Incubate reactions at 50ºC for 1h.

-Clean up reactions with a column (e.g. Zymo Research)

-Transform 1-4 µL in *E. coli* DH5α

-Plate in L-Kan plates and incubate 16h at 37ºC

### 4. Clone verification

-Pick several colonies and grow in liquid LB-Kan 16h at 37ºC, and purify plasmids.

-Digest candidate clones with *Apa*I+*Xho*I

Good clones: 8595 + **1409** bp

Bad clones (empty *pLB-PVX*): 9921 bp + **1738** bp

-Confirm insert sequence by Sanger sequencing with forward and reverse oligos AC-654 (`GGGAATCAATCACAGTGTTGGC`) and/or AC-655 (`GCTACTATGGCACGGGCTGTAC`), respectively.

**Text S3**. DNA sequence in FASTA format of all precursors used to express art-sRNAs in plants.

1. *AtTAS1c*-based precursors

***>AtTAS1c-GUS$_{At}$***

```
AAACCTAAACCTAAACGGCTAAGCCCGACGTCAAATACCAAAAAGAGAAAAACAAGAGCGCCGTCAAGCTCTGCAAATACGATCTGTAAG
TCCATCTTAACACAAAAGTGAGATGGGTTCTTAGATCATGTTCCGCCGTTAGATCGAGTCATGGTCTTGTCTCATAGAAAGGTACTTTCG
TTTACTTCTTTTGAGTATCGAGTAGAGCGTCGTCTATAGTTAGTTTGAGATTGCGTTTGTCAGAAGTTAGGTTCAATGTCCCGGTCCAAT
TTTCACCAGCCATGTGTCAGTTTCGTTCCTTCCCGTCCTCTTCTTTGATTTCGTTGGGTTACGGATGTTTTCGAGATGAAACAGCATTGT
TTTGTTGTGATTTTTCTCTACAAGCGAATAGACCATTTATGCGCTTGCTGAGTTTCCCCCTCGGTGGATCTTAGAAAATTATTCTAAGTC
CAACATAGCGTATTCTAAGTTCAACATATCGACGAACTAGAAAAGACATTGGACATATTCCAGGATATGCAAAAGAAAACAATGAATATT
GTTTTGAATGTGTTCAAGTAAATGAGATTTTCAAGTCGTCTAAAGAACAGTTGCTAATACAGTTACTTATTTCAATAAATAATTGGTTCT
AATAATACAAAACATATTCGAGGATATGCAGAAAAAAAGATGTTTGTTATTTTGAAAAGCTTGAGTAGTTTCTCTCCGAGGTGTAGCGAA
GAAGCATCATCTACTTTGTAATGTAATTTTCTTTATGTTTTCACTTTGTAATTTTATTTGTGTTAATGTACCATGGCCGATATCGGTTTT
ATTGAAAGAAAATTTATGTTACTTCTGTTTTGGCTTTGCAATCAGTTATGCTAGTTTTCTTATACCCTTTCGTAAGCTTCCTAAGGAATC
GTTCATTGATTTCCACTGCTTCATTGTATATTAAAACTTTACAACTGTATCGACCATCATATAATTCTGGGTCAAGAGATGAAAATAGAA
CACCACATCGTAAAGTGAAAT
```

*AtTAS1c*
AtmiR173a TS
syn-tasiR-GUS$_{At}$

***>AtTAS1c-AtFT***

```
AAACCTAAACCTAAACGGCTAAGCCCGACGTCAAATACCAAAAAGAGAAAAACAAGAGCGCCGTCAAGCTCTGCAAATACGATCTGTAAG
TCCATCTTAACACAAAAGTGAGATGGGTTCTTAGATCATGTTCCGCCGTTAGATCGAGTCATGGTCTTGTCTCATAGAAAGGTACTTTCG
TTTACTTCTTTTGAGTATCGAGTAGAGCGTCGTCTATAGTTAGTTTGAGATTGCGTTTGTCAGAAGTTAGGTTCAATGTCCCGGTCCAAT
TTTCACCAGCCATGTGTCAGTTTCGTTCCTTCCCGTCCTCTTCTTTGATTTCGTTGGGTTACGGATGTTTTCGAGATGAAACAGCATTGT
TTTGTTGTGATTTTTCTCTACAAGCGAATAGACCATTTATTGGTTATAAAGGAAGAGGCCTCGGTGGATCTTAGAAAATTATTCTAAGTC
CAACATAGCGTATTCTAAGTTCAACATATCGACGAACTAGAAAAGACATTGGACATATTCCAGGATATGCAAAAGAAAACAATGAATATT
GTTTTGAATGTGTTCAAGTAAATGAGATTTTCAAGTCGTCTAAAGAACAGTTGCTAATACAGTTACTTATTTCAATAAATAATTGGTTCT
AATAATACAAAACATATTCGAGGATATGCAGAAAAAAAGATGTTTGTTATTTTGAAAAGCTTGAGTAGTTTCTCTCCGAGGTGTAGCGAA
GAAGCATCATCTACTTTGTAATGTAATTTTCTTTATGTTTTCACTTTGTAATTTTATTTGTGTTAATGTACCATGGCCGATATCGGTTTT
ATTGAAAGAAAATTTATGTTACTTCTGTTTTGGCTTTGCAATCAGTTATGCTAGTTTTCTTATACCCTTTCGTAAGCTTCCTAAGGAATC
GTTCATTGATTTCCACTGCTTCATTGTATATTAAAACTTTACAACTGTATCGACCATCATATAATTCTGGGTCAAGAGATGAAAATAGAA
CACCACATCGTAAAGTGAAAT
```

*AtTAS1c*
AtmiR173a TS
syn-tasiR-AtFT

***>AtTAS1c-D2-AtCH42***

```
AAACCTAAACCTAAACGGCTAAGCCCGACGTCAAATACCAAAAAGAGAAAAACAAGAGCGCCGTCAAGCTCTGCAAATACGATCTGTAAG
TCCATCTTAACACAAAAGTGAGATGGGTTCTTAGATCATGTTCCGCCGTTAGATCGAGTCATGGTCTTGTCTCATAGAAAGGTACTTTCG
TTTACTTCTTTTGAGTATCGAGTAGAGCGTCGTCTATAGTTAGTTTGAGATTGCGTTTGTCAGAAGTTAGGTTCAATGTCCCGGTCCAAT
TTTCACCAGCCATGTGTCAGTTTCGTTCCTTCCCGTCCTCTTCTTTGATTTCGTTGGGTTACGGATGTTTTCGAGATGAAACAGCATTGT
TTTGTTGTGATTTTTCTCTACAAGCGAATAGACCATTTATTAAGTGTCACGGAAATCCCTTCGGTGGATCTTAGAAAATTATTCTAAGTC
CAACATAGCGTATTCTAAGTTCAACATATCGACGAACTAGAAAAGACATTGGACATATTCCAGGATATGCAAAAGAAAACAATGAATATT
GTTTTGAATGTGTTCAAGTAAATGAGATTTTCAAGTCGTCTAAAGAACAGTTGCTAATACAGTTACTTATTTCAATAAATAATTGGTTCT
AATAATACAAAACATATTCGAGGATATGCAGAAAAAAAGATGTTTGTTATTTTGAAAAGCTTGAGTAGTTTCTCTCCGAGGTGTAGCGAA
GAAGCATCATCTACTTTGTAATGTAATTTTCTTTATGTTTTCACTTTGTAATTTTATTTGTGTTAATGTACCATGGCCGATATCGGTTTT
ATTGAAAGAAAATTTATGTTACTTCTGTTTTGGCTTTGCAATCAGTTATGCTAGTTTTCTTATACCCTTTCGTAAGCTTCCTAAGGAATC
GTTCATTGATTTCCACTGCTTCATTGTATATTAAAACTTTACAACTGTATCGACCATCATATAATTCTGGGTCAAGAGATGAAAATAGAA
CACCACATCGTAAAGTGAAAT
```

*AtTAS1c*
AtmiR173a TS
syn-tasiR-AtCH42

***>AtTAS1c-AtFT-AtTry***

AAACCTAAACCTAAACGGCTAAGCCCGACGTCAAATACCAAAAAGAGAAAAACAAGAGCGCCGTCAAGCTCTGCAAATACGATCTGTAAG
TCCATCTTAACACAAAAGTGAGATGGGTTCTTAGATCATGTTCCGCCGTTAGATCGAGTCATGGTCTTGTCTCATAGAAAGGTACTTTCG
TTTACTTCTTTTGAGTATCGAGTAGAGCGTCGTCTATAGTTAGTTTGAGATTGCGTTTGTCAGAAGTTAGGTTCAATGTCCCGGTCCAAT
TTTCACCAGCCATGTGTCAGTTTCGTTCCTTCCCGTCCTCTTCTTTGATTTCGTTGGGTTACGGATGTTTTCGAGATGAAACAGCATTGT
TTTGTTGTGATTTTTCTCTACAAGCGAATAGACCATTTATTGGTTATAAAGGAAGAGGCCTCCCATTCGATACTGCTCGCCTCGGTGGAT
CTTAGAAAATTATTCTAAGTCCAACATAGCGTATTCTAAGTTCAACATATCGACGAACTAGAAAAGACATTGGACATATTCCAGG
ATATGCAAAAGAAAACAATGAATATTGTTTTGAATGTGTTCAAGTAAATGAGATTTTCAAGTCGTCTAAAGAACAGTTGCTAATACAGTT
ACTTATTTCAATAAATAATTGGTTCTAATAATACAAAACATATTCGAGGATATGCAGAAAAAAAGATGTTTGTTATTTTGAAAAGCTTGA
GTAGTTTCTCTCCGAGGTGTAGCGAAGAAGCATCATCTACTTTGTAATGTAATTTTCTTTATGTTTTCACTTTGTAATTTTATTTGTGTT
AATGTACCATGGCCGATATCGGTTTTATTGAAAGAAAATTTATGTTACTTCTGTTTTGGCTTTGCAATCAGTTATGCTAGTTTTCTTATA
CCCTTTCGTAAGCTTCCTAAGGAATCGTTCATTGATTTCCACTGCTTCATTGTATATTAAAACTTTACAACTGTATCGACCATCATATAA
TTCTGGGTCAAGAGATGAAAATAGAACACCACATCGTAAAGTGAAAT

*AtTAS1c*
AtmiR173a TS
syn-tasiR-AtFT
syn-tasiR-AtTry

***>AtTAS1c-AtTry-AtFt***
AAACCTAAACCTAAACGGCTAAGCCCGACGTCAAATACCAAAAAGAGAAAAACAAGAGCGCCGTCAAGCTCTGCAAATACGATCTGTAAG
TCCATCTTAACACAAAAGTGAGATGGGTTCTTAGATCATGTTCCGCCGTTAGATCGAGTCATGGTCTTGTCTCATAGAAAGGTACTTTCG
TTTACTTCTTTTGAGTATCGAGTAGAGCGTCGTCTATAGTTAGTTTGAGATTGCGTTTGTCAGAAGTTAGGTTCAATGTCCCGGTCCAAT
TTTCACCAGCCATGTGTCAGTTTCGTTCCTTCCCGTCCTCTTCTTTGATTTCGTTGGGTTACGGATGTTTTCGAGATGAAACAGCATTGT
TTTGTTGTGATTTTTCTCTACAAGCGAATAGACCATTTATCCCATTCGATACTGCTCGCCTTGGTTATAAAGGAAGAGGCCTCGGTGGAT
CTTAGAAAATTATTCTAAGTCCAACATAGCGTATTCTAAGTTCAACATATCGACGAACTAGAAAAGACATTGGACATATTCCAGGATATG
CAAAAGAAAACAATGAATATTGTTTTGAATGTGTTCAAGTAAATGAGATTTTCAAGTCGTCTAAAGAACAGTTGCTAATACAGTTACTTA
TTTCAATAAATAATTGGTTCTAATAATACAAAACATATTCGAGGATATGCAGAAAAAAAGATGTTTGTTATTTTGAAAAGCTTGAGTAGT
TTCTCTCCGAGGTGTAGCGAAGAAGCATCATCTACTTTGTAATGTAATTTTCTTTATGTTTTCACTTTGTAATTTTATTTGTGTTAATGT
ACCATGGCCGATATCGGTTTTATTGAAAGAAAATTTATGTTACTTCTGTTTTGGCTTTGCAATCAGTTATGCTAGTTTTCTTATACCCTT
TCGTAAGCTTCCTAAGGAATCGTTCATTGATTTCCACTGCTTCATTGTATATTAAAACTTTACAACTGTATCGACCATCATATAATTCTG
GGTCAAGAGATGAAAATAGAACACCACATCGTAAAGTGAAAT

*AtTAS1c*
AtmiR173a TS
syn-tasiR-AtTry
syn-tasiR-AtFT

***>AtTAS1c-NbSu***
AAACCTAAACCTAAACGGCTAAGCCCGACGTCAAATACCAAAAAGAGAAAAACAAGAGCGCCGTCAAGCTCTGCAAATACGATCTGTAAG
TCCATCTTAACACAAAAGTGAGATGGGTTCTTAGATCATGTTCCGCCGTTAGATCGAGTCATGGTCTTGTCTCATAGAAAGGTACTTTCG
TTTACTTCTTTTGAGTATCGAGTAGAGCGTCGTCTATAGTTAGTTTGAGATTGCGTTTGTCAGAAGTTAGGTTCAATGTCCCGGTCCAAT
TTTCACCAGCCATGTGTCAGTTTCGTTCCTTCCCGTCCTCTTCTTTGATTTCGTTGGGTTACGGATGTTTTCGAGATGAAACAGCATTGT
TTTGTTGTGATTTTTCTCTACAAGCGAATAGACCATTTATGTATGACTCCCGGAATTCCATCGGTGGATCTTAGAAAATTATTCTAAGTC
CAACATAGCGTATTCTAAGTTCAACATATCGACGAACTAGAAAAGACATTGGACATATTCCAGGATATGCAAAAGAAAACAATGAATATT
GTTTTGAATGTGTTCAAGTAAATGAGATTTTCAAGTCGTCTAAAGAACAGTTGCTAATACAGTTACTTATTTCAATAAATAATTGGTTCT
AATAATACAAAACATATTCGAGGATATGCAGAAAAAAAGATGTTTGTTATTTTGAAAAGCTTGAGTAGTTTCTCTCCGAGGTGTAGCGAA
GAAGCATCATCTACTTTGTAATGTAATTTTCTTTATGTTTTCACTTTGTAATTTTATTTGTGTTAATGTACCATGGCCGATATCGGTTTT
ATTGAAAGAAAATTTATGTTACTTCTGTTTTGGCTTTGCAATCAGTTATGCTAGTTTTCTTATACCCTTTCGTAAGCTTCCTAAGGAATC
GTTCATTGATTTCCACTGCTTCATTGTATATTAAAACTTTACAACTGTATCGACCATCATATAATTCTGGGTCAAGAGATGAAAATAGAA
CACCACATCGTAAAGTGAAAT

*AtTAS1c*
AtmiR173a TS
syn-tasiR-NbSu

### 2. Minimal syn-tasiRNA precursors

***>AtmiR173TS-GUS$_{At}$***
GTGATTTTTCTCTACAAGCGAATAGACCATTTATGCGCTTGCTGAGTTTCCCCC

*AtTAS1c*
AtmiR173a TS
syn-tasiR-GUS$_{At}$

***>AtmiR173TS-AtFT***
GTGATTTTTCTCTACAAGCGAATAGACCATTTATTGGTTATAAAGGAAGAGGCC

*AtTAS1c*
AtmiR173a TS
syn-tasiR-AtFT

***>AtmiR173TS-AtCH42***
GTGATTTTTCTCTACAAGCGAATAGACCATTTATTAAGTGTCACGGAAATCCCT

*AtTAS1c*
AtmiR173a TS
syn-tasiR-AtCH42

***>AtmiR173TS-AtFT-AtTry***
GTGATTTTTCTCTACAAGCGAATAGACCATTTATTGGTTATAAAGGAAGAGGCCTCCCATTCGATACTGCTCGCC

*AtTAS1c*
AtmiR173a TS
syn-tasiR-AtFT
syn-tasiR-AtTry

***>AtmiR173TS-AtTry-AtFT***
GTGATTTTTCTCTACAAGCGAATAGACCATTTATCCCATTCGATACTGCTCGCCTTGGTTATAAAGGAAGAGGCC

*AtTAS1c*
AtmiR173a TS
syn-tasiR-AtTry
syn-tasiR-AtFT

***>AtmiR173TS-NbSu***
GTGATTTTTCTCTACAAGCGAATAGACCATTTATGTATGACTCCCGGAATTCCA

AtTAS1c
AtmiR173a TS
syn-tasiR-NbSu

***>AtmiR173TS-TSWV(x4)***
GTGATTTTTCTCTACAAGCGAATAGACCATTTATGTAAGACGTGATTGTGTCCTTATCAGCTCTGGGTGAATCGGTTGGTATAG
TGGGGCATACCGTCAGAGTGCACAATCCATCTT

*AtTAS1c*
AtmiR173a TS
syn-tasiR-TSWV-1
syn-tasiR-TSWV-2
syn-tasiR-TSWV-3
syn-tasiR-TSWV-4

***>NbmiR482aTS-NbSu***

GTGGTATGGGGGGAGTCGGGAATAGACCATTTATGTATGACTCCCGGAATTCCA

*AtTAS1c*
NbmiR482a TS
syn-tasiR-NbSu

***>NbmiR482aTS-GUS(x4)***
GTGGTATGGGGGGAGTCGGGAATAGACCATTTATATTGACCCACACTTTGCCGATAACCTTCACCCGGTTGCCACTATTGACCC
ACACTTTGCCGATAACCTTCACCCGGTTGCCAC

*AtTAS1c*
NbmiR482a TS
syn-tasiR-GUS$_{Nb}$-1
syn-tasiR-GUS$_{Nb}$-2

***>NbmiR482aTS-TSWV(x4)***
GTGGTATGGGGGGAGTCGGGAATAGACCATTTATGTAAGACGTGATTGTGTCCTTATCAGCTCTGGGTGAATCGGTTGGTATAG
TGGGGCATACCGTCAGAGTGCACAATCCATCTT

*AtTAS1c*
NbmiR482a TS
syn-tasiR-TSWV-1
syn-tasiR-TSWV-2
syn-tasiR-TSWV-3
syn-tasiR-TSWV-4

***>NbmiR6019aTS-NbSu***
AAACATTTACAAGTCACCTGTATAGACCATTTATGTATGACTCCCGGAATTCCA

*AtTAS1c*
NbmiR6019a TS
syn-tasiR-NbSu

***>NbmiR156aTS-NbSu***
GTGCTCACTCTCTTCTGTCATAGACCATTTATGTATGACTCCCGGAATTCCA

*AtTAS1c*
NbmiR156a TS
syn-tasiR-NbSu

***>AtmiR173aTS-NbSu***
GTGATTTTTCTCTACAAGCGAATAGACCATTTATGTATGACTCCCGGAATTCCA

*AtTAS1c*
AtmiR173a TS
syn-tasiR-NbSu

### 3. amiRNA precursors

***>amiR-GUS$_{Nb}$***
GTAGAGAAGAATCTGTATATTGACCCACACTTTGCCGAATGATGATCACATTCGTTATCTATTTTTTAGGCAAAGTTTGGGTCAATACAt
tggctcttcttact

*AtMIR390a*
amiR-GUS
amiR-GUS*

***>amiR-TSWV***

GTAGAGAAGAATCTGTATGTAAGACGTGATTGTGTCCTATGATGATCACATTCGTTATCTATTTTTTAGGACACAATAACGTCTTACACA
ttggctcttcttact

*AtMIR390a*
amiR-TSWV
amiR-TSWV*

**Text S4.** DNA sequence of *Bsa*I-*ccd*B-based (B/c) vectors used for direct cloning of syn-tasiRNAs.

**>*pENTR-B/c*** (4049 bp)

CTTTCCTGCGTTATCCCCTGATTCTGTGGATAACCGTATTACCGCCTTTGAGTGAGCTGATACCGCTCGCCGCAGCCGAACGACCGAGCG
CAGCGAGTCAGTGAGCGAGGAAGCGGAAGAGCGCCCAATACGCAAACCGCCTCTCCCCGCGCGTTGGCCGATTCATTAATGCAGCTGGCA
CGACAGGTTTCCCGACTGGAAAGCGGGCAGTGAGCGCAACGCAATTAATACGCGTACCGCTAGCCAGGAAGAGTTTGTAGAAACGCAAAA
AGGCCATCCGTCAGGATGGCCTTCTGCTTAGTTTGATGCCTGGCAGTTTATGGCGGGCGTCCTGCCCGCCACCCTCCGGGCCGTTGCTTC
ACAACGTTCAAATCCGCTCCCGGCGGATTTGTCCTACTCAGGAGAGCGTTCACCGACAAACAACAGATAAAACGAAAGGCCCAGTCTTCC
GACTGAGCCTTTCGTTTTATTTGATGCCTGGCAGTTCCCTACTCTCGCGTTAACGCTAGCATGGATGTTTTCCCAGTCACGACGTTGTAA
AACGACGGCCAGTCTTAAGCTCGGGCCCCAAATAATGATTTTATTTTGACTGATAGTGACCTGTTCGTTGCAACAAATTGATGAGCAATG
CTTTTTTATAATGCCAACTTTGTACAAAAAAGCAGGCTCCGCGGCCGCCCCCTTCACCTGTAAGAGACCATTAGGCACCCCAGGCTTTAC
ACTTTATGCTTCCGGCTCGTATAATGTGTGGATTTTGAGTTAGGAGCCGTCGAGATTTTCAGGAGCTAAGGAAGCTAAAATGGAGAAAAA
AATCACTGGATATACCACCGTTGATATATCCCAATGGCATCGTAAAGAACATTTTGAGGCATTTCAGTCAGTTGCTCAATGTACCTATAA
CCAGACCGTTCAGCTGGATATTACGGCCTTTTTAAAGACCGTAAAGAAAAATAAGCACAAGTTTTATCCGGCCTTTATTCACATTCTTGC
CCGCCTGATGAATGCTCATCCGGAGTTCCGTATGGCAATGAAAGACGGTGAGCTGGTGATATGGGATAGTGTTCACCCTTGTTACACCGT
TTTCCATGAGCAAACTGAAACGTTTTCATCGCTCTGGAGTGAATACCACGACGATTTCCGGCAGTTTCTACACATATATTCGCAAGATGT
GGCGTGTTACGGTGAAAACCTGGCCTATTTCCCTAAAGGGTTTATTGAGAATATGTTTTTCGTCTCAGCCAATCCCTGGGTGAGTTTCAC
CAGTTTTGATTTAAACGTGGCCAATATGGACAACTTCTTCGCCCCCGTTTTCACCATGGGCAAATATTATACGCAAGGCGACAAGGTGCT
GATGCCGCTGGCGATTCAGGTTCATCATGCCGTTTGTGATGGCTTCCATGTCGGCAGAATGCTTAATGAATTACAACAGTACTGCGATGA
GTGGCAGGGCGGGGCGTAAACGCGTGGAGCCGGCTTACTAAAAGCCAGATAACAGTATGCGTATTTGCGCGCTGATTTTTGCGGTATAAG
AATATATACTGATATGTATACCCGAAGTATGTCAAAAAGAGGTATGCTATGAAGCAGCGTATTACAGTGACAGTTGACAGCGACAGCTAT
CAGTTGCTCAAGGCATATATGATGTCAATATCTCCGGTCTGGTAAGCACAACCATGCAGAATGAAGCCCGTCGTCTGCGTGCCGAACGCT
GGAAAGCGGAAAATCAGGAAGGGATGGCTGAGGTCGCCCGGTTTATTGAAATGAACGGCTCTTTTGCTGACGAGAACAGGGGCTGGTGAA
ATGCAGTTTAAGGTTTACACCTATAAAAGAGAGAGCCGTTATCGTCTGTTTGTGGATGTACAGAGTGATATTATTGACACGCCCGGCCGA
CGGATGGTGATCCCCCTGGCCAGTGCACGTCTGCTGTCAGATAAAGTCTCCCGTGAACTTTACCCGGTGGTGCATATCGGGGATGAAAGC
TGGCGCATGATGACCACCGATATGGCCAGTGTGCCGGTTTCCGTTATCGGGGAAGAAGTGGCTGATCTCAGCCACCGCGAAAATGACATC
AAAAACGCCATTAACCTGATGTTCTGGGGAATATAAATGTCAGGCTCCCTTATACACAGCCAGTCTGCACCTCGACGGTCTCACATTAAG
GGTGGGCGCGCCGACCCAGCTTTCTTGTACAAAGTTGGCATTATAAGAAAGCATTGCTTATCAATTTGTTGCAACGAACAGGTCACTATC
AGTCAAAATAAAATCATTATTTGCCATCCAGCTGATATCCCCTATAGTGAGTCGTATTACATGGTCATAGCTGTTTCCTGGCAGCTCTGG
CCCGTGTCTCAAAATCTCTGATGTTACATTGCACAAGATAAAAATATATCATCATGAACAATAAAACTGTCTGCTTACATAAACAGTAAT
ACAAGGGGTGTTATGAGCCATATTCAACGGGAAACGTCGAGGCCGCGATTAAATTCCAACATGGATGCTGATTTATATGGGTATAAATGG
GCTCGCGATAATGTCGGGCAATCAGGTGCGACAATCTATCGCTTGTATGGGAAGCCCGATGCGCCAGAGTTGTTTCTGAAACATGGCAAA
GGTAGCGTTGCCAATGATGTTACAGATGAGATGGTCAGACTAAACTGGCTGACGGAATTTATGCCTCTTCCGACCATCAAGCATTTTATC
CGTACTCCTGATGATGCATGGTTACTCACCACTGCGATCCCCGGAAAAACAGCATTCCAGGTATTAGAAGAATATCCTGATTCAGGTGAA
AATATTGTTGATGCGCTGGCAGTGTTCCTGCGCCGGTTGCATTCGATTCCTGTTTGTAATTGTCCTTTTAACAGCGATCGCGTATTTCGT
CTCGCTCAGGCGCAATCACGAATGAATAACGGTTTGGTTGATGCGAGTGATTTTGATGACGAGCGTAATGGCTGGCCTGTTGAACAAGTC
TGGAAAGAAATGCATAAACTTTTGCCATTCTCACCGGATTCAGTCGTCACTCATGGTGATTTCTCACTTGATAACCTTATTTTTGACGAG
GGGAAATTAATAGGTTGTATTGATGTTGGACGAGTCGGAATCGCAGACCGATACCAGGATCTTGCCATCCTATGGAACTGCCTCGGTGAG
TTTTCTCCTTCATTACAGAAACGGCTTTTTCAAAAATATGGTATTGATAATCCTGATATGAATAAATTGCAGTTTCATTTGATGCTCGAT
GAGTTTTTCTAATCAGAATTGGTTAATTGGTTGTAACACTGGCAGAGCATTACGCTGACTTGACGGGACGGCGCAAGCTCATGACCAAAA
TCCCTTAACGTGAGTTACGCGTCGTTCCACTGAGCGTCAGACCCCGTAGAAAAGATCAAAGGATCTTCTTGAGATCCTTTTTTTCTGCGC
GTAATCTGCTGCTTGCAAACAAAAAAACCACCGCTACCAGCGGTGGTTTGTTTGCCGGATCAAGAGCTACCAACTCTTTTTCCGAAGGTA
ACTGGCTTCAGCAGAGCGCAGATACCAAATACTGTCCTTCTAGTGTAGCCGTAGTTAGGCCACCACTTCAAGAACTCTGTAGCACCGCCT
ACATACCTCGCTCTGCTAATCCTGTTACCAGTGGCTGCTGCCAGTGGCGATAAGTCGTGTCTTACCGGGTTGGACTCAAGACGATAGTTA
CCGGATAAGGCGCAGCGGTCGGGCTGAACGGGGGGTTCGTGCACACAGCCCAGCTTGGAGCGAACGACCTACACCGAACTGAGATACCTA
CAGCGTGAGCATTGAGAAAGCGCCACGCTTCCCGAAGGGAGAAAGGCGGACAGGTATCCGGTAAGCGGCAGGGTCGGAACAGGAGAGCGC
ACGAGGGAGCTTCCAGGGGGAAACGCCTGGTATCTTTATAGTCCTGTCGGGTTTCGCCACCTCTGACTTGAGCGTCGATTTTTGTGATGC
TCGTCAGGGGGGCGGAGCCTATGGAAAAACGCCAGCAACGCGGCCTTTTTACGGTTCCTGGCCTTTTGCTGGCCTTTTGCTCACATGTT

M13-F binding site
M13-Reverse binding site
attL1
attL2
Chloramphenicol resistance gene
*ccd*B gene
*Bsa*I site
inverted *Bsa*I site
Kanamycin resistance gene

**> *pENTR-AtmiR173aTS-B/c*** (4082 bp)

CTTTCCTGCGTTATCCCCTGATTCTGTGGATAACCGTATTACCGCCTTTGAGTGAGCTGATACCGCTCGCCGCAGCCGAACGACCGAGCG
CAGCGAGTCAGTGAGCGAGGAAGCGGAAGAGCGCCCAATACGCAAACCGCCTCTCCCCGCGCGTTGGCCGATTCATTAATGCAGCTGGCA
CGACAGGTTTCCCGACTGGAAAGCGGGCAGTGAGCGCAACGCAATTAATACGCGTACCGCTAGCCAGGAAGAGTTTGTAGAAACGCAAAA
AGGCCATCCGTCAGGATGGCCTTCTGCTTAGTTTGATGCCTGGCAGTTTATGGCGGGCGTCCTGCCCGCCACCCTCCGGGCCGTTGCTTC
ACAACGTTCAAATCCGCTCCCGGCGGATTTGTCCTACTCAGGAGAGCGTTCACCGACAAACAACAGATAAAACGAAAGGCCCAGTCTTCC
GACTGAGCCTTTCGTTTTATTTGATGCCTGGCAGTTCCCTACTCTCGCGTTAACGCTAGCATGGATGTTTTCCCAGTCACGACGTTGTAA
AACGACGGCCAGTCTTAAGCTCGGGCCCCAAATAATGATTTTATTTTGACTGATAGTGACCTGTTCGTTGCAACAAATTGATGAGCAATG
CTTTTTTATAATGCCAACTTTGTACAAAAAAGCAGGCTCCGCGGCCGCCCCCTTCACCTGTA**GTGATTTTTCTCTACAAGCGAATAGACC**
**ATTTA**AGAGACCATTAGGCACCCCAGGCTTTACACTTTATGCTTCCGGCTCGTATAATGTGTGGATTTTGAGTTAGGAGCCGTCGAGATT
TTCAGGAGCTAAGGAAGCTAAAATGGAGAAAAAAATCACTGGATATACCACCGTTGATATATCCCAATGGCATCGTAAAGAACATTTTGA
GGCATTTCAGTCAGTTGCTCAATGTACCTATAACCAGACCGTTCAGCTGGATATTACGGCCTTTTTAAAGACCGTAAAGAAAAATAAGCA
CAAGTTTTATCCGGCCTTTATTCACATTCTTGCCCGCCTGATGAATGCTCATCCGGAGTTCCGTATGGCAATGAAAGACGGTGAGCTGGT
GATATGGGATAGTGTTCACCCTTGTTACACCGTTTTCCATGAGCAAACTGAAACGTTTTCATCGCTCTGGAGTGAATACCACGACGATTT
CCGGCAGTTTCTACACATATATTCGCAAGATGTGGCGTGTTACGGTGAAAACCTGGCCTATTTCCCTAAAGGGTTTATTGAGAATATGTT
TTTCGTCTCAGCCAATCCCTGGGTGAGTTTCACCAGTTTTGATTTAAACGTGGCCAATATGGACAACTTCTTCGCCCCCGTTTTCACCAT
GGGCAAATATTATACGCAAGGCGACAAGGTGCTGATGCCGCTGGCGATTCAGGTTCATCATGCCGTTTGTGATGGCTTCCATGTCGGCAG
AATGCTTAATGAATTACAACAGTACTGCGATGAGTGGCAGGGCGGGGCGTAAACGCGTGGAGCCGGCTTACTAAAAGCCAGATAACAGTA
TGCGTATTTGCGCGCTGATTTTTGCGGTATAAGAATATATACTGATATGTATACCCGAAGTATGTCAAAAAGAGGTATGCTATGAAGCAG
CGTATTACAGTGACAGTTGACAGCGACAGCTATCAGTTGCTCAAGGCATATATGATGTCAATATCTCCGGTCTGGTAAGCACAACCATGC
AGAATGAAGCCCGTCGTCTGCGTGCCGAACGCTGGAAAGCGGAAAATCAGGAAGGGATGGCTGAGGTCGCCCGGTTTATTGAAATGAACG
GCTCTTTTGCTGACGAGAACAGGGGCTGGTGAAATGCAGTTTAAGGTTTACACCTATAAAAGAGAGAGCCGTTATCGTCTGTTTGTGGAT
GTACAGAGTGATATTATTGACACGCCCGGCCGACGGATGGTGATCCCCCTGGCCAGTGCACGTCTGCTGTCAGATAAAGTCTCCCGTGAA
CTTTACCCGGTGGTGCATATCGGGGATGAAAGCTGGCGCATGATGACCACCGATATGGCCAGTGTGCCGGTTTCCGTTATCGGGGAAGAA
GTGGCTGATCTCAGCCACCGCGAAAATGACATCAAAAACGCCATTAACCTGATGTTCTGGGGAATATAAATGTCAGGCTCCCTTATACAC
AGCCAGTCTGCACCTCGACGGTCTCACATTAAGGGTGGGCGCGCCGACCCAGCTTTCTTGTACAAAGTTGGCATTATAAGAAAGCATTGC
TTATCAATTTGTTGCAACGAACAGGTCACTATCAGTCAAAATAAAATCATTATTTGCCATCCAGCTGATATCCCCTATAGTGAGTCGTAT
TACATGGTCATAGCTGTTTCCTGGCAGCTCTGGCCCGTGTCTCAAAATCTCTGATGTTACATTGCACAAGATAAAAATATATCATCATGA
ACAATAAAACTGTCTGCTTACATAAACAGTAATACAAGGGGTGTTATGAGCCATATTCAACGGGAAACGTCGAGGCCGCGATTAAATTCC
AACATGGATGCTGATTTATATGGGTATAAATGGGCTCGCGATAATGTCGGGCAATCAGGTGCGACAATCTATCGCTTGTATGGGAAGCCC
GATGCGCCAGAGTTGTTTCTGAAACATGGCAAAGGTAGCGTTGCCAATGATGTTACAGATGAGATGGTCAGACTAAACTGGCTGACGGAA
TTTATGCCTCTTCCGACCATCAAGCATTTTATCCGTACTCCTGATGATGCATGGTTACTCACCACTGCGATCCCCGGAAAAACAGCATTC
CAGGTATTAGAAGAATATCCTGATTCAGGTGAAAATATTGTTGATGCGCTGGCAGTGTTCCTGCGCCGGTTGCATTCGATTCCTGTTTGT
AATTGTCCTTTTAACAGCGATCGCGTATTTCGTCTCGCTCAGGCGCAATCACGAATGAATAACGGTTTGGTTGATGCGAGTGATTTTGAT
GACGAGCGTAATGGCTGGCCTGTTGAACAAGTCTGGAAAGAAATGCATAAACTTTTGCCATTCTCACCGGATTCAGTCGTCACTCATGGT
GATTTCTCACTTGATAACCTTATTTTTGACGAGGGGAAATTAATAGGTTGTATTGATGTTGGACGAGTCGGAATCGCAGACCGATACCAG
GATCTTGCCATCCTATGGAACTGCCTCGGTGAGTTTTCTCCTTCATTACAGAAACGGCTTTTTCAAAAATATGGTATTGATAATCCTGAT
ATGAATAAATTGCAGTTTCATTTGATGCTCGATGAGTTTTTCTAATCAGAATTGGTTAATTGGTTGTAACACTGGCAGAGCATTACGCTG
ACTTGACGGGACGGCGCAAGCTCATGACCAAAATCCCTTAACGTGAGTTACGCGTCGTTCCACTGAGCGTCAGACCCCGTAGAAAAGATC
AAAGGATCTTCTTGAGATCCTTTTTTTCTGCGCGTAATCTGCTGCTTGCAAACAAAAAAACCACCGCTACCAGCGGTGGTTTGTTTGCCG
GATCAAGAGCTACCAACTCTTTTTCCGAAGGTAACTGGCTTCAGCAGAGCGCAGATACCAAATACTGTCCTTCTAGTGTAGCCGTAGTTA
GGCCACCACTTCAAGAACTCTGTAGCACCGCCTACATACCTCGCTCTGCTAATCCTGTTACCAGTGGCTGCTGCCAGTGGCGATAAGTCG
TGTCTTACCGGGTTGGACTCAAGACGATAGTTACCGGATAAGGCGCAGCGGTCGGGCTGAACGGGGGGTTCGTGCACACAGCCCAGCTTG
GAGCGAACGACCTACACCGAACTGAGATACCTACAGCGTGAGCATTGAGAAAGCGCCACGCTTCCCGAAGGGAGAAAGGCGGACAGGTAT
CCGGTAAGCGGCAGGGTCGGAACAGGAGAGCGCACGAGGGAGCTTCCAGGGGGAAACGCCTGGTATCTTTATAGTCCTGTCGGGTTTCGC
CACCTCTGACTTGAGCGTCGATTTTTGTGATGCTCGTCAGGGGGGCGGAGCCTATGGAAAAACGCCAGCAACGCGGCCTTTTTACGGTTC
CTGGCCTTTTGCTGGCCTTTTGCTCACATGTT

**AtmiR173a target site**
**AtTAS1c-derived spacer**
M13-F binding site
M13-Reverse binding site
attL1
attL2
Chloramphenicol resistance gene
*ccd*B gene
*Bsa*I site
inverted *Bsa*I site
Kanamycin resistance gene

***>pENTR-NbmiR482aTS-B/c*** (4082 bp)

CTTTCCTGCGTTATCCCCTGATTCTGTGGATAACCGTATTACCGCCTTTGAGTGAGCTGATACCGCTCGCCGCAGCCGAACGACCGAGCG
CAGCGAGTCAGTGAGCGAGGAAGCGGAAGAGCGCCCAATACGCAAACCGCCTCTCCCCGCGCGTTGGCCGATTCATTAATGCAGCTGGCA
CGACAGGTTTCCCGACTGGAAAGCGGGCAGTGAGCGCAACGCAATTAATACGCGTACCGCTAGCCAGGAAGAGTTTGTAGAAACGCAAAA
AGGCCATCCGTCAGGATGGCCTTCTGCTTAGTTTGATGCCTGGCAGTTTATGGCGGGCGTCCTGCCCGCCACCCTCCGGGCCGTTGCTTC
ACAACGTTCAAATCCGCTCCCGGCGGATTTGTCCTACTCAGGAGAGCGTTCACCGACAAACAACAGATAAAACGAAAGGCCCAGTCTTCC
GACTGAGCCTTTCGTTTTATTTGATGCCTGGCAGTTCCCTACTCTCGCGTTAACGCTAGCATGGATGTTTTCCCAGTCACGACGTTGTAA
AACGACGGCCAGTCTTAAGCTCGGGCCCCAAATAATGATTTTATTTTGACTGATAGTGACCTGTTCGTTGCAACAAATTGATGAGCAATG
CTTTTTTATAATGCCAACTTTGTACAAAAAAGCAGGCTCCGCGGCCGCCCCCTTCACCTGTA**GTGGTATGGGGGGAGTCGGGAATAGACC**
**ATTTA**AGAGACCATTAGGCACCCCAGGCTTTACACTTTATGCTTCCGGCTCGTATAATGTGTGGATTTTGAGTTAGGAGCCGTCGAGATT
TTCAGGAGCTAAGGAAGCTAAAATGGAGAAAAAAATCACTGGATATACCACCGTTGATATATCCCAATGGCATCGTAAAGAACATTTTGA
GGCATTTCAGTCAGTTGCTCAATGTACCTATAACCAGACCGTTCAGCTGGATATTACGGCCTTTTTAAAGACCGTAAAGAAAAATAAGCA
CAAGTTTTATCCGGCCTTTATTCACATTCTTGCCCGCCTGATGAATGCTCATCCGGAGTTCCGTATGGCAATGAAAGACGGTGAGCTGGT
GATATGGGATAGTGTTCACCCTTGTTACACCGTTTTCCATGAGCAAACTGAAACGTTTTCATCGCTCTGGAGTGAATACCACGACGATTT
CCGGCAGTTTCTACACATATATTCGCAAGATGTGGCGTGTTACGGTGAAAACCTGGCCTATTTCCCTAAAGGGTTTATTGAGAATATGTT
TTTCGTCTCAGCCAATCCCTGGGTGAGTTTCACCAGTTTTGATTTAAACGTGGCCAATATGGACAACTTCTTCGCCCCCGTTTTCACCAT
GGGCAAATATTATACGCAAGGCGACAAGGTGCTGATGCCGCTGGCGATTCAGGTTCATCATGCCGTTTGTGATGGCTTCCATGTCGGCAG
AATGCTTAATGAATTACAACAGTACTGCGATGAGTGGCAGGGCGGGGCGTAAACGCGTGGAGCCGGCTTACTAAAAGCCAGATAACAGTA
TGCGTATTTGCGCGCTGATTTTTGCGGTATAAGAATATATACTGATATGTATACCCGAAGTATGTCAAAAAGAGGTATGCTATGAAGCAG
CGTATTACAGTGACAGTTGACAGCGACAGCTATCAGTTGCTCAAGGCATATATGATGTCAATATCTCCGGTCTGGTAAGCACAACCATGC
AGAATGAAGCCCGTCGTCTGCGTGCCGAACGCTGGAAAGCGGAAAATCAGGAAGGGATGGCTGAGGTCGCCCGGTTTATTGAAATGAACG
GCTCTTTTGCTGACGAGAACAGGGGCTGGTGAAATGCAGTTTAAGGTTTACACCTATAAAAGAGAGAGCCGTTATCGTCTGTTTGTGGAT
GTACAGAGTGATATTATTGACACGCCCGGCCGACGGATGGTGATCCCCCTGGCCAGTGCACGTCTGCTGTCAGATAAAGTCTCCCGTGAA
CTTTACCCGGTGGTGCATATCGGGGATGAAAGCTGGCGCATGATGACCACCGATATGGCCAGTGTGCCGGTTTCCGTTATCGGGGAAGAA
GTGGCTGATCTCAGCCACCGCGAAAATGACATCAAAAACGCCATTAACCTGATGTTCTGGGGAATATAAATGTCAGGCTCCCTTATACAC
AGCCAGTCTGCACCTCGACGGTCTCACATTAAGGGTGGGCGCGCCGACCCAGCTTTCTTGTACAAAGTTGGCATTATAAGAAAGCATTGC
TTATCAATTTGTTGCAACGAACAGGTCACTATCAGTCAAAATAAAATCATTATTTGCCATCCAGCTGATATCCCCTATAGTGAGTCGTAT
TACATGGTCATAGCTGTTTCCTGGCAGCTCTGGCCCGTGTCTCAAAATCTCTGATGTTACATTGCACAAGATAAAAATATATCATCATGA
ACAATAAAACTGTCTGCTTACATAAACAGTAATACAAGGGGTGTTATGAGCCATATTCAACGGGAAACGTCGAGGCCGCGATTAAATTCC
AACATGGATGCTGATTTATATGGGTATAAATGGGCTCGCGATAATGTCGGGCAATCAGGTGCGACAATCTATCGCTTGTATGGGAAGCCC
GATGCGCCAGAGTTGTTTCTGAAACATGGCAAAGGTAGCGTTGCCAATGATGTTACAGATGAGATGGTCAGACTAAACTGGCTGACGGAA
TTTATGCCTCTTCCGACCATCAAGCATTTTATCCGTACTCCTGATGATGCATGGTTACTCACCACTGCGATCCCCGGAAAAACAGCATTC
CAGGTATTAGAAGAATATCCTGATTCAGGTGAAAATATTGTTGATGCGCTGGCAGTGTTCCTGCGCCGGTTGCATTCGATTCCTGTTTGT
AATTGTCCTTTTAACAGCGATCGCGTATTTCGTCTCGCTCAGGCGCAATCACGAATGAATAACGGTTTGGTTGATGCGAGTGATTTTGAT
GACGAGCGTAATGGCTGGCCTGTTGAACAAGTCTGGAAAGAAATGCATAAACTTTTGCCATTCTCACCGGATTCAGTCGTCACTCATGGT
GATTTCTCACTTGATAACCTTATTTTTGACGAGGGGAAATTAATAGGTTGTATTGATGTTGGACGAGTCGGAATCGCAGACCGATACCAG
GATCTTGCCATCCTATGGAACTGCCTCGGTGAGTTTTCTCCTTCATTACAGAAACGGCTTTTTCAAAAATATGGTATTGATAATCCTGAT
ATGAATAAATTGCAGTTTCATTTGATGCTCGATGAGTTTTTCTAATCAGAATTGGTTAATTGGTTGTAACACTGGCAGAGCATTACGCTG
ACTTGACGGGACGGCGCAAGCTCATGACCAAAATCCCTTAACGTGAGTTACGCGTCGTTCCACTGAGCGTCAGACCCCGTAGAAAAGATC
AAAGGATCTTCTTGAGATCCTTTTTTTCTGCGCGTAATCTGCTGCTTGCAAACAAAAAAACCACCGCTACCAGCGGTGGTTTGTTTGCCG
GATCAAGAGCTACCAACTCTTTTTCCGAAGGTAACTGGCTTCAGCAGAGCGCAGATACCAAATACTGTCCTTCTAGTGTAGCCGTAGTTA
GGCCACCACTTCAAGAACTCTGTAGCACCGCCTACATACCTCGCTCTGCTAATCCTGTTACCAGTGGCTGCTGCCAGTGGCGATAAGTCG
TGTCTTACCGGGTTGGACTCAAGACGATAGTTACCGGATAAGGCGCAGCGGTCGGGCTGAACGGGGGGTTCGTGCACACAGCCCAGCTTG
GAGCGAACGACCTACACCGAACTGAGATACCTACAGCGTGAGCATTGAGAAAGCGCCACGCTTCCCGAAGGGAGAAAGGCGGACAGGTAT
CCGGTAAGCGGCAGGGTCGGAACAGGAGAGCGCACGAGGGAGCTTCCAGGGGGAAACGCCTGGTATCTTTATAGTCCTGTCGGGTTTCGC
CACCTCTGACTTGAGCGTCGATTTTTGTGATGCTCGTCAGGGGGGCGGAGCCTATGGAAAAACGCCAGCAACGCGGCCTTTTTACGGTTC
CTGGCCTTTTGCTGGCCTTTTGCTCACATGTT

**NbmiR482a target site**
**AtTAS1c-derived spacer**
M13-F binding site
M13-Reverse binding site
attL1
attL2
Chloramphenicol resistance gene
*ccd*B gene
*Bsa*I site
inverted *Bsa*I site
Kanamycin resistance gene

***>pMDC32B-B/c*** (11602 bp)

CCAGCCAGCCAACAGCTCCCCGACCGGCAGCTCGGCACAAAATCACCACTCGATACAGGCAGCCCATCAGTCCGGGACGGCGTCAGCGGG
AGAGCCGTTGTAAGGCGGCAGACTTTGCTCATGTTACCGATGCTATTCGGAAGAACGGCAACTAAGCTGCCGGGTTTGAAACACGGATGA
TCTCGCGGAGGGTAGCATGTTGATTGTAACGATGACAGAGCGTTGCTGCCTGTGATCACCGCGGTTTCAAAATCGGCTCCGTCGATACTA
TGTTATACGCCAACTTTGAAAACAACTTTGAAAAAGCTGTTTTCTGGTATTTAAGGTTTTAGAATGCAAGGAACAGTGAATTGGAGTTCG
TCTTGTTATAATTAGCTTCTTGGGGTATCTTTAAATACTGTAGAAAAGAGGAAGGAAATAATAAATGGCTAAAATGAGAATATCACCGGA
ATTGAAAAAACTGATCGAAAAATACCGCTGCGTAAAAGATACGGAAGGAATGTCTCCTGCTAAGGTATATAAGCTGGTGGGAGAAAATGA
AAACCTATATTTAAAAATGACGGACAGCCGGTATAAAGGGACCACCTATGATGTGGAACGGGAAAAGGACATGATGCTATGGCTGGAAGG
AAAGCTGCCTGTTCCAAAGGTCCTGCACTTTGAACGGCATGATGGCTGGAGCAATCTGCTCATGAGTGAGGCCGATGGCGTCCTTTGCTC
GGAAGAGTATGAAGATGAACAAAGCCCTGAAAAGATTATCGAGCTGTATGCGGAGTGCATCAGGCTCTTTCACTCCATCGACATATCGGA
TTGTCCCTATACGAATAGCTTAGACAGCCGCTTAGCCGAATTGGATTACTTACTGAATAACGATCTGGCCGATGTGGATTGCGAAAACTG
GGAAGAAGACACTCCATTTAAAGATCCGCGCGAGCTGTATGATTTTTTAAAGACGGAAAAGCCCGAAGAGGAACTTGTCTTTTCCCACGG
CGACCTGGGAGACAGCAACATCTTTGTGAAAGATGGCAAAGTAAGTGGCTTTATTGATCTTGGGAGAAGCGGCAGGGCGGACAAGTGGTA
TGACATTGCCTTCTGCGTCCGGTCGATCAGGGAGGATATCGGGGAAGAACAGTATGTCGAGCTATTTTTTGACTTACTGGGGATCAAGCC
TGATTGGGAGAAAATAAAATATTATATTTTACTGGATGAATTGTTTTAGTACCTAGAATGCATGACCAAAATCCCTTAACGTGAGTTTTC
GTTCCACTGAGCGTCAGACCCCGTAGAAAAGATCAAAGGATCTTCTTGAGATCCTTTTTTTCTGCGCGTAATCTGCTGCTTGCAAACAAA
AAAACCACCGCTACCAGCGGTGGTTTGTTTGCCGGATCAAGAGCTACCAACTCTTTTTCCGAAGGTAACTGGCTTCAGCAGAGCGCAGAT
ACCAAATACTGTCCTTCTAGTGTAGCCGTAGTTAGGCCACCACTTCAAGAACTCTGTAGCACCGCCTACATACCTCGCTCTGCTAATCCT
GTTACCAGTGGCTGCTGCCAGTGGCGATAAGTCGTGTCTTACCGGGTTGGACTCAAGACGATAGTTACCGGATAAGGCGCAGCGGTCGGG
CTGAACGGGGGGTTCGTGCACACAGCCCAGCTTGGAGCGAACGACCTACACCGAACTGAGATACCTACAGCGTGAGCTATGAGAAAGCGC
CACGCTTCCCGAAGGGAGAAAGGCGGACAGGTATCCGGTAAGCGGCAGGGTCGGAACAGGAGAGCGCACGAGGGAGCTTCCAGGGGGAAA
CGCCTGGTATCTTTATAGTCCTGTCGGGTTTCGCCACCTCTGACTTGAGCGTCGATTTTTGTGATGCTCGTCAGGGGGGCGGAGCCTATG
GAAAAACGCCAGCAACGCGGCCTTTTTACGGTTCCTGGCCTTTTGCTGGCCTTTTGCTCACATGTTCTTTCCTGCGTTATCCCCTGATTC
TGTGGATAACCGTATTACCGCCTTTGAGTGAGCTGATACCGCTCGCCGCAGCCGAACGACCGAGCGCAGCGAGTCAGTGAGCGAGGAAGC
GGAAGAGCGCCTGATGCGGTATTTTCTCCTTACGCATCTGTGCGGTATTTCACACCGCATATGGTGCACTCTCAGTACAATCTGCTCTGA
TGCCGCATAGTTAAGCCAGTATACACTCCGCTATCGCTACGTGACTGGGTCATGGCTGCGCCCCGACACCCGCCAACACCCGCTGACGCG
CCCTGACGGGCTTGTCTGCTCCCGGCATCCGCTTACAGACAAGCTGTGACCGTCTCCGGGAGCTGCATGTGTCAGAGGTTTTCACCGTCA
TCACCGAAACGCGCGAGGCAGGGTGCCTTGATGTGGGCGCCGGCGGTCGAGTGGCGACGGCGCGGCTTGTCCGCGCCCTGGTAGATTGCC
TGGCCGTAGGCCAGCCATTTTTGAGCGGCCAGCGGCCGCGATAGGCCGACGCGAAGCGGCGGGGCGTAGGGAGCGCAGCGACCGAAGGGT
AGGCGCTTTTTGCAGCTCTTCGGCTGTGCGCTGGCCAGACAGTTATGCACAGGCCAGGCGGGTTTTAAGAGTTTTAATAAGTTTTAAAGA
GTTTTAGGCGGAAAAATCGCCTTTTTTCTCTTTTATATCAGTCACTTACATGTGTGACCGGTTCCCAATGTACGGCTTTGGGTTCCCAAT
GTACGGGTTCCGGTTCCCAATGTACGGCTTTGGGTTCCCAATGTACGTGCTATCCACAGGAAAGAGAACTTTTCGACCTTTTTCCCCTGC
TAGGGCAATTTGCCCTAGCATCTGCTCCGTACATTAGGAACCGGCGGATGCTTCGCCCTCGATCAGGTTGCGGTAGCGCATGACTAGGAT
CGGGCCAGCCTGCCCCGCCTCCTCCTTCAAATCGTACTCCGGCAGGTCATTTGACCCGATCAGCTTGCGCACGGTGAAACAGAACTTCTT
GAACTCTCCGGCGCTGCCACTGCGTTCGTAGATCGTCTTGAACAACCATCTGGCTTCTGCCTTGCCTGCGGCGCGGCGTGCCAGGCGGTA
GAGAAAACGGCCGATGCCGGGATCGATCAAAAAGTAATCGGGGTGAACCGTCAGCACGTCCGGGTTCTTGCCTTCTGTGATCTCGCGGTA
CATCCAATCAGCTAGCTCGATCTCGATGTACTCCGGCCGCCCGGTTTCGCTCTTTACGATCTTGTAGCGGCTAATCAAGGCTTCACCCTC
GGATACCGTCACCAGGCGGCCGTTCTTGGCCTTCTTCGTACGCTGCATGGCAACGTGCGTGGTGTTTAACCGAATGCAGGTTTCTACCAG
GTCGTCTTTCTGCTTTCCGCCATCGGCTCGCCGGCAGAACTTGAGTACGTCCGCAACGTGTGGACGGAACACGCGGCCGGGCTTGTCTCC
CTTCCCTTCCCGGTATCGGTTCATGGATTCGGTTAGATGGGAAACCGCCATCAGTACCAGGTCGTAATCCCACACACTGGCCATGCCGGC
CGGCCCTGCGGAAACCTCTACGTGCCCGTCTGGAAGCTCGTAGCGGATCACCTCGCCAGCTCGTCGGTCACGCTTCGACAGACGGAAAAC
GGCCACGTCCATGATGCTGCGACTATCGCGGGTGCCCACGTCATAGAGCATCGGAACGAAAAAATCTGGTTGCTCGTCGCCCTTGGGCGG
CTTCCTAATCGACGGCGCACCGGCTGCCGGCGGTTGCCGGGATTCTTTGCGGATTCGATCAGCGGCCGCTTGCCACGATTCACCGGGGCG
TGCTTCTGCCTCGATGCGTTGCCGCTGGGCGGCCTGCGCGGCCTTCAACTTCTCCACCAGGTCATCACCCAGCGCCGCGCCGATTTGTAC
CGGGCCGGATGGTTTGCGACCGTCACGCCGATTCCTCGGGCTTGGGGGTTCCAGTGCCATTGCAGGGCCGGCAGACAACCCAGCCGCTTA
CGCCTGGCCAACCGCCCGTTCCTCCACACATGGGGCATTCCACGGCGTCGGTGCCTGGTTGTTCTTGATTTTCCATGCCGCCTCCTTTAG
CCGCTAAAATTCATCTACTCATTTATTCATTTGCTCATTTACTCTGGTAGCTGCGCGATGTATTCAGATAGCAGCTCGGTAATGGTCTTG
CCTTGGCGTACCGCGTACATCTTCAGCTTGGTGTGATCCTCCGCCGGCAACTGAAAGTTGACCCGCTTCATGGCTGGCGTGTCTGCCAGG
CTGGCCAACGTTGCAGCCTTGCTGCTGCGTGCGCTCGGACGGCCGGCACTTAGCGTGTTTGTGCTTTTGCTCATTTTCTCTTTACCTCAT
TAACTCAAATGAGTTTTGATTTAATTTCAGCGGCCAGCGCCTGGACCTCGCGGGCAGCGTCGCCCTCGGGTTCTGATTCAAGAACGGTTG
TGCCGGCGGCGGCAGTGCCTGGGTAGCTCACGCGCTGCGTGATACGGGACTCAAGAATGGGCAGCTCGTACCCGGCCAGCGCCTCGGCAA
CCTCACCGCCGATGCGCGTGCCTTTGATCGCCCGCGACACGACAAAGGCCGCTTGTAGCCTTCCATCCGTGACCTCAATGCGCTGCTTAA
CCAGCTCCACCAGGTCGGCGGTGGCCCATATGTCGTAAGGGCTTGGCTGCACCGGAATCAGCACGAAGTCGGCTGCCTTGATCGCGGACA
CAGCCAAGTCCGCCGCCTGGGGCGCTCCGTCGATCACTACGAAGTCGCGCCGGCCGATGGCCTTCACGTCGCGGTCAATCGTCGGGCGGT
CGATGCCGACAACGGTTAGCGGTTGATCTTCCCGCACGGCCGCCCAATCGCGGGCACTGCCCTGGGGATCGGAATCGACTAACAGAACAT
CGGCCCCGGCGAGTTGCAGGGCGCGGGCTAGATGGGTTGCGATGGTCGTCTTGCCTGACCCGCCTTTCTGGTTAAGTACAGCGATAACCT
TCATGCGTTCCCCTTGCGTATTTGTTTATTTACTCATCGCATCATATACGCAGCGACCGCATGACGCAAGCTGTTTTACTCAAATACACA
TCACCTTTTTAGACGGCGGCGCTCGGTTTCTTCAGCGGCCAAGCTGGCCGGCCAGGCCGCCAGCTTGGCATCAGACAAACCGGCCAGGAT
TTCATGCAGCCGCACGGTTGAGACGTGCGCGGGCGGCTCGAACACGTACCCGGCCGCGATCATCTCCGCCTCGATCTCTTCGGTAATGAA
AAACGGTTCGTCCTGGCCGTCCTGGTGCGGTTTCATGCTTGTTCCTCTTGGCGTTCATTCTCGGCGGCCGCCAGGGCGTCGGCCTCGGTC
AATGCGTCCTCACGGAAGGCACCGCGCCGCCTGGCCTCGGTGGGCGTCACTTCCTCGCTGCGCTCAAGTGCGCGGTACAGGGTCGAGCGA
TGCACGCCAAGCAGTGCAGCCGCCTCTTTCACGGTGCGGCCTTCCTGGTCGATCAGCTCGCGGGCGTGCGCGATCTGTGCCGGGGTGAGG
GTAGGGCGGGGGCCAAACTTCACGCCTCGGGCCTTGGCGGCCTCGCGCCCGCTCCGGGTGCGGTCGATGATTAGGGAACGCTCGAACTCG

GCAATGCCGGCGAACACGGTCAACACCATGCGGCCGGCCGGCGTGGTGGTGTCGGCCCACGGCTCTGCCAGGCTACGCAGGCCCGCGCCG
GCCTCCTGGATGCGCTCGGCAATGTCCAGTAGGTCGCGGGTGCTGCGGGCCAGGCGGTCTAGCCTGGTCACTGTCACAACGTCGCCAGGG
CGTAGGTGGTCAAGCATCCTGGCCAGCTCCGGGCGGTCGCGCCTGGTGCCGGTGATCTTCTCGGAAAACAGCTTGGTGCAGCCGGCCGCG
TGCAGTTCGGCCCGTTGGTTGGTCAAGTCCTGGTCGTCGGTGCTGACGCGGGCATAGCCCAGCAGGCCAGCGGCGGCGCTCTTGTTCATG
GCGTAATGTCTCCGGTTCTAGTCGCAAGTATTCTACTTTATGCGACTAAAACACGCGACAAGAAAACGCCAGGAAAAGGGCAGGGCGGCA
GCCTGTCGCGTAACTTAGGACTTGTGCGACATGTCGTTTTCAGAAGACGGCTGCACTGAACGTCAGAAGCCGACTGCACTATAGCAGCGG
AGGGGTTGGATCAAAGTACTTTGATCCCGAGGGGAACCCTGTGGTTGGCATGCACATACAAATGGACGAACGGATAAACCTTTTCACGCC
CTTTTAAATATCCGTTATTCTAATAAACGCTCTTTTCTCTTAGGTTTACCCGCCAATATATCCTGTCAAACACTGATAGTTTAAACTGAA
GGCGGGAAACGACAATCTGATCCAAGCTCAAGCTGCTCTAGCATTCGCCATTCAGGCTGCGCAACTGTTGGGAAGGGCGATCGGTGCGGG
CCTCTTCGCTATTACGCCAGCTGGCGAAAGGGGGATGTGCTGCAAGGCGATTAAGTTGGGTAACGCCAGGGTTTTCCCAGTCACGACGTT
GTAAAACGACGGCCAGTGCCAAGCTTGGCGTGCCTGCAGGTCAACATGGTGGAGCACGACACACTTGTCTACTCCAAAAATATCAAAGAT
ACAGTCTCAGAAGACCAAAGGGCAATTGAGACTTTTCAACAAAGGGTAATATCCGGAAACCTCCTCGGATTCCATTGCCCAGCTATCTGT
CACTTTATTGTGAAGATAGTGGAAAAGGAAGGTGGCTCCTACAAATGCCATCATTGCGATAAAGGAAAGGCCATCGTTGAAGATGCCTCT
GCCGACAGTGGTCCCAAAGATGGACCCCCACCCACGAGGAGCATCGTGGAAAAAGAAGACGTTCCAACCACGTCTTCAAAGCAAGTGGAT
TGATGTGATAACATGGTGGAGCACGACACACTTGTCTACTCCAAAAATATCAAAGATACAGTCTCAGAAGACCAAAGGGCAATTGAGACT
TTTCAACAAAGGGTAATATCCGGAAACCTCCTCGGATTCCATTGCCCAGCTATCTGTCACTTTATTGTGAAGATAGTGGAAAAGGAAGGT
GGCTCCTACAAATGCCATCATTGCGATAAAGGAAAGGCCATCGTTGAAGATGCCTCTGCCGACAGTGGTCCCAAAGATGGACCCCCACCC
ACGAGGAGCATCGTGGAAAAAGAAGACGTTCCAACCACGTCTTCAAAGCAAGTGGATTGATGTGATATCTCCACTGACGTAAGGGATGAC
GCACAATCCCACTATCCTTCGCAAGACCCTTCCTCTATATAAGGAAGTTCATTTCATTTGGAGAGGACCTCGACTCTAGAGGATCCCCGG
GTACCGGGCCCCCCCTCGAGGCGCGCCAAGCTATCAAACAAGTTTGTACAAAAAAGCAGGCTCCGCGGCCGCCCCCTTCACCTGTAAGAG
ACCATTAGGCACCCCAGGCTTTACACTTTATGCTTCCGGCTCGTATAATGTGTGGATTTTGAGTTAGGAGCCGTCGAGATTTTCAGGAGC
TAAGGAAGCTAAAATGGAGAAAAAAATCACTGGATATACCACCGTTGATATATCCCAATGGCATCGTAAAGAACATTTTGAGGCATTTCA
GTCAGTTGCTCAATGTACCTATAACCAGACCGTTCAGCTGGATATTACGGCCTTTTTAAAGACCGTAAAGAAAAATAAGCACAAGTTTTA
TCCGGCCTTTATTCACATTCTTGCCCGCCTGATGAATGCTCATCCGGAGTTCCGTATGGCAATGAAAGACGGTGAGCTGGTGATATGGGA
TAGTGTTCACCCTTGTTACACCGTTTTCCATGAGCAAACTGAAACGTTTTCATCGCTCTGGAGTGAATACCACGACGATTTCCGGCAGTT
TCTACACATATATTCGCAAGATGTGGCGTGTTACGGTGAAAACCTGGCCTATTTCCCTAAAGGGTTTATTGAGAATATGTTTTTCGTCTC
AGCCAATCCCTGGGTGAGTTTCACCAGTTTTGATTTAAACGTGGCCAATATGGACAACTTCTTCGCCCCCGTTTTCACCATGGGCAAATA
TTATACGCAAGGCGACAAGGTGCTGATGCCGCTGGCGATTCAGGTTCATCATGCCGTTTGTGATGGCTTCCATGTCGGCAGAATGCTTAA
TGAATTACAACAGTACTGCGATGAGTGGCAGGGCGGGGCGTAAACGCGTGGAGCCGGCTTACTAAAAGCCAGATAACAGTATGCGTATTT
GCGCGCTGATTTTTGCGGTATAAGAATATATACTGATATGTATACCCGAAGTATGTCAAAAAGAGGTATGCTATGAAGCAGCGTATTACA
GTGACAGTTGACAGCGACAGCTATCAGTTGCTCAAGGCATATATGATGTCAATATCTCCGGTCTGGTAAGCACAACCATGCAGAATGAAG
CCCGTCGTCTGCGTGCCGAACGCTGGAAAGCGGAAAATCAGGAAGGGATGGCTGAGGTCGCCCGGTTTATTGAAATGAACGGCTCTTTTG
CTGACGAGAACAGGGGCTGGTGAAATGCAGTTTAAGGTTTACACCTATAAAAGAGAGAGCCGTTATCGTCTGTTTGTGGATGTACAGAGT
GATATTATTGACACGCCCGGCCGACGGATGGTGATCCCCCTGGCCAGTGCACGTCTGCTGTCAGATAAAGTCTCCCGTGAACTTTACCCG
GTGGTGCATATCGGGGATGAAAGCTGGCGCATGATGACCACCGATATGGCCAGTGTGCCGGTTTCCGTTATCGGGGAAGAAGTGGCTGAT
CTCAGCCACCGCGAAAATGACATCAAAAACGCCATTAACCTGATGTTCTGGGGAATATAAATGTCAGGCTCCCTTATACACAGCCAGTCT
GCACCTCGACGGTCTCACATTAAGGGTGGGCGCGCCGACCCAGCTTTCTTGTACAAAGTGGTTCGATAATTCCTTAATTAACTAGTTCTA
GAGCGGCCGCCCACCGCGGTGGAGCTCGAATTTCCCCGATCGTTCAAACATTTGGCAATAAAGTTTCTTAAGATTGAATCCTGTTGCCGG
TCTTGCGATGATTATCATATAATTTCTGTTGAATTACGTTAAGCATGTAATAATTAACATGTAATGCATGACGTTATTTATGAGATGGGT
TTTTATGATTAGAGTCCCGCAATTATACATTTAATACGCGATAGAAAACAAAATATAGCGCGCAAACTAGGATAAATTATCGCGCGCGGT
GTCATCTATGTTACTGAATTCGTAATCATGGTCATAGCTGTTTCCTGTGTGAAATTGTTATCCGCTCACAATTCCACACAACATACGAGC
CGGAAGCATAAAGTGTAAAGCCTGGGGTGCCTAATGAGTGAGCTAACTCACATTAATTGCGTTGCGCTCACTGCCCGCTTTCCAGTCGGG
AAACCTGTCGTGCCAGCTGCATTAATGAATCGGCCAACGCGCGGGGAGAGGCGGTTTGCGTATTGGCTAGAGCAGCTTGCCAACATGGTG
GAGCACGACACTCTCGTCTACTCCAAGAATATCAAAGATACAGTCTCAGAAGACCAAAGGGCTATTGAGACTTTTCAACAAAGGGTAATA
TCGGGAAACCTCCTCGGATTCCATTGCCCAGCTATCTGTCACTTCATCAAAAGGACAGTAGAAAAGGAAGGTGGCACCTACAAATGCCAT
CATTGCGATAAAGGAAAGGCTATCGTTCAAGATGCCTCTGCCGACAGTGGTCCCAAAGATGGACCCCCACCCACGAGGAGCATCGTGGAA
AAAGAAGACGTTCCAACCACGTCTTCAAAGCAAGTGGATTGATGTGATAACATGGTGGAGCACGACACTCTCGTCTACTCCAAGAATATC
AAAGATACAGTCTCAGAAGACCAAAGGGCTATTGAGACTTTTCAACAAAGGGTAATATCGGGAAACCTCCTCGGATTCCATTGCCCAGCT
ATCTGTCACTTCATCAAAAGGACAGTAGAAAAGGAAGGTGGCACCTACAAATGCCATCATTGCGATAAAGGAAAGGCTATCGTTCAAGAT
GCCTCTGCCGACAGTGGTCCCAAAGATGGACCCCCACCCACGAGGAGCATCGTGGAAAAAGAAGACGTTCCAACCACGTCTTCAAAGCAA
GTGGATTGATGTGATATCTCCACTGACGTAAGGGATGACGCACAATCCCACTATCCTTCGCAAGACCTTCCTCTATATAAGGAAGTTCAT
TTCATTTGGAGAGGACACGCTGAAATCACCAGTCTCTCTCTACAAATCTATCTCTCTCGAGCTTTCGCAGATCCCGGGGGGCAATGAGAT
ATGAAAAAGCCTGAACTCACCGCGACGTCTGTCGAGAAGTTTCTGATCGAAAAGTTCGACAGCGTCTCCGACCTGATGCAGCTCTCGGAG
GGCGAAGAATCTCGTGCTTTCAGCTTCGATGTAGGAGGGCGTGGATATGTCCTGCGGGTAAATAGCTGCGCCGATGGTTTCTACAAAGAT
CGTTATGTTTATCGGCACTTTGCATCGGCCGCGCTCCCGATTCCGGAAGTGCTTGACATTGGGGAGTTTAGCGAGAGCCTGACCTATTGC
ATCTCCCGCCGTGCACAGGGTGTCACGTTGCAAGACCTGCCTGAAACCGAACTGCCCGCTGTTCTACAACCGGTCGCGGAGGCTATGGAT
GCGATCGCTGCGGCCGATCTTAGCCAGACGAGCGGGTTCGGCCCATTCGGACCGCAAGGAATCGGTCAATACACTACATGGCGTGATTTC
ATATGCGCGATTGCTGATCCCCATGTGTATCACTGGCAAACTGTGATGGACGACACCGTCAGTGCGTCCGTCGCGCAGGCTCTCGATGAG
CTGATGCTTTGGGCCGAGGACTGCCCCGAAGTCCGGCACCTCGTGCACGCGGATTTCGGCTCCAACAATGTCCTGACGGACAATGGCCGC
ATAACAGCGGTCATTGACTGGAGCGAGGCGATGTTCGGGGATTCCCAATACGAGGTCGCCAACATCTTCTTCTGGAGGCCGTGGTTGGCT
TGTATGGAGCAGCAGACGCGCTACTTCGAGCGGAGGCATCCGGAGCTTGCAGGATCGCCACGACTCCGGGCGTATATGCTCCGCATTGGT
CTTGACCAACTCTATCAGAGCTTGGTTGACGGCAATTTCGATGATGCAGCTTGGGCGCAGGGTCGATGCGACGCAATCGTCCGATCCGGA
GCCGGGACTGTCGGGCGTACACAAATCGCCCGCAGAAGCGCGGCCGTCTGGACCGATGGCTGTGTAGAAGTACTCGCCGATAGTGGAAAC

CGACGCCCCAGCACTCGTCCGAGGGCAAAGAAATAGAGTAGATGCCGACCGGATCTGTCGATCGACAAGCTCGAGTTTCTCCATAATAAT
GTGTGAGTAGTTCCCAGATAAGGGAATTAGGGTTCCTATAGGGTTTCGCTCATGTGTTGAGCATATAAGAAACCCTTAGTATGTATTTGT
ATTTGTAAAATACTTCTATCAATAAAATTTCTAATTCCTAAAACCAAAATCCAGTACTAAAATCCAGATCCCCCGAATTAATTCGGCGTT
AATTCAGTACATTAAAAACGTCCGCAATGTGTTATTAAGTTGTCTAAGCGTCAATTTGTTTACACCACAATATATCCTGCCA

T-DNA right border
T-DNA left border
*ccd*B gene
*Bsa*I site
Inverted *Bsa*I site
Chloramphenicol resistance gene
attB1
attB2
Nos terminator
CaMV promoter
kanamycin resistance gene
Hygromycin resistance gene
2x35S CaMV promoter
CaMV terminator

**> *pMDC32B-AtmiR173aTS-B/c* (**11635 bp)

CCAGCCAGCCAACAGCTCCCCGACCGGCAGCTCGGCACAAAATCACCACTCGATACAGGCAGCCCATCAGTCCGGGACGGCGTCAGCGGG
AGAGCCGTTGTAAGGCGGCAGACTTTGCTCATGTTACCGATGCTATTCGGAAGAACGGCAACTAAGCTGCCGGGTTTGAAACACGGATGA
TCTCGCGGAGGGTAGCATGTTGATTGTAACGATGACAGAGCGTTGCTGCCTGTGATCACCGCGGTTTCAAAATCGGCTCCGTCGATACTA
TGTTATACGCCAACTTTGAAAACAACTTTGAAAAAGCTGTTTTCTGGTATTTAAGGTTTTAGAATGCAAGGAACAGTGAATTGGAGTTCG
TCTTGTTATAATTAGCTTCTTGGGGTATCTTTAAATACTGTAGAAAAGAGGAAGGAAATAATAAATGGCTAAAATGAGAATATCACCGGA
ATTGAAAAAACTGATCGAAAAATACCGCTGCGTAAAAGATACGGAAGGAATGTCTCCTGCTAAGGTATATAAGCTGGTGGGAGAAAATGA
AAACCTATATTTAAAAATGACGGACAGCCGGTATAAAGGGACCACCTATGATGTGGAACGGGAAAAGGACATGATGCTATGGCTGGAAGG
AAAGCTGCCTGTTCCAAAGGTCCTGCACTTTGAACGGCATGATGGCTGGAGCAATCTGCTCATGAGTGAGGCCGATGGCGTCCTTTGCTC
GGAAGAGTATGAAGATGAACAAAGCCCTGAAAAGATTATCGAGCTGTATGCGGAGTGCATCAGGCTCTTTCACTCCATCGACATATCGGA
TTGTCCCTATACGAATAGCTTAGACAGCCGCTTAGCCGAATTGGATTACTTACTGAATAACGATCTGGCCGATGTGGATTGCGAAAACTG
GGAAGAAGACACTCCATTTAAAGATCCGCGCGAGCTGTATGATTTTTTAAAGACGGAAAAGCCCGAAGAGGAACTTGTCTTTTCCCACGG
CGACCTGGGAGACAGCAACATCTTTGTGAAAGATGGCAAAGTAAGTGGCTTTATTGATCTTGGGAGAAGCGGCAGGGCGGACAAGTGGTA
TGACATTGCCTTCTGCGTCCGGTCGATCAGGGAGGATATCGGGGAAGAACAGTATGTCGAGCTATTTTTTGACTTACTGGGGATCAAGCC
TGATTGGGAGAAAATAAAATATTATATTTTACTGGATGAATTGTTTTAGTACCTAGAATGCATGACCAAAATCCCTTAACGTGAGTTTTC
GTTCCACTGAGCGTCAGACCCCGTAGAAAAGATCAAAGGATCTTCTTGAGATCCTTTTTTTCTGCGCGTAATCTGCTGCTTGCAAACAAA
AAAACCACCGCTACCAGCGGTGGTTTGTTTGCCGGATCAAGAGCTACCAACTCTTTTTCCGAAGGTAACTGGCTTCAGCAGAGCGCAGAT
ACCAAATACTGTCCTTCTAGTGTAGCCGTAGTTAGGCCACCACTTCAAGAACTCTGTAGCACCGCCTACATACCTCGCTCTGCTAATCCT
GTTACCAGTGGCTGCTGCCAGTGGCGATAAGTCGTGTCTTACCGGGTTGGACTCAAGACGATAGTTACCGGATAAGGCGCAGCGGTCGGG
CTGAACGGGGGGTTCGTGCACACAGCCCAGCTTGGAGCGAACGACCTACACCGAACTGAGATACCTACAGCGTGAGCTATGAGAAAGCGC
CACGCTTCCCGAAGGGAGAAAGGCGGACAGGTATCCGGTAAGCGGCAGGGTCGGAACAGGAGAGCGCACGAGGGAGCTTCCA GGGGGAAA
CGCCTGGTATCTTTATAGTCCTGTCGGGTTTCGCCACCTCTGACTTGAGCGTCGATTTTTGTGATGCTCGTCAGGGGGGCGGAGCCTATG
GAAAAACGCCAGCAACGCGGCCTTTTTACGGTTCCTGGCCTTTTGCTGGCCTTTTGCTCACATGTTCTTTCCTGCGTTATCCCCTGATTC
TGTGGATAACCGTATTACCGCCTTTGAGTGAGCTGATACCGCTCGCCGCAGCCGAACGACCGAGCGCAGCGAGTCAGTGAGCGAGGAAGC
GGAAGAGCGCCTGATGCGGTATTTTCTCCTTACGCATCTGTGCGGTATTTCACACCGCATATGGTGCACTCTCAGTACAATCTGCTCTGA
TGCCGCATAGTTAAGCCAGTATACACTCCGCTATCGCTACGTGACTGGGTCATGGCTGCGCCCCGACACCCGCCAACACCCGCTGACGCG
CCCTGACGGGCTTGTCTGCTCCCGGCATCCGCTTACAGACAAGCTGTGACCGTCTCCGGGAGCTGCATGTGTCAGAGGTTTTCACCGTCA
TCACCGAAACGCGCGAGGCAGGGTGCCTTGATGTGGGCGCCGGCGGTCGAGTGGCGACGGCGCGGCTTGTCCGCGCCCTGGTAGATTGCC
TGGCCGTAGGCCAGCCATTTTTGAGCGGCCAGCGGCCGCGATAGGCCGACGCGAAGCGGCGGGGCGTAGGGAGCGCAGCGACCGAAGGGT
AGGCGCTTTTTGCAGCTCTTCGGCTGTGCGCTGGCCAGACAGTTATGCACAGGCCAGGCGGGTTTTAAGAGTTTTAATAAGTTTTAAAGA
GTTTTAGGCGGAAAAATCGCCTTTTTTCTCTTTTATATCAGTCACTTACATGTGTGACCGGTTCCCAATGTACGGCTTTGGGTTCCCAAT
GTACGGGTTCCGGTTCCCAATGTACGGCTTTGGGTTCCCAATGTACGTGCTATCCACAGGAAAGAGAACTTTTCGACCTTTTTCCCCTGC
TAGGGCAATTTGCCCTAGCATCTGCTCCGTACATTAGGAACCGGCGGATGCTTCGCCCTCGATCAGGTTGCGGTAGCGCATGACTAGGAT
CGGGCCAGCCTGCCCCGCCTCCTCCTTCAAATCGTACTCCGGCAGGTCATTTGACCCGATCAGCTTGCGCACGGTGAAACAGAACTTCTT
GAACTCTCCGGCGCTGCCACTGCGTTCGTAGATCGTCTTGAACAACCATCTGGCTTCTGCCTTGCCTGCGGCGCGGCGTGCCAGGCGGTA
GAGAAAACGGCCGATGCCGGGATCGATCAAAAAGTAATCGGGGTGAACCGTCAGCACGTCCGGGTTCTTGCCTTCTGTGATCTCGCGGTA
CATCCAATCAGCTAGCTCGATCTCGATGTACTCCGGCCGCCCGGTTTCGCTCTTTACGATCTTGTAGCGGCTAATCAAGGCTTCACCCTC
GGATACCGTCACCAGGCGGCCGTTCTTGGCCTTCTTCGTACGCTGCATGGCAACGTGCGTGGTGTTTAACCGAATGCAGGTTTCTACCAG
GTCGTCTTTCTGCTTTCCGCCATCGGCTCGCCGGCAGAACTTGAGTACGTCCGCAACGTGTGGACGGAACACGCGGCCGGGCTTGTCTCC
CTTCCCTTCCCGGTATCGGTTCATGGATTCGGTTAGATGGGAAACCGCCATCAGTACCAGGTCGTAATCCCACACACTGGCCATGCCGGC
CGGCCCTGCGGAAACCTCTACGTGCCCGTCTGGAAGCTCGTAGCGGATCACCTCGCCAGCTCGTCGGTCACGCTTCGACAGACGGAAAAC
GGCCACGTCCATGATGCTGCGACTATCGCGGGTGCCCACGTCATAGAGCATCGGAACGAAAAAATCTGGTTGCTCGTCGCCCTTGGGCGG
CTTCCTAATCGACGGCGCACCGGCTGCCGGCGGTTGCCGGGATTCTTTGCGGATTCGATCAGCGGCCGCTTGCCACGATTCACCGGGGCG
TGCTTCTGCCTCGATGCGTTGCCGCTGGGCGGCCTGCGCGGCCTTCAACTTCTCCACCAGGTCATCACCCAGCGCCGCGCCGATTTGTAC
CGGGCCGGATGGTTTGCGACCGTCACGCCGATTCCTCGGGCTTGGGGGTTCCAGTGCCATTGCAGGGCCGGCAGACAACCCAGCCGCTTA
CGCCTGGCCAACCGCCCGTTCCTCCACACATGGGGCATTCCACGGCGTCGGTGCCTGGTTGTTCTTGATTTTCCATGCCGCCTCCTTTAG
CCGCTAAAATTCATCTACTCATTTATTCATTTGCTCATTTACTCTGGTAGCTGCGCGATGTATTCAGATAGCAGCTCGGTAATGGTCTTG
CCTTGGCGTACCGCGTACATCTTCAGCTTGGTGTGATCCTCCGCCGGCAACTGAAAGTTGACCCGCTTCATGGCTGGCGTGTCTGCCAGG
CTGGCCAACGTTGCAGCCTTGCTGCTGCGTGCGCTCGGACGGCCGGCACTTAGCGTGTTTGTGCTTTTGCTCATTTTCTCTTTACCTCAT
TAACTCAAATGAGTTTTGATTTAATTTCAGCGGCCAGCGCCTGGACCTCGCGGGCAGCGTCGCCCTCGGGTTCTGATTCAAGAACGGTTG
TGCCGGCGGCGGCAGTGCCTGGGTAGCTCACGCGCTGCGTGATACGGGACTCAAGAATGGGCAGCTCGTACCCGGCCAGCGCCTCGGCAA
CCTCACCGCCGATGCGCGTGCCTTTGATCGCCCGCGACACGACAAAGGCCGCTTGTAGCCTTCCATCCGTGACCTCAATGCGCTGCTTAA
CCAGCTCCACCAGGTCGGCGGTGGCCCATATGTCGTAAGGGCTTGGCTGCACCGGAATCAGCACGAAGTCGGCTGCCTTGATCGCGGACA
CAGCCAAGTCCGCCGCCTGGGGCGCTCCGTCGATCACTACGAAGTCGCGCCGGCCGATGGCCTTCACGTCGCGGTCAATCGTCGGGCGGT
CGATGCCGACAACGGTTAGCGGTTGATCTTCCCGCACGGCCGCCCAATCGCGGGCACTGCCCTGGGGATCGGAATCGACTAACAGAACAT
CGGCCCCGGCGAGTTGCAGGGCGCGGGCTAGATGGGTTGCGATGGTCGTCTTGCCTGACCCGCCTTTCTGGTTAAGTACAGCGATAACCT
TCATGCGTTCCCCTTGCGTATTTGTTTATTTACTCATCGCATCATATACGCAGCGACCGCATGACGCAAGCTGTTTTACTCAAATACACA
TCACCTTTTTAGACGGCGGCGCTCGGTTTCTTCAGCGGCCAAGCTGGCCGGCCAGGCCGCCAGCTTGGCATCAGACAAAC CGGCCAGGAT
TTCATGCAGCCGCACGGTTGAGACGTGCGCGGGCGGCTCGAACACGTACCCGGCCGCGATCATCTCCGCCTCGATCTCTTCGGTAATGAA
AAACGGTTCGTCCTGGCCGTCCTGGTGCGGTTTCATGCTTGTTCCTCTTGGCGTTCATTCTCGGCGGCCGCCAGGGCGTCGGCCTCGGTC
AATGCGTCCTCACGGAAGGCACCGCGCCGCCTGGCCTCGGTGGGCGTCACTTCCTCGCTGCGCTCAAGTGCGCGGTACAGGGTCGAGCGA
TGCACGCCAAGCAGTGCAGCCGCCTCTTTCACGGTGCGGCCTTCCTGGTCGATCAGCTCGCGGGCGTGCGCGATCTGTGCCGGGGTGAGG
GTAGGGCGGGGGCCAAACTTCACGCCTCGGGCCTTGGCGGCCTCGCGCCCGCTCCGGGTGCGGTCGATGATTAGGGAACGCTCGAACTCG

GCAATGCCGGCGAACACGGTCAACACCATGCGGCCGGCCGGCGTGGTGGTGTCGGCCCACGGCTCTGCCAGGCTACGCAGGCCCGCGCCG
GCCTCCTGGATGCGCTCGGCAATGTCCAGTAGGTCGCGGGTGCTGCGGGCCAGGCGGTCTAGCCTGGTCACTGTCACAACGTCGCCAGGG
CGTAGGTGGTCAAGCATCCTGGCCAGCTCCGGGCGGTCGCGCCTGGTGCCGGTGATCTTCTCGGAAAACAGCTTGGTGCAGCCGGCCGCG
TGCAGTTCGGCCCGTTGGTTGGTCAAGTCCTGGTCGTCGGTGCTGACGCGGGCATAGCCCAGCAGGCCAGCGGCGGCGCTCTTGTTCATG
GCGTAATGTCTCCGGTTCTAGTCGCAAGTATTCTACTTTATGCGACTAAAACACGCGACAAGAAAACGCCAGGAAAAGGGCAGGGCGGCA
GCCTGTCGCGTAACTTAGGACTTGTGCGACATGTCGTTTTCAGAAGACGGCTGCACTGAACGTCAGAAGCCGACTGCACTATAGCAGCGG
AGGGGTTGGATCAAAGTACTTTGATCCCGAGGGGAACCCTGTGGTTGGCATGCACATACAAATGGACGAACGGATAAACCTTTTCACGCC
CTTTTAAATATCCGTTATTCTAATAAACGCTCTTTTCTCTTAGGTTTACCCGCCAATATATCCTGTCAAACACTGATAGTTTAAACTGAA
GGCGGGAAACGACAATCTGATCCAAGCTCAAGCTGCTCTAGCATTCGCCATTCAGGCTGCGCAACTGTTGGGAAGGGCGATCGGTGCGGG
CCTCTTCGCTATTACGCCAGCTGGCGAAAGGGGGATGTGCTGCAAGGCGATTAAGTTGGGTAACGCCAGGGTTTTCCCAGTCACGACGTT
GTAAAACGACGGCCAGTGCCAAGCTTGGCGTGCCTGCAGGTCAACATGGTGGAGCACGACACACTTGTCTACTCCAAAAATATCAAAGAT
ACAGTCTCAGAAGACCAAAGGGCAATTGAGACTTTTCAACAAAGGGTAATATCCGGAAACCTCCTCGGATTCCATTGCCCAGCTATCTGT
CACTTTATTGTGAAGATAGTGGAAAAGGAAGGTGGCTCCTACAAATGCCATCATTGCGATAAAGGAAAGGCCATCGTTGAAGATGCCTCT
GCCGACAGTGGTCCCAAAGATGGACCCCCACCCACGAGGAGCATCGTGGAAAAAGAAGACGTTCCAACCACGTCTTCAAAGCAAGTGGAT
TGATGTGATAACATGGTGGAGCACGACACACTTGTCTACTCCAAAAATATCAAAGATACAGTCTCAGAAGACCAAAGGGCAATTGAGACT
TTTCAACAAAGGGTAATATCCGGAAACCTCCTCGGATTCCATTGCCCAGCTATCTGTCACTTTATTGTGAAGATAGTGGAAAAGGAAGGT
GGCTCCTACAAATGCCATCATTGCGATAAAGGAAAGGCCATCGTTGAAGATGCCTCTGCCGACAGTGGTCCCAAAGATGGACCCCCACCC
ACGAGGAGCATCGTGGAAAAAGAAGACGTTCCAACCACGTCTTCAAAGCAAGTGGATTGATGTGATATCTCCACTGACGTAAGGGATGAC
GCACAATCCCACTATCCTTCGCAAGACCCTTCCTCTATATAAGGAAGTTCATTTCATTTGGAGAGGACCTCGACTCTAGAGGATCCCCGG
GTACCGGGCCCCCCCTCGAGGCGCGCCAAGCTATCAAACAAGTTTGTACAAAAAAGCAGGCTCCGCGGCCGCCCCCTTCACCTGTA**GTGA**
**TTTTTCTCTACAAGCGAATAGACCATTTAA**GAGACCATTAGGCACCCCAGGCTTTACACTTTATGCTTCCGGCTCGTATAATGTGTGGAT
TTTGAGTTAGGAGCCGTCGAGATTTTCAGGAGCTAAGGAAGCTAAAATGGAGAAAAAAATCACTGGATATACCACCGTTGATATATCCCA
ATGGCATCGTAAAGAACATTTTGAGGCATTTCAGTCAGTTGCTCAATGTACCTATAACCAGACCGTTCAGCTGGATATTACGGCCTTTTT
AAAGACCGTAAAGAAAAATAAGCACAAGTTTTATCCGGCCTTTATTCACATTCTTGCCCGCCTGATGAATGCTCATCCGGAGTTCCGTAT
GGCAATGAAAGACGGTGAGCTGGTGATATGGGATAGTGTTCACCCTTGTTACACCGTTTTCCATGAGCAAACTGAAACGTTTTCATCGCT
CTGGAGTGAATACCACGACGATTTCCGGCAGTTTCTACACATATATTCGCAAGATGTGGCGTGTTACGGTGAAAACCTGGCCTATTTCCC
TAAAGGGTTTATTGAGAATATGTTTTTCGTCTCAGCCAATCCCTGGGTGAGTTTCACCAGTTTTGATTTAAACGTGGCCAATATGGACAA
CTTCTTCGCCCCCGTTTTCACCATGGGCAAATATTATACGCAAGGCGACAAGGTGCTGATGCCGCTGGCGATTCAGGTTCATCATGCCGT
TTGTGATGGCTTCCATGTCGGCAGAATGCTTAATGAATTACAACAGTACTGCGATGAGTGGCAGGGCGGGGCGTAAACGCGTGGAGCCGG
CTTACTAAAAGCCAGATAACAGTATGCGTATTTGCGCGCTGATTTTTGCGGTATAAGAATATATACTGATATGTATACCCGAAGTATGTC
AAAAAGAGGTATGCTATGAAGCAGCGTATTACAGTGACAGTTGACAGCGACAGCTATCAGTTGCTCAAGGCATATATGATGTCAATATCT
CCGGTCTGGTAAGCACAACCATGCAGAATGAAGCCCGTCGTCTGCGTGCCGAACGCTGGAAAGCGGAAAATCAGGAAGGGATGGCTGAGG
TCGCCCGGTTTATTGAAATGAACGGCTCTTTTGCTGACGAGAACAGGGGCTGGTGAAATGCAGTTTAAGGTTTACACCTATAAAAGAGAG
AGCCGTTATCGTCTGTTTGTGGATGTACAGAGTGATATTATTGACACGCCCGGCCGACGGATGGTGATCCCCCTGGCCAGTGCACGTCTG
CTGTCAGATAAAGTCTCCCGTGAACTTTACCCGGTGGTGCATATCGGGGATGAAAGCTGGCGCATGATGACCACCGATATGGCCAGTGTG
CCGGTTTCCGTTATCGGGGAAGAAGTGGCTGATCTCAGCCACCGCGAAAATGACATCAAAAACGCCATTAACCTGATGTTCTGGGGAATA
TAAATGTCAGGCTCCCTTATACACAGCCAGTCTGCACCTCGACGGTCTCACATTAAGGGTGGGCGCGCCGACCCAGCTTTCTTGTACAAA
GTGGTTCGATAATTCCTTAATTAACTAGTTCTAGAGCGGCCGCCCACCGCGGTGGAGCTCGAATTTCCCCGATCGTTCAAACATTTGGCA
ATAAAGTTTCTTAAGATTGAATCCTGTTGCCGGTCTTGCGATGATTATCATATAATTTCTGTTGAATTACGTTAAGCATGTAATAATTAA
CATGTAATGCATGACGTTATTTATGAGATGGGTTTTTATGATTAGAGTCCCGCAATTATACATTTAATACGCGATAGAAAACAAAATATA
GCGCGCAAACTAGGATAAATTATCGCGCGCGGTGTCATCTATGTTACTGAATTCGTAATCATGGTCATAGCTGTTTCCTGTGTGAAATTG
TTATCCGCTCACAATTCCACACAACATACGAGCCGGAAGCATAAAGTGTAAAGCCTGGGGTGCCTAATGAGTGAGCTAACTCACATTAAT
TGCGTTGCGCTCACTGCCCGCTTTCCAGTCGGGAAACCTGTCGTGCCAGCTGCATTAATGAATCGGCCAACGCGCGGGGAGAGGCGGTTT
GCGTATTGGCTAGAGCAGCTTGCCAACATGGTGGAGCACGACACTCTCGTCTACTCCAAGAATATCAAAGATACAGTCTCAGAAGACCAA
AGGGCTATTGAGACTTTTCAACAAAGGGTAATATCGGGAAACCTCCTCGGATTCCATTGCCCAGCTATCTGTCACTTCATCAAAAGGACA
GTAGAAAAGGAAGGTGGCACCTACAAATGCCATCATTGCGATAAAGGAAAGGCTATCGTTCAAGATGCCTCTGCCGACAGTGGTCCCAAA
GATGGACCCCCACCCACGAGGAGCATCGTGGAAAAAGAAGACGTTCCAACCACGTCTTCAAAGCAAGTGGATTGATGTGATAACATGGTG
GAGCACGACACTCTCGTCTACTCCAAGAATATCAAAGATACAGTCTCAGAAGACCAAAGGGCTATTGAGACTTTTCAACAAAGGGTAATA
TCGGGAAACCTCCTCGGATTCCATTGCCCAGCTATCTGTCACTTCATCAAAAGGACAGTAGAAAAGGAAGGTGGCACCTACAAATGCCAT
CATTGCGATAAAGGAAAGGCTATCGTTCAAGATGCCTCTGCCGACAGTGGTCCCAAAGATGGACCCCCACCCACGAGGAGCATCGTGGAA
AAAGAAGACGTTCCAACCACGTCTTCAAAGCAAGTGGATTGATGTGATATCTCCACTGACGTAAGGGATGACGCACAATCCCACTATCCT
TCGCAAGACCTTCCTCTATATAAGGAAGTTCATTTCATTTGGAGAGGACACGCTGAAATCACCAGTCTCTCTCTACAAATCTATCTCTCT
CGAGCTTTCGCAGATCCCGGGGGGCAATGAGATATGAAAAAGCCTGAACTCACCGCGACGTCTGTCGAGAAGTTTCTGATCGAAAAGTTC
GACAGCGTCTCCGACCTGATGCAGCTCTCGGAGGGCGAAGAATCTCGTGCTTTCAGCTTCGATGTAGGAGGGCGTGGATATGTCCTGCGG
GTAAATAGCTGCGCCGATGGTTTCTACAAAGATCGTTATGTTTATCGGCACTTTGCATCGGCCGCGCTCCCGATTCCGGAAGTGCTTGAC
ATTGGGGAGTTTAGCGAGAGCCTGACCTATTGCATCTCCCGCCGTGCACAGGGTGTCACGTTGCAAGACCTGCCTGAAACCGAACTGCCC
GCTGTTCTACAACCGGTCGCGGAGGCTATGGATGCGATCGCTGCGGCCGATCTTAGCCAGACGAGCGGGTTCGGCCCATTCGGACCGCAA
GGAATCGGTCAATACACTACATGGCGTGATTTCATATGCGCGATTGCTGATCCCCATGTGTATCACTGGCAAACTGTGATGGACGACACC
GTCAGTGCGTCCGTCGCGCAGGCTCTCGATGAGCTGATGCTTTGGGCCGAGGACTGCCCCGAAGTCCGGCACCTCGTGCACGCGGATTTC
GGCTCCAACAATGTCCTGACGGACAATGGCCGCATAACAGCGGTCATTGACTGGAGCGAGGCGATGTTCGGGGATTCCCAATACGAGGTC
GCCAACATCTTCTTCTGGAGGCCGTGGTTGGCTTGTATGGAGCAGCAGACGCGCTACTTCGAGCGGAGGCATCCGGAGCTTGCAGGATCG
CCACGACTCCGGGCGTATATGCTCCGCATTGGTCTTGACCAACTCTATCAGAGCTTGGTTGACGGCAATTTCGATGATGCAGCTTGGGCG
CAGGGTCGATGCGACGCAATCGTCCGATCCGGAGCCGGGACTGTCGGGCGTACACAAATCGCCCGCAGAAGCGCGGCCGTCTGGACCGAT

GGCTGTGTAGAAGTACTCGCCGATAGTGGAAACCGACGCCCCAGCACTCGTCCGAGGGCAAAGAAATAGAGTAGATGCCGACCGGATCTG
TCGATCGACAAGCTCGAGTTTCTCCATAATAATGTGTGAGTAGTTCCCAGATAAGGGAATTAGGGTTCCTATAGGGTTTCGCTCATGTGT
TGAGCATATAAGAAACCCTTAGTATGTATTTGTATTTGTAAAATACTTCTATCAATAAAATTTCTAATTCCTAAAACCAAAATCCAGTAC
TAAAATCCAGATCCCCCGAATTAATTCGGCGTTAATTCAGTACATTAAAAACGTCCGCAATGTGTTATTAAGTTGTCTAAGCGTCAATTT
GTTTACACCACAATATATCCTGCCA

**AtmiR173a target site**
**AtTAS1c-derived spacer**
T-DNA right border
T-DNA left border
*ccd*B gene
*Bsa*I site
Inverted *Bsa*I site
Chloramphenicol resistance gene
attB1
attB2
Nos terminator
CaMV promoter
kanamycin resistance gene
Hygromycin resistance gene
2x35S CaMV promoter
CaMV terminator

**>*pMDC32B-NbmiR482aTS-B/c*** (11635 bp)

CCAGCCAGCCAACAGCTCCCCGACCGGCAGCTCGGCACAAAATCACCACTCGATACAGGCAGCCCATCAGTCCGGGACGGCGTCAGCGGG
AGAGCCGTTGTAAGGCGGCAGACTTTGCTCATGTTACCGATGCTATTCGGAAGAACGGCAACTAAGCTGCCGGGTTTGAAACACGGATGA
TCTCGCGGAGGGTAGCATGTTGATTGTAACGATGACAGAGCGTTGCTGCCTGTGATCACCGCGGTTTCAAAATCGGCTCCGTCGATACTA
TGTTATACGCCAACTTTGAAAACAACTTTGAAAAAGCTGTTTTCTGGTATTTAAGGTTTTAGAATGCAAGGAACAGTGAATTGGAGTTCG
TCTTGTTATAATTAGCTTCTTGGGGTATCTTTAAATACTGTAGAAAAGAGGAAGGAAATAATAAATGGCTAAAATGAGAATATCACCGGA
ATTGAAAAAACTGATCGAAAAATACCGCTGCGTAAAAGATACGGAAGGAATGTCTCCTGCTAAGGTATATAAGCTGGTGGGAGAAAATGA
AAACCTATATTTAAAAATGACGGACAGCCGGTATAAAGGGACCACCTATGATGTGGAACGGGAAAAGGACATGATGCTATGGCTGGAAGG
AAAGCTGCCTGTTCCAAAGGTCCTGCACTTTGAACGGCATGATGGCTGGAGCAATCTGCTCATGAGTGAGGCCGATGGCGTCCTTTGCTC
GGAAGAGTATGAAGATGAACAAAGCCCTGAAAAGATTATCGAGCTGTATGCGGAGTGCATCAGGCTCTTTCACTCCATCGACATATCGGA
TTGTCCCTATACGAATAGCTTAGACAGCCGCTTAGCCGAATTGGATTACTTACTGAATAACGATCTGGCCGATGTGGATTGCGAAAACTG
GGAAGAAGACACTCCATTTAAAGATCCGCGCGAGCTGTATGATTTTTTAAAGACGGAAAAGCCCGAAGAGGAACTTGTCTTTTCCCACGG
CGACCTGGGAGACAGCAACATCTTTGTGAAAGATGGCAAAGTAAGTGGCTTTATTGATCTTGGGAGAAGCGGCAGGGCGGACAAGTGGTA
TGACATTGCCTTCTGCGTCCGGTCGATCAGGGAGGATATCGGGGAAGAACAGTATGTCGAGCTATTTTTTGACTTACTGGGGATCAAGCC
TGATTGGGAGAAAATAAAATATTATATTTTACTGGATGAATTGTTTTAGTACCTAGAATGCATGACCAAAATCCCTTAACGTGAGTTTTC
GTTCCACTGAGCGTCAGACCCCGTAGAAAAGATCAAAGGATCTTCTTGAGATCCTTTTTTTCTGCGCGTAATCTGCTGCTTGCAAACAAA
AAAACCACCGCTACCAGCGGTGGTTTGTTTGCCGGATCAAGAGCTACCAACTCTTTTTCCGAAGGTAACTGGCTTCAGCAGAGCGCAGAT
ACCAAATACTGTCCTTCTAGTGTAGCCGTAGTTAGGCCACCACTTCAAGAACTCTGTAGCACCGCCTACATACCTCGCTCTGCTAATCCT
GTTACCAGTGGCTGCTGCCAGTGGCGATAAGTCGTGTCTTACCGGGTTGGACTCAAGACGATAGTTACCGGATAAGGCGCAGCGGTCGGG
CTGAACGGGGGGTTCGTGCACACAGCCCAGCTTGGAGCGAACGACCTACACCGAACTGAGATACCTACAGCGTGAGCTATGAGAAAGCGC
CACGCTTCCCGAAGGGAGAAAGGCGGACAGGTATCCGGTAAGCGGCAGGGTCGGAACAGGAGAGCGCACGAGGGAGCTTCCA GGGGGAAA
CGCCTGGTATCTTTATAGTCCTGTCGGGTTTCGCCACCTCTGACTTGAGCGTCGATTTTTGTGATGCTCGTCAGGGGGGCGGAGCCTATG
GAAAAACGCCAGCAACGCGGCCTTTTTACGGTTCCTGGCCTTTTGCTGGCCTTTTGCTCACATGTTCTTTCCTGCGTTATCCCCTGATTC
TGTGGATAACCGTATTACCGCCTTTGAGTGAGCTGATACCGCTCGCCGCAGCCGAACGACCGAGCGCAGCGAGTCAGTGAGCGAGGAAGC
GGAAGAGCGCCTGATGCGGTATTTTCTCCTTACGCATCTGTGCGGTATTTCACACCGCATATGGTGCACTCTCAGTACAATCTGCTCTGA
TGCCGCATAGTTAAGCCAGTATACACTCCGCTATCGCTACGTGACTGGGTCATGGCTGCGCCCCGACACCCGCCAACACCCGCTGACGCG
CCCTGACGGGCTTGTCTGCTCCCGGCATCCGCTTACAGACAAGCTGTGACCGTCTCCGGGAGCTGCATGTGTCAGAGGTTTTCACCGTCA
TCACCGAAACGCGCGAGGCAGGGTGCCTTGATGTGGGCGCCGGCGGTCGAGTGGCGACGGCGCGGCTTGTCCGCGCCCTGGTAGATTGCC
TGGCCGTAGGCCAGCCATTTTTGAGCGGCCAGCGGCCGCGATAGGCCGACGCGAAGCGGCGGGGCGTAGGGAGCGCAGCGACCGAAGGGT
AGGCGCTTTTTGCAGCTCTTCGGCTGTGCGCTGGCCAGACAGTTATGCACAGGCCAGGCGGGTTTTAAGAGTTTTAATAAGTTTTAAAGA
GTTTTAGGCGGAAAAATCGCCTTTTTTCTCTTTTATATCAGTCACTTACATGTGTGACCGGTTCCCAATGTACGGCTTTGGGTTCCCAAT
GTACGGGTTCCGGTTCCCAATGTACGGCTTTGGGTTCCCAATGTACGTGCTATCCACAGGAAAGAGAACTTTTCGACCTTTTTCCCCTGC
TAGGGCAATTTGCCCTAGCATCTGCTCCGTACATTAGGAACCGGCGGATGCTTCGCCCTCGATCAGGTTGCGGTAGCGCATGACTAGGAT
CGGGCCAGCCTGCCCCGCCTCCTCCTTCAAATCGTACTCCGGCAGGTCATTTGACCCGATCAGCTTGCGCACGGTGAAACAGAACTTCTT
GAACTCTCCGGCGCTGCCACTGCGTTCGTAGATCGTCTTGAACAACCATCTGGCTTCTGCCTTGCCTGCGGCGCGGCGTGCCAGGCGGTA
GAGAAAACGGCCGATGCCGGGATCGATCAAAAAGTAATCGGGGTGAACCGTCAGCACGTCCGGGTTCTTGCCTTCTGTGATCTCGCGGTA
CATCCAATCAGCTAGCTCGATCTCGATGTACTCCGGCCGCCCGGTTTCGCTCTTTACGATCTTGTAGCGGCTAATCAAGGCTTCACCCTC
GGATACCGTCACCAGGCGGCCGTTCTTGGCCTTCTTCGTACGCTGCATGGCAACGTGCGTGGTGTTTAACCGAATGCAGGTTTCTACCAG
GTCGTCTTTCTGCTTTCCGCCATCGGCTCGCCGGCAGAACTTGAGTACGTCCGCAACGTGTGGACGGAACACGCGGCCGGGCTTGTCTCC
CTTCCCTTCCCGGTATCGGTTCATGGATTCGGTTAGATGGGAAACCGCCATCAGTACCAGGTCGTAATCCCACACACTGGCCATGCCGGC
CGGCCCTGCGGAAACCTCTACGTGCCCGTCTGGAAGCTCGTAGCGGATCACCTCGCCAGCTCGTCGGTCACGCT TCGACAGACGGAAAAC
GGCCACGTCCATGATGCTGCGACTATCGCGGGTGCCCACGTCATAGAGCATCGGAACGAAAAAATCTGGTTGCTCGTCGCCCTTGGGCGG
CTTCCTAATCGACGGCGCACCGGCTGCCGGCGGTTGCCGGGATTCTTTGCGGATTCGATCAGCGGCCGCTTGCCACGATTCACCGGGGCG
TGCTTCTGCCTCGATGCGTTGCCGCTGGGCGGCCTGCGCGGCCTTCAACTTCTCCACCAGGTCATCACCCAGCGCCGCGCCGATTTGTAC
CGGGCCGGATGGTTTGCGACCGTCACGCCGATTCCTCGGGCTTGGGGGTTCCAGTGCCATTGCAGGGCCGGCAGACAACCCAGCCGCTTA
CGCCTGGCCAACCGCCCGTTCCTCCACACATGGGGCATTCCACGGCGTCGGTGCCTGGTTGTTCTTGATTTTCCATGCCGCCTCCTTTAG
CCGCTAAAATTCATCTACTCATTTATTCATTTGCTCATTTACTCTGGTAGCTGCGCGATGTATTCAGATAGCAGCTCGGTAATGGTCTTG
CCTTGGCGTACCGCGTACATCTTCAGCTTGGTGTGATCCTCCGCCGGCAACTGAAAGTTGACCCGCTTCATGGCTGGCGTGTCTGCCAGG
CTGGCCAACGTTGCAGCCTTGCTGCTGCGTGCGCTCGGACGGCCGGCACTTAGCGTGTTTGTGCTTTTGCTCATTTTCTCTTTACCTCAT
TAACTCAAATGAGTTTTGATTTAATTTCAGCGGCCAGCGCCTGGACCTCGCGGGCAGCGTCGCCCTCGGGTTCTGATTCAAGAACGGTTG
TGCCGGCGGCGGCAGTGCCTGGGTAGCTCACGCGCTGCGTGATACGGGACTCAAGAATGGGCAGCTCGTACCCGGCCAGCGCCTCGGCAA
CCTCACCGCCGATGCGCGTGCCTTTGATCGCCCGCGACACGACAAAGGCCGCTTGTAGCCTTCCATCCGTGACCTCAATGCGCTGCTTAA
CCAGCTCCACCAGGTCGGCGGTGGCCCATATGTCGTAAGGGCTTGGCTGCACCGGAATCAGCACGAAGTCGGCTGCCTTGATCGCGGACA
CAGCCAAGTCCGCCGCCTGGGGCGCTCCGTCGATCACTACGAAGTCGCGCCGGCCGATGGCCTTCACGTCGCGGTCAATCGTCGGGCGGT
CGATGCCGACAACGGTTAGCGGTTGATCTTCCCGCACGGCCGCCCAATCGCGGGCACTGCCCTGGGGATCGGAATCGACTAACAGAACAT
CGGCCCCGGCGAGTTGCAGGGCGCGGGCTAGATGGGTTGCGATGGTCGTCTTGCCTGACCCGCCTTTCTGGTTAAGTACAGCGATAACCT
TCATGCGTTCCCCTTGCGTATTTGTTTATTTACTCATCGCATCATATACGCAGCGACCGCATGACGCAAGCTGTTTTACTCAAATACACA
TCACCTTTTTAGACGGCGGCGCTCGGTTTCTTCAGCGGCCAAGCTGGCCGGCCAGGCCGCCAGCTTGGCATCAGACAAAC CGGCCAGGAT
TTCATGCAGCCGCACGGTTGAGACGTGCGCGGGCGGCTCGAACACGTACCCGGCCGCGATCATCTCCGCCTCGATCTCTTCGGTAATGAA
AAACGGTTCGTCCTGGCCGTCCTGGTGCGGTTTCATGCTTGTTCCTCTTGGCGTTCATTCTCGGCGGCCGCCAGGGCGTCGGCCTCGGTC
AATGCGTCCTCACGGAAGGCACCGCGCCGCCTGGCCTCGGTGGGCGTCACTTCCTCGCTGCGCTCAAGTGCGCGGTACAGGGTCGAGCGA
TGCACGCCAAGCAGTGCAGCCGCCTCTTTCACGGTGCGGCCTTCCTGGTCGATCAGCTCGCGGGCGTGCGCGATCTGTGCCGGGGTGAGG
GTAGGGCGGGGGCCAAACTTCACGCCTCGGGCCTTGGCGGCCTCGCGCCCGCTCCGGGTGCGGTCGATGATTAGGGAACGCTCGAACTCG

GCAATGCCGGCGAACACGGTCAACACCATGCGGCCGGCCGGCGTGGTGGTGTCGGCCCACGGCTCTGCCAGGCTACGCAGGCCCGCGCCG
GCCTCCTGGATGCGCTCGGCAATGTCCAGTAGGTCGCGGGTGCTGCGGGCCAGGCGGTCTAGCCTGGTCACTGTCACAACGTCGCCAGGG
CGTAGGTGGTCAAGCATCCTGGCCAGCTCCGGGCGGTCGCGCCTGGTGCCGGTGATCTTCTCGGAAAACAGCTTGGTGCAGCCGGCCGCG
TGCAGTTCGGCCCGTTGGTTGGTCAAGTCCTGGTCGTCGGTGCTGACGCGGGCATAGCCCAGCAGGCCAGCGGCGGCGCTCTTGTTCATG
GCGTAATGTCTCCGGTTCTAGTCGCAAGTATTCTACTTTATGCGACTAAAACACGCGACAAGAAAACGCCAGGAAAAGGGCAGGGCGGCA
GCCTGTCGCGTAACTTAGGACTTGTGCGACATGTCGTTTTCAGAAGACGGCTGCACTGAACGTCAGAAGCCGACTGCACTATAGCAGCGG
AGGGGTTGGATCAAAGTACTTTGATCCCGAGGGGAACCCTGTGGTTGGCATGCACATACAAATGGACGAACGGATAAACCTTTTCACGCC
CTTTTAAATATCCGTTATTCTAATAAACGCTCTTTTCTCTTAGGTTTACCCGCCAATATATCCTGTCAAACACTGATAGTTTAAACTGAA
GGCGGGAAACGACAATCTGATCCAAGCTCAAGCTGCTCTAGCATTCGCCATTCAGGCTGCGCAACTGTTGGGAAGGGCGATCGGTGCGGG
CCTCTTCGCTATTACGCCAGCTGGCGAAAGGGGGATGTGCTGCAAGGCGATTAAGTTGGGTAACGCCAGGGTTTTCCCAGTCACGACGTT
GTAAAACGACGGCCAGTGCCAAGCTTGGCGTGCCTGCAGGTCAACATGGTGGAGCACGACACACTTGTCTACTCCAAAAATATCAAAGAT
ACAGTCTCAGAAGACCAAAGGGCAATTGAGACTTTTCAACAAAGGGTAATATCCGGAAACCTCCTCGGATTCCATTGCCCAGCTATCTGT
CACTTTATTGTGAAGATAGTGGAAAAGGAAGGTGGCTCCTACAAATGCCATCATTGCGATAAAGGAAAGGCCATCGTTGAAGATGCCTCT
GCCGACAGTGGTCCCAAAGATGGACCCCCACCCACGAGGAGCATCGTGGAAAAAGAAGACGTTCCAACCACGTCTTCAAAGCAAGTGGAT
TGATGTGATAACATGGTGGAGCACGACACACTTGTCTACTCCAAAAATATCAAAGATACAGTCTCAGAAGACCAAAGGGCAATTGAGACT
TTTCAACAAAGGGTAATATCCGGAAACCTCCTCGGATTCCATTGCCCAGCTATCTGTCACTTTATTGTGAAGATAGTGGAAAAGGAAGGT
GGCTCCTACAAATGCCATCATTGCGATAAAGGAAAGGCCATCGTTGAAGATGCCTCTGCCGACAGTGGTCCCAAAGATGGACCCCCACCC
ACGAGGAGCATCGTGGAAAAAGAAGACGTTCCAACCACGTCTTCAAAGCAAGTGGATTGATGTGATATCTCCACTGACGTAAGGGATGAC
GCACAATCCCACTATCCTTCGCAAGACCCTTCCTCTATATAAGGAAGTTCATTTCATTTGGAGAGGACCTCGACTCTAGAGGATCCCCGG
GTACCGGGCCCCCCCTCGAGGCGCGCCAAGCTATCAAACAAGTTTGTACAAAAAAGCAGGCTCCGCGGCCGCCCCCTTCACCTGTA**GTGG**
**TATGGGGGGAGTCGGGAATAGACCATTTA**AGAGACCATTAGGCACCCCAGGCTTTACACTTTATGCTTCCGGCTCGTATAATGTGTGGAT
TTTGAGTTAGGAGCCGTCGAGATTTTCAGGAGCTAAGGAAGCTAAAATGGAGAAAAAAATCACTGGATATACCACCGTTGATATATCCCA
ATGGCATCGTAAAGAACATTTTGAGGCATTTCAGTCAGTTGCTCAATGTACCTATAACCAGACCGTTCAGCTGGATATTACGGCCTTTTT
AAAGACCGTAAAGAAAAATAAGCACAAGTTTTATCCGGCCTTTATTCACATTCTTGCCCGCCTGATGAATGCTCATCCGGAGTTCCGTAT
GGCAATGAAAGACGGTGAGCTGGTGATATGGGATAGTGTTCACCCTTGTTACACCGTTTTCCATGAGCAAACTGAAACGTTTTCATCGCT
CTGGAGTGAATACCACGACGATTTCCGGCAGTTTCTACACATATATTCGCAAGATGTGGCGTGTTACGGTGAAAACCTGGCCTATTTCCC
TAAAGGGTTTATTGAGAATATGTTTTTCGTCTCAGCCAATCCCTGGGTGAGTTTCACCAGTTTTGATTTAAACGTGGCCAATATGGACAA
CTTCTTCGCCCCCGTTTTCACCATGGGCAAATATTATACGCAAGGCGACAAGGTGCTGATGCCGCTGGCGATTCAGGTTCATCATGCCGT
TTGTGATGGCTTCCATGTCGGCAGAATGCTTAATGAATTACAACAGTACTGCGATGAGTGGCAGGGCGGGGCGTAAACGCGTGGAGCCGG
CTTACTAAAAGCCAGATAACAGTATGCGTATTTGCGCGCTGATTTTTGCGGTATAAGAATATATACTGATATGTATACCCGAAGTATGTC
AAAAAGAGGTATGCTATGAAGCAGCGTATTACAGTGACAGTTGACAGCGACAGCTATCAGTTGCTCAAGGCATATATGATGTCAATATCT
CCGGTCTGGTAAGCACAACCATGCAGAATGAAGCCCGTCGTCTGCGTGCCGAACGCTGGAAAGCGGAAAATCAGGAAGGGATGGCTGAGG
TCGCCCGGTTTATTGAAATGAACGGCTCTTTTGCTGACGAGAACAGGGGCTGGTGAAATGCAGTTTAAGGTTTACACCTATAAAAGAGAG
AGCCGTTATCGTCTGTTTGTGGATGTACAGAGTGATATTATTGACACGCCCGGCCGACGGATGGTGATCCCCCTGGCCAGTGCACGTCTG
CTGTCAGATAAAGTCTCCCGTGAACTTTACCCGGTGGTGCATATCGGGGATGAAAGCTGGCGCATGATGACCACCGATATGGCCAGTGTG
CCGGTTTCCGTTATCGGGGAAGAAGTGGCTGATCTCAGCCACCGCGAAAATGACATCAAAAACGCCATTAACCTGATGTTCTGGGGAATA
TAAATGTCAGGCTCCCTTATACACAGCCAGTCTGCACCTCGACGGTCTCACATTAAGGGTGGGCGCGCCGACCCAGCTTTCTTGTACAAA
GTGGTTCGATAATTCCTTAATTAACTAGTTCTAGAGCGGCCGCCCACCGCGGTGGAGCTCGAATTTCCCCGATCGTTCAAACATTTGGCA
ATAAAGTTTCTTAAGATTGAATCCTGTTGCCGGTCTTGCGATGATTATCATATAATTTCTGTTGAATTACGTTAAGCATGTAATAATTAA
CATGTAATGCATGACGTTATTTATGAGATGGGTTTTTATGATTAGAGTCCCGCAATTATACATTTAATACGCGATAGAAAACAAAATATA
GCGCGCAAACTAGGATAAATTATCGCGCGCGGTGTCATCTATGTTACTGAATTCGTAATCATGGTCATAGCTGTTTCCTGTGTGAAATTG
TTATCCGCTCACAATTCCACACAACATACGAGCCGGAAGCATAAAGTGTAAAGCCTGGGGTGCCTAATGAGTGAGCTAACTCACATTAAT
TGCGTTGCGCTCACTGCCCGCTTTCCAGTCGGGAAACCTGTCGTGCCAGCTGCATTAATGAATCGGCCAACGCGCGGGGAGAGGCGGTTT
GCGTATTGGCTAGAGCAGCTTGCCAACATGGTGGAGCACGACACTCTCGTCTACTCCAAGAATATCAAAGATACAGTCTCAGAAGACCAA
AGGGCTATTGAGACTTTTCAACAAAGGGTAATATCGGGAAACCTCCTCGGATTCCATTGCCCAGCTATCTGTCACTTCATCAAAAGGACA
GTAGAAAAGGAAGGTGGCACCTACAAATGCCATCATTGCGATAAAGGAAAGGCTATCGTTCAAGATGCCTCTGCCGACAGTGGTCCCAAA
GATGGACCCCCACCCACGAGGAGCATCGTGGAAAAAGAAGACGTTCCAACCACGTCTTCAAAGCAAGTGGATTGATGTGATAACATGGTG
GAGCACGACACTCTCGTCTACTCCAAGAATATCAAAGATACAGTCTCAGAAGACCAAAGGGCTATTGAGACTTTTCAACAAAGGGTAATA
TCGGGAAACCTCCTCGGATTCCATTGCCCAGCTATCTGTCACTTCATCAAAAGGACAGTAGAAAAGGAAGGTGGCACCTACAAATGCCAT
CATTGCGATAAAGGAAAGGCTATCGTTCAAGATGCCTCTGCCGACAGTGGTCCCAAAGATGGACCCCCACCCACGAGGAGCATCGTGGAA
AAAGAAGACGTTCCAACCACGTCTTCAAAGCAAGTGGATTGATGTGATATCTCCACTGACGTAAGGGATGACGCACAATCCCACTATCCT
TCGCAAGACCTTCCTCTATATAAGGAAGTTCATTTCATTTGGAGAGGACACGCTGAAATCACCAGTCTCTCTCTACAAATCTATCTCTCT
CGAGCTTTCGCAGATCCCGGGGGGCAATGAGATATGAAAAAGCCTGAACTCACCGCGACGTCTGTCGAGAAGTTTCTGATCGAAAAGTTC
GACAGCGTCTCCGACCTGATGCAGCTCTCGGAGGGCGAAGAATCTCGTGCTTTCAGCTTCGATGTAGGAGGGCGTGGATATGTCCTGCGG
GTAAATAGCTGCGCCGATGGTTTCTACAAAGATCGTTATGTTTATCGGCACTTTGCATCGGCCGCGCTCCCGATTCCGGAAGTGCTTGAC
ATTGGGGAGTTTAGCGAGAGCCTGACCTATTGCATCTCCCGCCGTGCACAGGGTGTCACGTTGCAAGACCTGCCTGAAACCGAACTGCCC
GCTGTTCTACAACCGGTCGCGGAGGCTATGGATGCGATCGCTGCGGCCGATCTTAGCCAGACGAGCGGGTTCGGCCCATTCGGACCGCAA
GGAATCGGTCAATACACTACATGGCGTGATTTCATATGCGCGATTGCTGATCCCCATGTGTATCACTGGCAAACTGTGATGGACGACACC
GTCAGTGCGTCCGTCGCGCAGGCTCTCGATGAGCTGATGCTTTGGGCCGAGGACTGCCCCGAAGTCCGGCACCTCGTGCACGCGGATTTC
GGCTCCAACAATGTCCTGACGGACAATGGCCGCATAACAGCGGTCATTGACTGGAGCGAGGCGATGTTCGGGGATTCCCAATACGAGGTC
GCCAACATCTTCTTCTGGAGGCCGTGGTTGGCTTGTATGGAGCAGCAGACGCGCTACTTCGAGCGGAGGCATCCGGAGCTTGCAGGATCG
CCACGACTCCGGGCGTATATGCTCCGCATTGGTCTTGACCAACTCTATCAGAGCTTGGTTGACGGCAATTTCGATGATGCAGCTTGGGCG
CAGGGTCGATGCGACGCAATCGTCCGATCCGGAGCCGGGACTGTCGGGCGTACACAAATCGCCCGCAGAAGCGCGGCCGTCTGGACCGAT

GGCTGTGTAGAAGTACTCGCCGATAGTGGAAACCGACGCCCCAGCACTCGTCCGAGGGCAAAGAAATAGAGTAGATGCCGACCGGATCTG
TCGATCGACAAGCTCGAGTTTCTCCATAATAATGTGTGAGTAGTTCCCAGATAAGGGAATTAGGGTTCCTATAGGGTTTCGCTCATGTGT
TGAGCATATAAGAAACCCTTAGTATGTATTTGTATTTGTAAAATACTTCTATCAATAAAATTTCTAATTCCTAAAACCAAAATCCAGTAC
TAAAATCCAGATCCCCCGAATTAATTCGGCGTTAATTCAGTACATTAAAAACGTCCGCAATGTGTTATTAAGTTGTCTAAGCGTCAATTT
GTTTACACCACAATATATCCTGCCA

**NbmiR482a target site**
**AtTAS1c-derived spacer**
T-DNA right border
T-DNA left border
*ccd*B gene
*Bsa*I site
Inverted *Bsa*I site
Chloramphenicol resistance gene
attB1
attB2
Nos terminator
CaMV promoter
kanamycin resistance gene
Hygromycin resistance gene
2x35S CaMV promoter
CaMV terminator
